# Alzheimer’s Disease Selectively Perturbs Age-Sensitive Brain Radiomic Features Across the Disease Continuum

**DOI:** 10.64898/2026.06.17.732875

**Authors:** Megha Suman Sharma, Rajat Agarwal, Nikita Tiwari, Megha Sharma, Ajay Kaushik, the Alzheimer’s Disease Neuroimaging Initiative

## Abstract

Normal brain aging and Alzheimer’s disease both involve progressive structural brain alterations, making it challenging to distinguish pathological neurodegeneration from normative aging-related atrophy. This study investigated whether Alzheimer’s disease exhibits radiomic patterns that mimic, diverge from, or selectively perturb age-associated structural brain changes. T1-weighted magnetic resonance imaging scans from the Alzheimer’s Disease Neuroimaging Initiative were analyzed using a region-wise radiomics framework across 10 anatomically defined brain regions. Radiomic features were extracted following automated segmentation, bias field correction, and intensity normalization. Age-associated radiomic patterns were first identified in cognitively normal subjects using Spearman correlation analysis. Features demonstrating significant age sensitivity were subsequently compared between cognitively normal and Alzheimer’s disease cohorts across age bins using Welch’s two-sample t-tests with permutation-based significance estimation and false discovery rate correction. Medial temporal and limbic regions, particularly the hippocampus, entorhinal cortex, and cingulum, demonstrated consistent age-aligned radiomic trajectories with systematic, statistically significant disease-related shifts across all age bins, supported by large effect sizes and bootstrap-validated confidence intervals. In contrast, several other regions demonstrated more heterogeneous and less stable patterns of group separation across age bins. Secondary analysis using late mild cognitive impairment subjects demonstrated that these radiomic divergences are detectable at the transition from normal cognition to mild cognitive impairment, with statistically significant CN–LMCI separation but no significant LMCI–AD separation, positioning the identified markers as early-stage rather than late-stage indicators of neurodegeneration. These findings indicate that Alzheimer’s disease does not uniformly mimic normal aging across the brain but instead selectively perturbs radiomic features associated with normative aging trajectories. The identified markers represent promising candidates for age-adjusted radiomic biomarkers, warranting validation in independent cohorts to establish their generalisability. The fully automated nature of the analytical pipeline — spanning segmentation, feature extraction, and statistical comparison without manual annotation — may facilitate scalable validation of these biomarkers in larger neuroimaging cohorts.

## 1 Introduction

Normal brain aging is associated with gradual structural and microstructural changes, including cortical thinning, tissue loss, and region-specific atrophy [13], [21]. Many neurodegenerative disorders also manifest through progressive brain volume loss, making it difficult to determine whether an observed imaging pattern reflects normative aging or disease-related degeneration [18]. This overlap is especially relevant in disorders such as Alzheimer’s disease, where pathological changes develop in brain regions that are also vulnerable to age-associated decline [10], [19], [22].

Distinguishing pathological neurodegeneration from normal aging is important for early diagnosis, disease monitoring, and biomarker development [23]. Conventional MRI-based assessments often rely on visual inspection or coarse volumetric measurements, which may not fully capture subtle alterations in tissue heterogeneity. Radiomics addresses this limitation by converting medical images into high-dimensional quantitative features that describe intensity, shape, and texture beyond visual interpretation [12], [39]. In neuroimaging, these features offer the potential to detect subtle structural and intensity variations that are not readily visible through conventional inspection [7].

Because aging and neurodegeneration affect multiple cortical and subcortical structures, a region-wise radiomic analysis may help determine whether disease-related alterations follow the same directional trends as normal aging or instead represent a distinct imaging phenotype. The present study adopts this region-wise, hypothesis-driven strategy: by first identifying radiomic features that vary systematically with age in cognitively normal individuals across anatomically defined brain regions, and subsequently examining whether Alzheimer’s disease produces consistent deviations from these age-associated patterns, we investigate whether Alzheimer’s disease manifests as amplification, divergence, or selective perturbation of normative structural aging processes. Features demonstrating consistent and systematic deviation from normative aging trajectories may represent biologically grounded candidates for age-adjusted biomarker development, particularly in a disease occurring within an already-aging population

Using anatomically guided brain-region segmentation, radiomic feature extraction, correlation analysis with age, and statistical comparison between normal and Alzheimer’s groups across age bins, we aim to characterise how Alzheimer’s disease modulates radiomic features sensitive to normative aging and to identify region-specific markers demonstrating quantitatively separable deviation from expected aging-associated patterns — features that may serve as candidate biomarkers for future age-adjusted assessment of Alzheimer’s disease and related neurodegenerative conditions.

## 2 Methodology

### 2.1 Dataset

Data used in the preparation of this article were obtained from the Alzheimer’s Disease Neuroimaging Initiative (ADNI) database (adni.loni.usc.edu). The ADNI was launched in 2003 as a public-private partnership, led by Principal Investigator Michael W. Weiner, MD. The primary goal of ADNI has been to test whether serial magnetic resonance imaging (MRI), positron emission tomography (PET), other biological markers, and clinical and neuropsychological assessment can be combined to measure the progression of mild cognitive impairment (MCI) and early Alzheimer’s disease (AD). For up-to-date information, see www.adni-info.org.

#### 2.1.1 Data Selection Method

To ensure consistency in data acquisition and reduce variability in imaging parameters [15], MRI scans were selected from the Alzheimer’s Disease Neuroimaging Initiative (ADNI) database for cognitively normal and Alzheimer’s disease subjects aged 60–90 using predefined filtering criteria (Table I) [20], [30], [33].

**TABLE I.** Data selection criteria used for filtering mri scans.

| Parameter | Selected Criteria |
| --- | --- |
| Project | ADNI |
| Research Groups | AD (Alzheimer’s disease), CN (Cognitively Normal) |
| Modality | MRI |
| Field Strength | 3 Tesla |
| Acquisition Plane | Sagittal |
| Acquisition Type | 3D |
| Manufacturer | Siemens |
| Image Weighting | T1-weighted |
| Image Sequence | MPRAGE |

#### 2.1.2 Details of the Image Data Selected for the Alzheimer’s Experiment

To construct a balanced dataset, subjects were stratified by sex (male and female) and further divided into predefined age bins. Within each sex-specific subgroup, samples from the cognitively normal (CN) and Alzheimer’s disease (AD) cohorts were selected such that the mean age and standard deviation within each age bin were closely matched between CN and AD subjects within that sex (Tables II, III, IV, and V).

**TABLE II.** Age distribution and sample size of selected male Alzheimer’s subjects.

| Age Group | Mean Age | Std. Dev. | Sample Size |
| --- | --- | --- | --- |
| 60–70 | 65.42 | 2.81 | 7 |
| 70–80 | 76.86 | 3.01 | 18 |
| 80–90 | 83.80 | 2.66 | 29 |

**TABLE III.** Age distribution and sample size of selected female alzheimer’s subjects.

| Age Group | Mean Age | Std. Dev. | Sample Size |
| --- | --- | --- | --- |
| 60–69 | 66.26 | 3.10 | 14 |
| 70–79 | 74.85 | 3.18 | 29 |
| 80–89 | 86.0 | 3.65 | 11 |

**TABLE IV.** Age distribution and sample size of selected cognitively normal male subjects.

| Age Group | Mean Age | Std. Dev. | Sample Size |
| --- | --- | --- | --- |
| 60–70 | 66.66 | 3.14 | 10 |
| 70–80 | 76.0 | 2.80 | 27 |
| 80–90 | 84.04 | 2.49 | 25 |

**TABLE V.** Age distribution and sample size of selected cognitively normal female subjects.

| Age Group | Mean Age | Std. Dev. | Sample Size |
| --- | --- | --- | --- |
| 60–69 | 67.88 | 1.93 | 7 |
| 70–79 | 74.81 | 3.39 | 25 |
| 80–89 | 86.00 | 2.78 | 7 |

This matching was performed to ensure comparable age distributions between CN and AD subjects within both male and female populations. Controlling for age in this manner reduces potential confounding effects, as normal aging itself can influence brain structure and imaging-derived features [4]. By aligning the age statistics across cohorts within each sex, the subsequent statistical comparisons are more likely to reflect disease-related differences rather than demographic variability, thereby improving the validity of the t-test analysis.

### 2.2 Proposed Methodology

The overall workflow of the proposed study consists of four main steps: brain region segmentation, radiomic feature extraction, correlation analysis with age, and identification of promising discriminative features for distinguishing Alzheimer’s disease from normal aging.

#### 2.2.1 Brain Region Segmentation

After selecting the MRI scans, brain region segmentation was performed to obtain anatomically meaningful regions of interest (ROIs).

- Insula
- Caudate Nucleus
- Putamen
- Rolandic Operculum
- Cingulum
- Thalamus
- Heschl’s Gyrus
- Vermis
- Hippocampus
- Entorhinal Cortex

First, eight regions—namely the insula, caudate nucleus, putamen, rolandic operculum, cingulum, thalamus, Heschl’s gyrus, and vermis—were selected based on prior computational research identifying these regions as informative for capturing age-associated structural variations in brain MRI-derived features [38].

In contrast, the hippocampus and entorhinal cortex were included based on established clinical and medical evidence, as they are among the most consistently implicated regions in Alzheimer’s disease, particularly in early neurodegenerative processes affecting memory-related structures [16]. Their inclusion ensures that the analysis is not limited to normative aging markers alone, but also captures canonical disease-vulnerable regions, enabling the evaluation of whether Alzheimer’s-related effects overlap with, diverge from, or extend beyond age-sensitive radiomic patterns.

Before finalizing the segmentation approach, preliminary experiments were conducted. Initially, a voxel-based morphometry–style pipeline was explored in which MRI scans were registered to the standard Montreal Neurological Institute (MNI) template, and the Automated Anatomical Labeling (AAL) atlas [29] was used for brain parcellation to define region-wise masks. Radiomic features were then extracted from these atlas-defined regions. However, due to spatial normalization during registration, the geometric shape of brain regions became standardized across subjects, leading to highly similar shape-based radiomic features and thereby reducing their discriminative value.

To address this, automated segmentation methods were evaluated using AssemblyNet [5], SynthSeg [3], and SLaNT [14]. Based on qualitative assessment of segmentation quality and anatomical consistency, and using the publicly available SLaNT implementation (github.com/MASILab/SLANTbrainSeg.git), MRI scans were segmented with SLaNT for the final analysis. The segmented regions were then used for subsequent radiomic feature extraction.

#### 2.2.2 Bias Field Correction

MRI scans were first subjected to intensity inhomogeneity correction prior to radiomic feature extraction. Although the dataset was filtered to maintain consistency in acquisition parameters (Table I), residual intensity non-uniformities may still arise due to RF coil sensitivity profiles, magnetic field inhomogeneity, and variations in subject positioning within the scanner. These effects can introduce smooth spatial intensity variations across the image, commonly referred to as a bias field [35].

Importantly, such intensity variations are primarily acquisition-related and are not expected to be systematically associated with disease status. However, they can increase variability in intensity-based radiomic features and reduce feature reproducibility across scans. To mitigate this effect and improve inter-subject intensity consistency, N4 bias field correction [28] was applied to all MRI volumes prior to subsequent preprocessing and radiomic feature extraction. This step reduces low-frequency intensity shading while preserving anatomical structures, thereby improving the stability and comparability of extracted radiomic features across subjects.

N4 bias field correction was performed on the MRI images using FSL FAST (FMRIB’s Automated Segmentation Tool) [17], [37] with the `-B` option enabled to estimate and correct intensity non-uniformity.

Voxel spacing was verified to be consistent across all included scans; therefore, spatial resampling was not performed [39].

#### 2.2.3 Intensity Normalization

To standardize intra-subject intensity distributions and reduce inter-subject variability, voxel-wise z-score intensity normalization was applied independently within the brain mask [11], whereby voxel intensities were centered and scaled using the mean and standard deviation of brain tissue voxels. This procedure transformed the original scanner-dependent intensity distributions into standardized values. Post-normalization inspection across all cohorts (Alzheimer’s and cognitively normal; male and female) demonstrated comparable intensity distributions without systematic inter-group shifts [6]. This observed consistency formed the basis for selecting discretization parameters in the subsequent radiomic feature extraction step. Accordingly, a fixed bin width of 1 was selected based on the observed effective intensity dynamic range, resulting in approximately 20 discrete gray levels [8]. This provided a balanced discretization, retaining relevant intensity variation without amplifying noise, and ensuring stable feature estimation across subjects, particularly for texture-based radiomic measures.

#### 2.2.4 Radiomic Feature Extraction

Radiomic features were extracted from the segmented brain regions to quantitatively characterize structural and intensity patterns within the MRI scans. Feature extraction was performed on the MRI images using the *PyRadiomics* library [25], [31]. Feature extraction was restricted to the original image type. All features from the first-order, shape, GLCM, GLRLM, GLSZM, GLDM, and NGTDM classes were enabled, yielding 107 radiomic features per ROI with default PyRadiomics settings retained. The extraction settings used in this study included a bin width of 1, PyRadiomics normalization disabled, and the mask label set to 1.

#### 2.2.5 Correlation Analysis with Age

To investigate age-related variations in radiomic characteristics, a correlation analysis was performed between each radiomic feature and the chronological age of the normal cohort subjects across all selected brain regions. Since ignoring sex as a covariate may cause radiomic features to capture sex-related variation rather than age-related or disease-specific patterns, and given that sex-related differences are known to influence brain structure and aging [26], the analysis was conducted separately for male and female populations. This stratification helped reduce the potential confounding effect of sex, thereby improving the interpretability of the identified candidate radiomic features.

Spearman’s rank correlation analysis [1] was used to evaluate the monotonic relationship between radiomic features and age. The analysis was implemented using the scipy.stats module in the SciPy library [32]. For each brain region, the correlation between each extracted feature and age was computed independently.

For every radiomic feature within each brain region, the following statistical hypotheses were tested:

Null hypothesis (*H*_0_): There is no monotonic association between the radiomic feature and age.

Alternative hypothesis (*H*_1_): There exists a significant monotonic association between the radiomic feature and age.

For each test, the Spearman correlation coefficient (*ρ*) and the corresponding p-value were calculated to assess the strength and statistical significance of the relationship. A significance level of *α* = 0.05 was used for hypothesis testing. By first identifying radiomic features that were strongly associated with age, this analysis defined a cross-sectional reference profile of age-sensitive radiomic variation within the cognitively normal cohort across the regions considered. These age-associated features were then used as candidate features for subsequent comparison between cognitively normal and Alzheimer’s groups, allowing the later disease-focused analysis to assess whether Alzheimer’s-related changes occurred in features that were also sensitive to age in the cognitively normal brain.

Spearman’s rank correlation was used instead of Pearson correlation as it evaluates monotonic relationships without assuming linearity or normality and is less sensitive to outliers. This makes it more appropriate for capturing potential nonlinear associations between age and radiomic features.

#### 2.2.6 Radiomic Feature Selection and Statistical Analysis

Because the primary aim of this study was to evaluate whether Alzheimer’s-related alterations occur within radiomic features that are also sensitive to normative aging, feature selection was restricted to age-associated features from the cognitively normal cohort. Features exhibiting moderate to moderately strong Spearman correlations (|*ρ*| = 0.4–0.65) with statistical significance [27] were retained for downstream analysis.

Subsequently, statistical comparisons were conducted between the normal and Alzheimer’s groups across different age bins. For each selected radiomic feature, group-wise comparisons were performed using an independent two-sample t-test with Welch’s correction [34], [36] to account for unequal variances between groups. The analysis was implemented using the `ttest_ind` function from the `scipy.stats` module of SciPy with the parameter `equal_var=False.`

To obtain robust estimates of statistical significance and reduce sensitivity to potential deviations from parametric assumptions, permutation-based p-value estimation [9], [32] was employed. Specifically, 10,000 permutations were performed, where group labels were randomly reassigned and the test statistic recomputed to generate an empirical null distribution.

The statistical hypotheses evaluated in this study were defined as: Null hypothesis (*H*_0_): The mean value of the radiomic feature is equal between the normal and Alzheimer’s groups (*µ*_normal_ = *µ*_AD_).

Alternative hypothesis (*H*_1_): The mean value of the radiomic feature differs between the normal and Alzheimer’s groups (*µ*_normal_ ≠ *µ*_AD_).

All statistical tests were conducted as two-tailed tests. Radiomic features were further screened for their ability to preserve group-wise separation across age strata, identifying those that reflect a consistent disease-specific signal beyond normal biological variation.

## 3 Experiments and evaluation

### 3.1 Experimental Setup

#### 3.1.1 Setup for Age-Associated Radiomic Analysis

The Spearman correlation framework described in Section 2.2.5 was applied across 10 brain regions, with 107 radiomic features per region, resulting in a total of 1,070 statistical tests. To control for multiple comparisons, Benjamini–Hochberg false discovery rate (FDR) correction was performed within each brain region [2]. Radiomic features satisfying the correlation criteria defined in Section 2.2.6 were first filtered based on adjusted p-values below 0.05, and subsequently ranked according to the magnitude of the Spearman correlation coefficient (| *ρ*|). The top 10 features per region were then carried forward as the most robust age-associated radiomic signatures for downstream analysis.

#### 3.1.2 Experimental Design for Statistical Comparisons

To analyze how radiomic feature distributions vary across age bins and diagnostic groups, the dataset was stratified into discrete age bins. For each selected radiomic feature, the corresponding normal and Alzheimer’s groups were compared within each bin, resulting in 30 statistical comparisons per brain region (10 features × 3 bins) and a total of 300 tests across all regions.

Group-wise comparisons were performed using Welch’s two-sample t-test within each matched age bin. Statistical significance was assessed using permutation-based p-values to ensure robustness under potential deviations from normality and heterogeneous group sizes.

To control for multiple comparisons, Benjamini–Hochberg FDR correction was applied independently within each brain region across the 30 tests performed per region. Only comparisons with sufficient sample sizes in both groups were included to ensure stable variance estimation.

This framework enabled a comprehensive evaluation of radiomic feature differences between normal aging and Alzheimer’s disease across the lifespan.

## 4 Results

### 4.1 Spearman Correlation with Aging

Tables VI and VII summarize all age-associated radiomic features that showed significant monotonic relationships in the cognitively normal cohort.

**TABLE VI.** Complete summary of age-associated radiomic features identified by Spearman correlation analysis in the cognitively normal cohort (Part I)

| Region | Feature | $\rho$ | $p$-value | Trend |
| --- | --- | --- | --- | --- |
| Cingulum | Median | 0.64 | < 0.001 | Increase |
|  | Interquartile Range | 0.57 | < 0.001 | Increase |
|  | Robust Mean Absolute Deviation | 0.56 | < 0.001 | Increase |
|  | Kurtosis | -0.60 | < 0.001 | Decrease |
|  | Energy | -0.45 | < 0.001 | Decrease |
|  | Gray Level Non-Uniformity Normalized | 0.49 | < 0.001 | Increase |
| Entorhinal Cortex | Cluster Shade | -0.49 | ≤ 0.001 | Decrease |
| | Skewness | -0.43 | ≤ 0.001 | Decrease |
| | Surface Volume Ratio | 0.42 | ≤ 0.001 | Increase |
| | Median | 0.41 | ≤ 0.001 | Increase |
| Hippocampus | Cluster Shade | -0.60 | < 0.001 | Decrease |
|  | Gray Level Non-Uniformity | -0.53 | < 0.001 | Decrease |
|  | Uniformity | -0.53 | < 0.001 | Decrease |
|  | Cluster Tendency | 0.52 | < 0.001 | Increase |
|  | MCC | 0.50 | < 0.001 | Increase |
|  | Entropy | 0.47 | < 0.001 | Increase |
| Heschl's Gyrus | Minimum | 0.53 | ≤ 0.001 | Increase |
| | 10th Percentile | 0.43 | ≤ 0.001 | Increase |
| | Idmn | -0.42 | ≤ 0.01 | Decrease |
| | Kurtosis | -0.42 | ≤ 0.01 | Decrease |
| | Run Entropy | -0.40 | ≤ 0.01 | Decrease |
|  | NGTDM Contrast | 0.41 | < 0.01 | Increase |
| Insula | Run Entropy | -0.56 | < 0.001 | Decrease |
|  | Root Mean Squared | -0.51 | < 0.001 | Decrease |
|  | Gray Level Variance | -0.48 | < 0.001 | Decrease |
|  | Median | 0.49 | < 0.001 | Increase |
|  | Minimum | 0.45 | < 0.001 | Increase |

**TABLE VII.** Part ii (continued)

| Region | Feature | $\rho$ | $p$-value | Trend |
| --- | --- | --- | --- | --- |
| Putamen | Median | 0.66 | < 0.001 | Increase |
|  | Mean | 0.65 | < 0.001 | Increase |
|  | Maximum Probability | 0.60 | < 0.001 | Increase |
|  | Uniformity | 0.59 | < 0.001 | Increase |
|  | Dependence Entropy | -0.60 | < 0.001 | Decrease |
|  | Gray Level Non-Uniformity | -0.53 | < 0.001 | Decrease |
| Vermis | Elongation | -0.48 | < 0.001 | Decrease |
|  | Imc1 | -0.46 | < 0.001 | Decrease |
|  | Kurtosis | -0.44 | < 0.001 | Decrease |
|  | Imc2 | 0.44 | < 0.001 | Increase |
|  | MCC | 0.44 | < 0.001 | Increase |
|  | Correlation | 0.43 | < 0.001 | Increase |
| Rolandic Operculum | Kurtosis | -0.53 | < 0.001 | Decrease |
|  | Idmn | -0.43 | < 0.01 | Decrease |
|  | Maximum Probability | -0.42 | < 0.001 | Decrease |
|  | NGTDM Contrast | 0.44 | < 0.001 | Increase |
|  | Minimum | 0.43 | < 0.01 | Increase |
| Thalamus | Gray Level Non-Uniformity Normalized | -0.67 | < 0.001 | Decrease |
|  | Gray Level Variance | 0.65 | < 0.001 | Increase |
|  | Cluster Prominence | 0.59 | < 0.001 | Increase |
|  | Variance | 0.57 | < 0.001 | Increase |
|  | Root Mean Squared | 0.55 | < 0.001 | Increase |
| Caudate Nucleus | Uniformity | -0.67 | < 0.001 | Decrease |
|  | Run Variance | -0.66 | < 0.001 | Decrease |
|  | Joint Energy | -0.66 | < 0.001 | Decrease |
|  | Sum Squares | 0.67 | < 0.001 | Increase |
|  | Cluster Tendency | 0.66 | < 0.001 | Increase |
|  | Median | 0.66 | < 0.001 | Increase |

The age-associated radiomic patterns also reveal distinct region-specific signatures. Several subcortical regions show a consistent trend toward increased signal intensity and texture regularization. In the putamen, this manifests as a uniform upward shift in T1 signal intensity with progressive texture homogenization. The thalamus similarly shows rising signal intensity, but accompanied by a widening and increased irregularity of the intensity distribution. The caudate nucleus follows a related pattern, with increasing signal intensity alongside local contrast heterogeneity and fragmentation of homogeneous texture zones.

In contrast, cortical and limbic regions tend toward greater textural complexity with age. In the cingulum, the intensity distribution shifts upward while simultaneously broadening and flattening, with progressive fragmentation of homogeneous texture zones. The hippocampus shows greater intensity variance and entropy alongside stronger spatial clustering, reflecting an increasingly heterogeneous texture pattern. The insula exhibits a narrowing and upward shift of the intensity distribution alongside increasingly fragmented and locally heterogeneous texture. In the entorhinal cortex, a slight upward shift in signal intensity, reduced distributional skewness, and an increased surface-to-volume ratio.

The remaining regions present more specific profiles. In Heschl’s gyrus, aging is associated with a compressed intensity range, increased local contrast, and a more uniform texture pattern. The rolandic operculum shows a similar dynamic range narrowing with increased local inter-voxel contrast and reduced texture homogeneity. In the vermis, aging is associated with morphological changes suggestive of progressive shape compaction, together with flattening of the intensity distribution, increasing spatial correlation between neighboring voxels, and growth of extended low-intensity zones.

A complete list of Spearman correlation coefficients, raw *p*values, FDR-corrected *p*-values, and significance decisions for all reported regions is provided in the Supplementary Material, Section 2.

### 4.2 T-Test Results for Normal and Alzheimer’s Cohorts

Hippocampus: Significant differences between the normal and Alzheimer’s groups were observed for multiple radiomic features in the hippocampus across all age bins (60–70, 70–80, and 80–90 years). These included Gray Level Non-Uniformity and Maximal Correlation Coefficient (MCC) (all *p <* 0.05). Entorhinal cortex: Cluster Shade and Surface Volume Ratio demonstrated significant differences between the normal and Alzheimer’s groups across all age bins (all *p <* 0.05). In the female subgroup, Short Run Low Gray Level Emphasis and Contrast also showed significant differences between groups. Cingulum: Kurtosis and Gray Level Non-Uniformity Normalized showed significant group differences across all age bins (all *p <* 0.05). In the female subgroup, Kurtosis showed significant differences between groups. The corresponding t-statistic (*t*) value and FDR-adjusted *p* values for these radiomic features are summarized in Table VIII.

However, the other regions showed more heterogeneous patterns, with statistically significant effects often limited to specific age bins rather than consistently preserved across the evaluated age range. The complete Welch t-test results for all evaluated regions and age bins are provided in the Supplementary Material, Section 3.

Given that female subgroup sample sizes within individual age bins were insufficient for stable feature identification, the primary analysis focuses on the male cohort; female subgroup Spearman correlation and t-test findings are provided as supportive evidence in the Supplementary Material, Section 6.

### 4.3 Effect Size and Discriminative Performance of Selected Features

To quantify the magnitude and discriminative ability of radiomic feature differences, effect sizes were computed using Cohen’s d and Hedges’ g. Feature-level receiver operating characteristic (ROC) analysis was performed to estimate the area under the curve (AUC) for distinguishing Alzheimer’s disease from cognitively normal subjects within each age bin, with 95% confidence intervals computed using bootstrap resampling. The corresponding results are summarized in Table IX, and complete AUC, Cohen’s *d*, and Hedges’ *g* results are provided in the Supplementary Material, Section 4.

### 4.4 Discussion

The present study aimed to determine whether Alzheimer’s disease manifests as amplification, divergence, or selective perturbation of radiomic features associated with normative structural aging processes across anatomically distinct brain regions. The results demonstrate that Alzheimer’s disease does not uniformly mimic normal aging across the brain, but instead exhibits region-specific relationships with age-sensitive radiomic patterns. In particular, medial temporal and limbic regions, including the hippocampus, entorhinal cortex, and cingulum, showed age-aligned but consistently shifted radiomic features across age bins, indicating that Alzheimer’s disease modulates imaging markers that are also associated with normative aging in these structures. Collectively, these findings suggest that Alzheimer’s disease selectively perturbs normative aging-related radiomic trajectories within specific structural systems rather than producing globally consistent aging-related radiomic alterations across the brain.

#### Cingulum

The observed positive correlation of cingulum Gray Level Non-Uniformity Normalized feature with age is reflected in an increasing trend across age bins in both groups (Fig. 1), with the Alzheimer’s group consistently demonstrating higher mean values than the normal group and maintaining statistically significant differences across all bins. Across the age range, the Alzheimer’s group remains shifted toward higher values, indicating persistently elevated Gray Level Non-Uniformity relative to normal aging. In contrast, the cingulum firstorder kurtosis feature exhibits a decreasing trend across age bins in both groups, with the Alzheimer’s group consistently demonstrating lower mean values than the normal group.

**Fig. 1.**
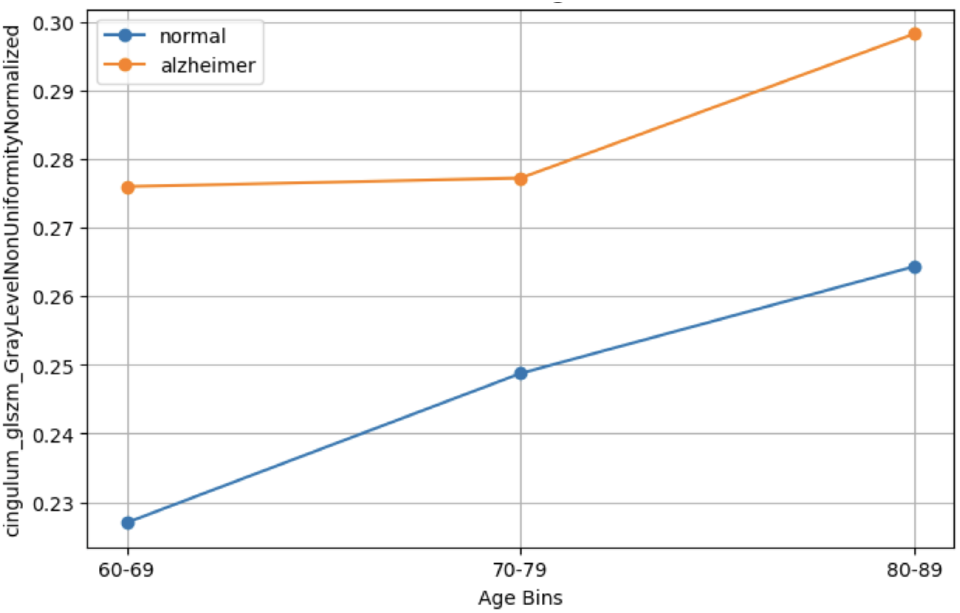
Cingulum gray level non-uniformity normalized feature values. The plot compares age-bin trends between cognitively normal and Alzheimer’s disease groups.

For entorhinal cortex GLCM Cluster Shade, both groups exhibit a monotonic decline across age bins, consistent with an age-associated reduction in these texture features. Despite this parallel trend, the Alzheimer’s group consistently demonstrates lower mean values than the normal group across all age ranges, indicating a stable disease-related shift. Additionally, the observed increase in standard deviation with age reflects greater inter-subject variability in tissue characteristics in older populations.

The surface-to-volume ratio of the entorhinal cortex increases with age in both groups, consistent with age-related morphological alterations. Across all age bins, the Alzheimer’s group consistently exhibits higher values compared to controls (Fig. 2(a)), indicating a stable disease-associated shift in structural geometry. As shown in Figure 2 (panel b), the distributions exhibit increasing spread with age, leading to greater overlap between groups in the oldest age bin. This increased overlap appears to be driven by rising inter-subject variability rather than a convergence of central tendency, as the Alzheimer’s group remains consistently elevated across all age ranges.

**Fig. 2.**
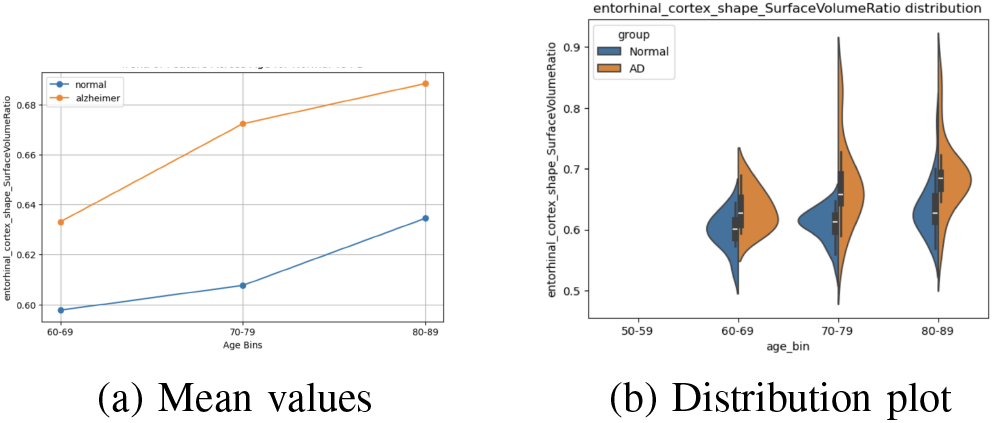
Comparison of the radiomic feature entorhinal cortex shape SurfaceVolumeRatio in the normal and Alzheimer’s groups: (a) mean values across age bins and (b) distributions across age bins.

For the hippocampus, radiomic features including GLDM Gray Level Non-Uniformity exhibit lower mean values in the Alzheimer’s group than in the normal group across the evaluated age bins. In contrast, the GLCM MCC feature demonstrates higher mean values in the Alzheimer’s cohort relative to controls.

The remaining regions did not demonstrate consistent adherence to these patterns. Although statistically significant differences were observed in certain cases, these effects were often confined to specific age bins and were not consistently preserved across the evaluated age ranges. Such findings suggest that while partial group separability may exist in these regions, the observed disease-related alterations are less stable and less uniformly expressed than those observed in the hippocampus, entorhinal cortex, and cingulum.

#### 4.4.1 Disease-Continuum Analysis Using LMCI

To further examine where the radiomic markers identified in the primary analysis become discriminative along the disease continuum, a secondary analysis was performed using the late mild cognitive impairment (LMCI) cohort from ADNI.

The LMCI cohort was acquired using the same MRI acquisition criteria, preprocessing pipeline, and radiomics extraction procedure described in Sections 2.1 and 2.2. The age-bin summary statistics for the LMCI group are reported in Table X.

Using the same set of previously selected radiomic features, two pairwise group comparisons were conducted: CN versus LMCI and LMCI versus AD. For each comparison, Welch’s two-sample t-test was applied, and p-values were corrected for multiple comparisons within each region using the false discovery rate (FDR). Based on statistical significance across the two comparisons, features were categorized as early-stage markers (significant in CN versus LMCI), late-stage markers (significant only in LMCI versus AD), or progressive markers (significant in both comparisons). The results of this experiment are summarized in Table XI, with the complete Normal–LMCI and LMCI–Alzheimer t-test statistics provided in the Supplementary Material, Section 5.

Building on these region-specific findings, the LMCI analysis further refines this interpretation by showing that LMCI occupies a distinct radiomic position relative to normal aging, with significant textural and morphological differences detected across the hippocampus, cingulum, and entorhinal cortex, predominantly within the 70–80 and 80–90 age bins (Table XI). In contrast, the LMCI and Alzheimer’s disease groups were not separated by statistically significant differences across the evaluated features and regions.The absence of significant LMCI–AD separation suggests that the structural divergence captured by these radiomic markers emerges early along the disease continuum, at the transition from normal cognition to LMCI, and does not progress further in a radiomically detectable manner within the present feature set (Fig. 3).

**Fig. 3.**
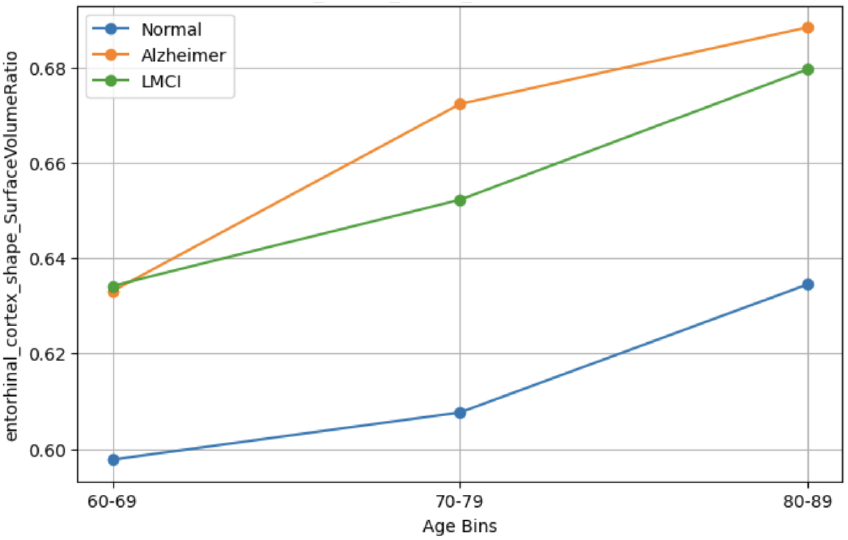
Age-bin trend of entorhinal cortex surface volume ratio across cognitively normal, LMCI, and Alzheimer’s disease groups. Both LMCI and Alzheimer’s disease show elevated values relative to cognitively normal subjects, particularly in the older age bins, while LMCI and Alzheimer’s disease remain comparatively close.

Collectively, these findings position the identified radiomic markers as candidate early-stage biomarkers of neurodegener-ation rather than markers of later disease progression.

An important limitation of the LMCI analysis is the biological heterogeneity of the ADNI LMCI cohort, since not all LMCI subjects necessarily represent prodromal Alzheimer’s disease. This heterogeneity may contribute to overlapping radiomic patterns and reduced group separability between LMCI and Alzheimer’s disease. Future studies incorporating amyloid PET biomarkers may help define biologically more homogeneous early-stage cohorts and further clarify the relationship between the identified radiomic markers and underlying Alzheimer’s pathology.

A deliberate design choice of the present study was to prioritize age-associated radiomic markers identified in the cognitively normal cohort. This age-referenced framework enhanced sensitivity to Alzheimer’s-related effects occurring within features linked to normative aging, while future investigations may broaden the search space to include disease-specific radiomic alterations beyond these markers.

Acquisition-level harmonization was implemented to reduce technical variability; however, factors such as head size, motion, and clinical heterogeneity may still contribute modest residual variance. Incorporating these variables into larger and more comprehensive future models may further refine the robustness of observed associations.

Furthermore, sample sizes in certain younger age-bin cohorts were relatively small, and exceptionally high discriminative performance estimates in these bins should therefore be interpreted with caution.

Finally, all findings are derived from a single dataset (ADNI). While the consistency of the identified radiomic markers across age bins and the LMCI staging analysis supports their robustness within this cohort, external validation in an independent dataset represents an important next step. Validation in a cohort such as OASIS-3 is a planned aim of follow-up work, with the goal of further establishing the generalizability of these markers.

Collectively, these findings highlight the potential of regionspecific radiomic trajectories to improve differentiation between healthy aging and pathological neurodegeneration, and lay the groundwork for future age-adjusted imaging biomarker reference ranges. Radiomic markers demonstrating consistent stage-wise separation in this study (Tables VIII and IX) represent promising candidates for this purpose. Features showing both normative age sensitivity and stable disease-associated deviation could ultimately support neuroimaging frameworks in which patient measurements are interpreted relative to expected values for cognitively normal individuals of similar age, rather than against a single global threshold. Establishing such reference ranges will require validation in larger, independent cohorts, including external datasets beyond ADNI, to confirm the generalizability of the identified markers.

**TABLE VIII.** T-test results for statistical comparison of radiomic features between cognitively normal and alzheimer’s disease cohorts across age bins.

| Region | Feature | Age bin | *t* value | FDR-adjusted *p* value |
| --- | --- | --- | --- | --- |
| Entorhinal cortex | Cluster shade | 60–69 | 2.85 | 0.013 |
|  |  | 70–79 | 4.64 | 0.001 |
|  |  | 80–89 | 3.88 | 0.001 |
|  | Surface volume ratio | 60–69 | -2.21 | 0.042 |
|  |  | 70–79 | -3.73 | 0.001 |
|  |  | 80–89 | -3.69 | 0.001 |
| Hippocampus | GLNU | 60–69 | 2.63 | 0.029 |
|  |  | 70–79 | 6.28 | 0.000 |
|  |  | 80–89 | 6.72 | 0.000 |
|  | MCC | 60–69 | -3.76 | 0.000 |
|  |  | 70–79 | -2.69 | 0.012 |
|  |  | 80–89 | -4.78 | 0.000 |
| Cingulum | Kurtosis | 60–69 | 3.09 | 0.025 |
|  |  | 70–79 | 3.11 | 0.015 |
|  |  | 80–89 | 5.75 | 0.006 |
|  | GLNU normalised | 60–69 | -2.34 | 0.036 |
|  |  | 70–79 | -2.11 | 0.048 |
|  |  | 80–89 | -3.36 | 0.011 |

Reported *p*-values are FDR-adjusted.

**TABLE IX.**
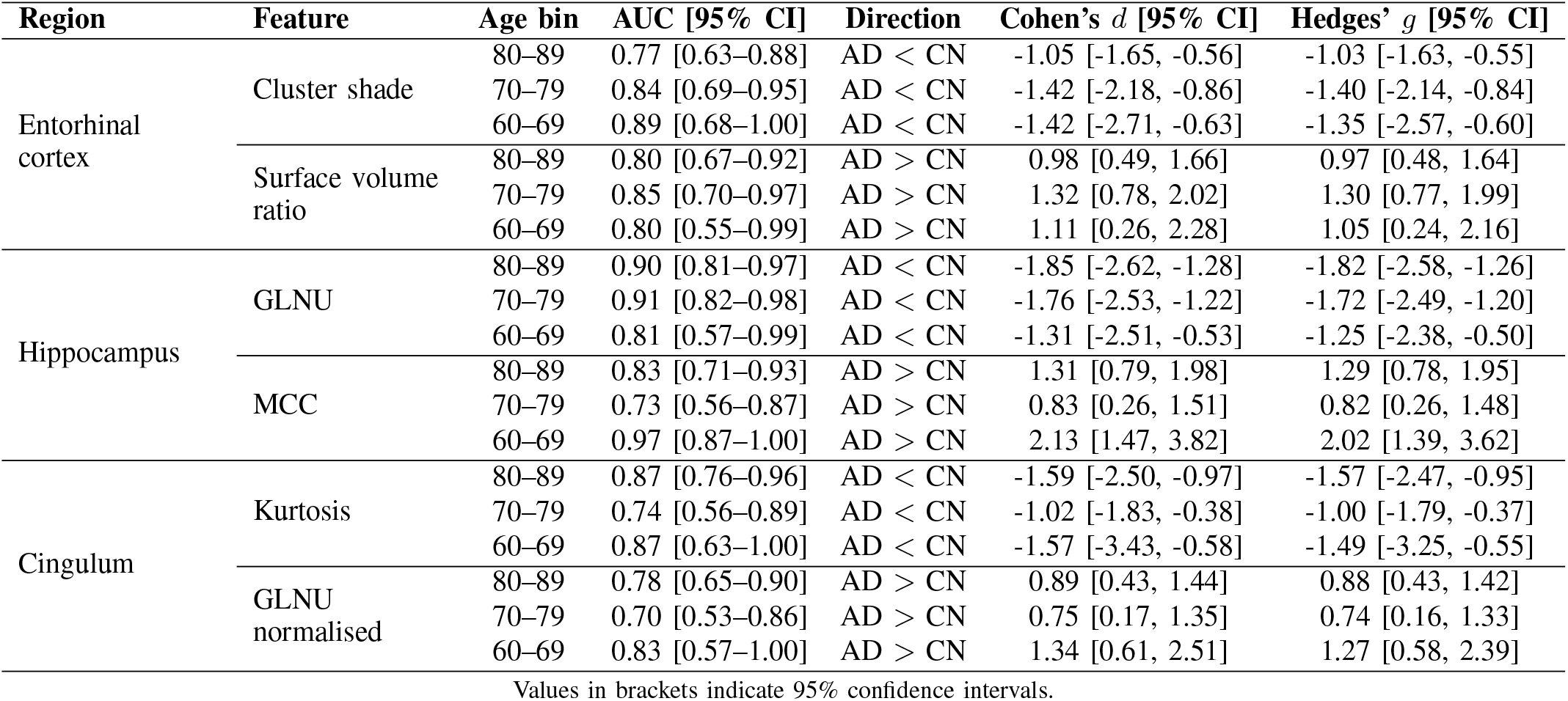
Discriminative performance and effect size of selected radiomic features across age bins for the normal and alzheimer’s disease cohorts.

| Region | Feature | Age bin | AUC [95% CI] | Direction | Cohen's *d* [95% CI] | Hedges' *g* [95% CI] |
| --- | --- | --- | --- | --- | --- | --- |
| Entorhinal cortex | Cluster shade | 80–89 | 0.77 [0.63–0.88] | AD < CN | -1.05 [-1.65, -0.56] | -1.03 [-1.63, -0.55] |
|  |  | 70–79 | 0.84 [0.69–0.95] | AD < CN | -1.42 [-2.18, -0.86] | -1.40 [-2.14, -0.84] |
|  |  | 60–69 | 0.89 [0.68–1.00] | AD < CN | -1.42 [-2.71, -0.63] | -1.35 [-2.57, -0.60] |
|  | Surface volume ratio | 80–89 | 0.80 [0.67–0.92] | AD > CN | 0.98 [0.49, 1.66] | 0.97 [0.48, 1.64] |
|  |  | 70–79 | 0.85 [0.70–0.97] | AD > CN | 1.32 [0.78, 2.02] | 1.30 [0.77, 1.99] |
|  |  | 60–69 | 0.80 [0.55–0.99] | AD > CN | 1.11 [0.26, 2.28] | 1.05 [0.24, 2.16] |
| Hippocampus | GLNU | 80–89 | 0.90 [0.81–0.97] | AD < CN | -1.85 [-2.62, -1.28] | -1.82 [-2.58, -1.26] |
|  |  | 70–79 | 0.91 [0.82–0.98] | AD < CN | -1.76 [-2.53, -1.22] | -1.72 [-2.49, -1.20] |
|  |  | 60–69 | 0.81 [0.57–0.99] | AD < CN | -1.31 [-2.51, -0.53] | -1.25 [-2.38, -0.50] |
|  | MCC | 80–89 | 0.83 [0.71–0.93] | AD > CN | 1.31 [0.79, 1.98] | 1.29 [0.78, 1.95] |
|  |  | 70–79 | 0.73 [0.56–0.87] | AD > CN | 0.83 [0.26, 1.51] | 0.82 [0.26, 1.48] |
|  |  | 60–69 | 0.97 [0.87–1.00] | AD > CN | 2.13 [1.47, 3.82] | 2.02 [1.39, 3.62] |
| Cingulum | Kurtosis | 80–89 | 0.87 [0.76–0.96] | AD < CN | -1.59 [-2.50, -0.97] | -1.57 [-2.47, -0.95] |
|  |  | 70–79 | 0.74 [0.56–0.89] | AD < CN | -1.02 [-1.83, -0.38] | -1.00 [-1.79, -0.37] |
|  |  | 60–69 | 0.87 [0.63–1.00] | AD < CN | -1.57 [-3.43, -0.58] | -1.49 [-3.25, -0.55] |
|  | GLNU normalised | 80–89 | 0.78 [0.65–0.90] | AD > CN | 0.89 [0.43, 1.44] | 0.88 [0.43, 1.42] |
|  |  | 70–79 | 0.70 [0.53–0.86] | AD > CN | 0.75 [0.17, 1.35] | 0.74 [0.16, 1.33] |
|  |  | 60–69 | 0.83 [0.57–1.00] | AD > CN | 1.34 [0.61, 2.51] | 1.27 [0.58, 2.39] |

Values in brackets indicate 95% confidence intervals.

**TABLE X.** Age distribution and sample size of selected LMCI subjects.

| Age Group | Mean Age | Std. Dev. | Sample Size |
| --- | --- | --- | --- |
| 60–70 | 65.7 | 2.45 | 10 |
| 70–80 | 75.4 | 2.83 | 20 |
| 80–90 | 82.2 | 2.14 | 14 |

**TABLE XI.**
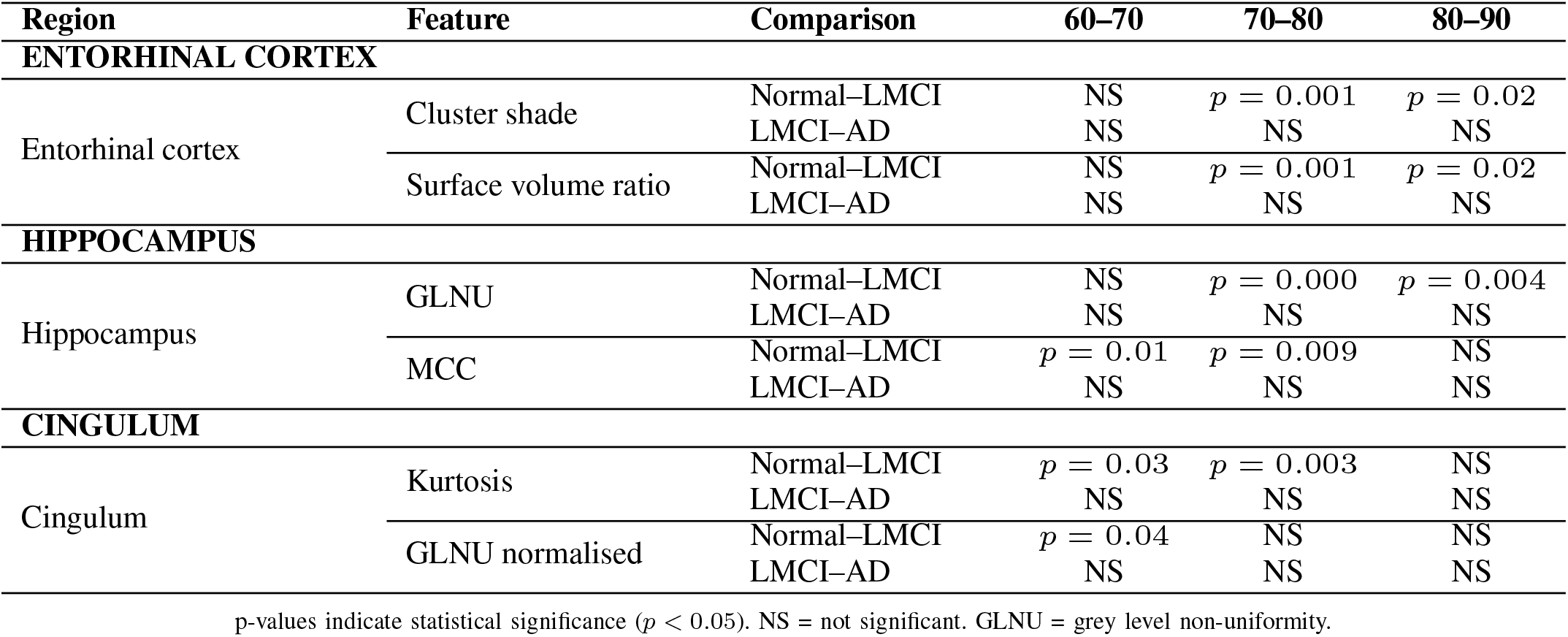
T-test significance of radiomic features across age bins for normal–lmci and lmci–alzheimer comparisons in hippocampus, cingulum, and entorhinal cortex.

In this regard, the fully automated nature of the present pipeline may represent a practical advantage for future largescale validation efforts. Because the analytical pipeline—from automated brain segmentation through radiomic feature extraction to age-stratified statistical comparison—is fully computational and requires no manual annotation, it can be readily applied across large imaging cohorts with minimal manual intervention.

Following appropriate external validation, these markers may ultimately contribute to age-adjusted Alzheimer’s disease screening and early detection

## CRediT authorship contribution statement

Megha Suman Sharma: Conceptualization (supporting), Data curation, Formal analysis, Investigation, Methodology, Software, Visualization, Writing – original draft, Writing – review and editing.

Rajat Agarwal: Conceptualization (lead), Funding acquisition, Formal analysis, Investigation, Methodology, Project administration, Resources, Supervision, Writing – review and editing. Nikita Tiwari: Software, Validation, Writing – review and editing.

Megha Sharma: Validation, Writing – review and editing. Ajay Kaushik: Funding acquisition, Resources, Writing – review and editing.

## Data availability

The MRI data used in this study are available from the Alzheimer’s Disease Neuroimaging Initiative (ADNI) database at adni.loni.usc.edu upon registration and approval, subject to the ADNI Data Use Agreement. The analysis code used in this study is available from the corresponding author upon reasonable request.

## Author note

Megha Suman Sharma (co-first author) and Megha Sharma (fourth author) are distinct individuals affiliated with Radpretation Technologies in different roles. They share a partial name similarity but are separate contributors to this work.

Acknowledgment

The authors thank Dr. Ajay Aggarwal, Consultant Radiologist and former Director at the Diwanchand Aggarwal X-ray and Imaging Research Centre, New Delhi, for his expert radiological review and valuable clinical insights during the interpretation of neuroimaging findings and manuscript development.

Data collection and sharing for the Alzheimer’s Disease Neuroimaging Initiative (ADNI) is funded by the National Institute on Aging (National Institutes of Health Grant U19 AG024904). The grantee organization is the Northern California Institute for Research and Education.

In the past, ADNI has also received funding from the National Institute of Biomedical Imaging and Bioengineering, the Canadian Institutes of Health Research, and private sector contributions through the Foundation for the National Institutes of Health (FNIH) including generous contributions from the following: AbbVie, Alzheimer’s Association; Alzheimer’s Drug Discovery Foundation; Araclon Biotech; BioClinica, Inc.; Biogen; Bristol-Myers Squibb Company; CereSpir, Inc.; Cogstate; Eisai Inc.; Elan Pharmaceuticals, Inc.; Eli Lilly and Company; EuroImmun; F. Hoffmann-La Roche Ltd and its affiliated company Genentech, Inc.; Fujirebio; GE Healthcare; IXICO Ltd.; Janssen Alzheimer Immunotherapy Research & Development, LLC.; Johnson & Johnson Pharmaceutical Research & Development LLC.; Lumosity; Lundbeck; Merck & Co., Inc.; Meso Scale Diagnostics, LLC.; NeuroRx Research; Neurotrack Technologies; Novartis Pharmaceuticals Corporation; Pfizer Inc.; Piramal Imaging; Servier; Takeda Pharmaceutical Company; and Transition Therapeutics.

### Declaration of generative AI and AI-assisted technologies in the manuscript preparation process

During the preparation of this work, the authors used ChatGPT and Claude for language refinement and editorial assistance, and Prism for assistance with manuscript formatting in LaTeX. After using these tools, the authors reviewed and edited the content as needed and take full responsibility for the content of the published article.

