## Supplementary Data for "Alzheimer’s Disease Selectively Perturbs Age-Sensitive Brain Radiomic Features Across the Disease Continuum"

#### Purpose of This Document

This supplementary document presents extended statistical results supporting the main manuscript. Specifically, it reports complete correlation analyses, group comparison statistics, receiver operating characteristic performance metrics, effect size estimates, and subgroup analyses for the evaluated radiomic features. The purpose of this document is to provide additional methodological transparency and allow readers to examine the full set of supporting results without expanding the main manuscript.

#### 1 Supplementary Tables

#### 2 Complete Spearman Correlation Results

This section reports the complete Spearman correlation results for radiomic features across the evaluated brain regions. For each region, the tables list radiomic features that showed high Spearman correlation values together with statistically significant raw $p$-values after false discovery rate (FDR) correction.

##### *2.1 Caudate Nucleus*

TABLE I
Spearman correlation results for significant caudate nucleus radiomic features.

| Radiomic feature | Spearman $\rho$ | $p$-value | FDR-corrected $p$ | Significant |
|---|---|---|---|---|
| caudate_nucleus_firstorder_Uniformity | -0.672873 | $2.091194 \times 10^{-9}$ | $8.066257 \times 10^{-8}$ | True |
| caudate_nucleus_glcm_SumSquares | 0.669922 | $2.607316 \times 10^{-9}$ | $8.066257 \times 10^{-8}$ | True |
| caudate_nucleus_glrlm_RunVariance | -0.663994 | $4.030698 \times 10^{-9}$ | $8.066257 \times 10^{-8}$ | True |
| caudate_nucleus_glcm_JointEnergy | -0.662354 | $4.539021 \times 10^{-9}$ | $8.066257 \times 10^{-8}$ | True |
| caudate_nucleus_gldm_LargeDependenceEmphasis | -0.661900 | $4.690187 \times 10^{-9}$ | $8.066257 \times 10^{-8}$ | True |
| caudate_nucleus_glrlm_RunPercentage | 0.660513 | $5.182198 \times 10^{-9}$ | $8.066257 \times 10^{-8}$ | True |
| caudate_nucleus_glcm_ClusterTendency | 0.657334 | $6.500087 \times 10^{-9}$ | $8.066257 \times 10^{-8}$ | True |
| caudate_nucleus_firstorder_Median | 0.657006 | $6.652818 \times 10^{-9}$ | $8.066257 \times 10^{-8}$ | True |
| caudate_nucleus_glrlm_LongRunEmphasis | -0.656729 | $6.784702 \times 10^{-9}$ | $8.066257 \times 10^{-8}$ | True |
| caudate_nucleus_firstorder_90Percentile | 0.650523 | $1.046900 \times 10^{-8}$ | $1.067356 \times 10^{-7}$ | True |

##### *2.2 Cingulum*

TABLE II
Spearman correlation results for significant cingulum radiomic features.

| Radiomic feature | Spearman $\rho$ | $p$-value | FDR-corrected $p$ | Significant |
|---|---|---|---|---|
| cingulum_firstorder_Median | 0.638566 | $2.349524 \times 10^{-8}$ | 0.000003 | True |
| cingulum_firstorder_Kurtosis | -0.602140 | $2.252737 \times 10^{-7}$ | 0.000012 | True |
| cingulum_firstorder_InterquartileRange | 0.566244 | $1.612222 \times 10^{-6}$ | 0.000058 | True |
| cingulum_firstorder_RobustMeanAbsoluteDeviation | 0.557818 | $2.475911 \times 10^{-6}$ | 0.000066 | True |
| cingulum_firstorder_Mean | 0.552798 | $3.179322 \times 10^{-6}$ | 0.000068 | True |
| cingulum_glszm_GrayLevelNonUniformityNorm. | 0.493114 | $4.661966 \times 10^{-5}$ | 0.000831 | True |
| cingulum_firstorder_Minimum | 0.485597 | $6.317576 \times 10^{-5}$ | 0.000952 | True |
| cingulum_firstorder_90Percentile | 0.482595 | $7.118692 \times 10^{-5}$ | 0.000952 | True |
| cingulum_firstorder_MeanAbsoluteDeviation | 0.463423 | $1.487392 \times 10^{-4}$ | 0.001768 | True |
| cingulum_firstorder_Energy | -0.446371 | $2.764505 \times 10^{-4}$ | 0.002958 | True |

#### 2.3 *Rolandic Operculum*

TABLE III
SPEARMAN CORRELATION RESULTS FOR SIGNIFICANT ROLANDIC OPERCULUM RADIOMIC FEATURES.

| Radiomic feature | Spearman $\rho$ | $p$-value | FDR-corrected $p$ | Significant |
|---|---|---|---|---|
| rolandic_operculum_firstorder_Kurtosis | -0.527951 | 0.000010 | 0.001107 | True |
| rolandic_operculum_ngtdm_Contrast | 0.438576 | 0.000363 | 0.011798 | True |
| rolandic_operculum_firstorder_Minimum | 0.434338 | 0.000420 | 0.011798 | True |
| rolandic_operculum_glcm_Idmn | -0.427729 | 0.000525 | 0.011798 | True |
| rolandic_operculum_firstorder_Range | -0.426266 | 0.000551 | 0.011798 | True |
| rolandic_operculum_glcm_MaximumProbability | -0.418849 | 0.000704 | 0.012549 | True |
| rolandic_operculum_glcm_Idn | -0.407422 | 0.001014 | 0.015500 | True |
| rolandic_operculum_glszm_GLN... | 0.390369 | 0.001709 | 0.022540 | True |
| rolandic_operculum_glszm_ZoneEntropy | -0.386863 | 0.001896 | 0.022540 | True |
| rolandic_operculum_glrlm_LRHGLE... | -0.360981 | 0.003947 | 0.039972 | True |

#### 2.4 *Putamen*

TABLE IV
SPEARMAN CORRELATION RESULTS FOR SIGNIFICANT PUTAMEN RADIOMIC FEATURES.

| Radiomic feature | Spearman $\rho$ | $p$-value | FDR-corrected $p$ | Significant |
|---|---|---|---|---|
| putamen_firstorder_Median | 0.663464 | $4.188709 \times 10^{-9}$ | $4.191696 \times 10^{-7}$ | True |
| putamen_firstorder_Mean | 0.654685 | $7.834946 \times 10^{-9}$ | $4.191696 \times 10^{-7}$ | True |
| putamen_glcm_MaximumProbability | 0.598987 | $2.703929 \times 10^{-7}$ | $7.236814 \times 10^{-6}$ | True |
| putamen_gldm_DependenceEntropy | -0.597927 | $2.873718 \times 10^{-7}$ | $7.236814 \times 10^{-6}$ | True |
| putamen_firstorder_90Percentile | 0.595077 | $3.381689 \times 10^{-7}$ | $7.236814 \times 10^{-6}$ | True |
| putamen_firstorder_Uniformity | 0.585441 | $5.795924 \times 10^{-7}$ | $1.033607 \times 10^{-5}$ | True |
| putamen_glcm_JointEnergy | 0.582136 | $6.944448 \times 10^{-7}$ | $1.061508 \times 10^{-5}$ | True |
| putamen_ngtdm_Coarseness | 0.573837 | $1.084068 \times 10^{-6}$ | $1.449941 \times 10^{-5}$ | True |
| putamen_firstorder_10Percentile | 0.539757 | $5.975601 \times 10^{-6}$ | $7.104325 \times 10^{-5}$ | True |
| putamen_glrlm_GrayLevelNonUniformity | -0.528228 | $1.021580 \times 10^{-5}$ | $1.093091 \times 10^{-4}$ | True |

#### 2.5 *Insula*

TABLE V
SPEARMAN CORRELATION RESULTS FOR SIGNIFICANT INSULA RADIOMIC FEATURES.

| Radiomic feature | Spearman $\rho$ | $p$-value | FDR-corrected $p$ | Significant |
|---|---|---|---|---|
| insula_glrlm_RunEntropy | -0.559357 | 0.000002 | 0.000245 | True |
| insula_firstorder_RootMeanSquared | -0.506459 | 0.000027 | 0.001429 | True |
| insula_firstorder_Median | 0.492862 | 0.000047 | 0.001680 | True |
| insula_glrlm_GrayLevelVariance | -0.481132 | 0.000075 | 0.002018 | True |
| insula_firstorder_Energy | -0.474321 | 0.000098 | 0.002105 | True |
| insula_firstorder_TotalEnergy | -0.466955 | 0.000130 | 0.002323 | True |
| insula_firstorder_Mean | 0.454620 | 0.000206 | 0.003144 | True |
| insula_firstorder_Minimum | 0.448464 | 0.000257 | 0.003433 | True |
| insula_glcm_Idn | -0.443394 | 0.000307 | 0.003650 | True |
| insula_glcm_Idmn | -0.429116 | 0.000501 | 0.005363 | True |

#### 2.6 *Heschl's Gyrus*

TABLE VI
SPEARMAN CORRELATION RESULTS FOR SIGNIFICANT HESCHL'S GYRUS RADIOMIC FEATURES.

| Radiomic feature | Spearman $\rho$ | $p$-value | FDR-corrected $p$ | Significant |
|---|---|---|---|---|
| heschls_gyrus_firstorder_Minimum | 0.529364 | 0.000010 | 0.001038 | True |
| heschls_gyrus_firstorder_10Percentile | 0.430756 | 0.000474 | 0.016489 | True |
| heschls_gyrus_glcm_Idmn | -0.418774 | 0.000705 | 0.016489 | True |
| heschls_gyrus_firstorder_Kurtosis | -0.418042 | 0.000722 | 0.016489 | True |
| heschls_gyrus_firstorder_Range | -0.411105 | 0.000903 | 0.016489 | True |
| heschls_gyrus_ngtdm_Contrast | 0.410348 | 0.000925 | 0.016489 | True |
| heschls_gyrus_glrlm_RunEntropy | -0.403790 | 0.001136 | 0.017362 | True |
| heschls_gyrus_glcm_Idn | -0.389108 | 0.001774 | 0.022615 | True |
| heschls_gyrus_gldm_DependenceEntropy | -0.385047 | 0.002000 | 0.022615 | True |
| heschls_gyrus_glrlm_LRHGLE | -0.383155 | 0.002114 | 0.022615 | True |
| heschls_gyrus_glrlm_LRHGLE | -0.378917 | 0.002389 | 0.023243 | True |

##### 2.7 *Entorhinal Cortex*

TABLE VII
SPEARMAN CORRELATION RESULTS FOR ENTORHINAL CORTEX RADIOMIC FEATURES.

| Radiomic feature | Spearman $\rho$ | $p$-value | FDR-corrected $p$ | Significant |
|---|---|---|---|---|
| entorhinal_cortex_glcm_ClusterShade | -0.491550 | 0.000050 | 0.005317 | True |
| entorhinal_cortex_firstorder_Skewness | -0.434010 | 0.000425 | 0.022102 | True |
| entorhinal_cortex_shape_SurfaceVolumeRatio | 0.422734 | 0.000620 | 0.022102 | True |
| entorhinal_cortex_firstorder_Median | 0.405253 | 0.001085 | 0.029031 | True |
| entorhinal_cortex_glrlm_SRLGLE | 0.367944 | 0.003259 | 0.069185 | False |
| entorhinal_cortex_gldm_DependenceNonUniformity | -0.351723 | 0.005058 | 0.069185 | False |
| entorhinal_cortex_gldm_DependenceEntropy | -0.349377 | 0.005381 | 0.069185 | False |
| entorhinal_cortex_shape_VoxelVolume | -0.346746 | 0.005764 | 0.069185 | False |
| entorhinal_cortex_shape_MeshVolume | -0.346376 | 0.005819 | 0.069185 | False |
| entorhinal_cortex_glrlm_GrayLevelNonUniformity | -0.339943 | 0.006866 | 0.073470 | False |

##### 2.8 *Hippocampus*

TABLE VIII
SPEARMAN CORRELATION RESULTS FOR SIGNIFICANT HIPPOCAMPUS RADIOMIC FEATURES.

| Radiomic feature | Spearman $\rho$ | $p$-value | FDR-corrected $p$ | Significant |
|---|---|---|---|---|
| hippocampus_glcm_ClusterShade | -0.603553 | $2.074498 \times 10^{-7}$ | 0.000022 | True |
| hippocampus_gldm_GrayLevelNonUniformity | -0.530398 | $9.249045 \times 10^{-6}$ | 0.000385 | True |
| hippocampus_firstorder_Uniformity | -0.527043 | $1.078309 \times 10^{-5}$ | 0.000385 | True |
| hippocampus_glcm_ClusterTendency | 0.517482 | $1.655091 \times 10^{-5}$ | 0.000443 | True |
| hippocampus_glcm_MCC | 0.506963 | $2.613262 \times 10^{-5}$ | 0.000559 | True |
| hippocampus_glcm_SumSquares | 0.500732 | $3.401180 \times 10^{-5}$ | 0.000607 | True |
| hippocampus_gldm_GrayLevelVariance | 0.495359 | $4.251577 \times 10^{-5}$ | 0.000650 | True |
| hippocampus_firstorder_Entropy | 0.472732 | $1.045708 \times 10^{-4}$ | 0.001260 | True |
| hippocampus_firstorder_Variance | 0.470007 | $1.160516 \times 10^{-4}$ | 0.001260 | True |
| hippocampus_glcm_SumEntropy | 0.469629 | $1.177347 \times 10^{-4}$ | 0.001260 | True |

##### 2.9 *Thalamus*

TABLE IX
SPEARMAN CORRELATION RESULTS FOR SIGNIFICANT THALAMUS RADIOMIC FEATURES.

| Radiomic feature | Spearman $\rho$ | $p$-value | FDR-corrected $p$ | Significant |
|---|---|---|---|---|
| thalamus_glrlm_GrayLevelNonUniformityNormalized | -0.666920 | $3.254824 \times 10^{-9}$ | $3.482662 \times 10^{-7}$ | True |
| thalamus_glrlm_GrayLevelVariance | 0.652390 | $9.198022 \times 10^{-9}$ | $4.920942 \times 10^{-7}$ | True |
| thalamus_glcm_ClusterProminence | 0.594219 | $3.550338 \times 10^{-7}$ | $1.266287 \times 10^{-5}$ | True |
| thalamus_firstorder_Variance | 0.573231 | $1.119333 \times 10^{-6}$ | $2.994215 \times 10^{-5}$ | True |
| thalamus_firstorder_RootMeanSquared | 0.550629 | $3.537717 \times 10^{-6}$ | $7.570715 \times 10^{-5}$ | True |
| thalamus_firstorder_90Percentile | 0.507114 | $2.596421 \times 10^{-5}$ | $4.509244 \times 10^{-4}$ | True |
| thalamus_firstorder_Median | 0.504113 | $2.949972 \times 10^{-5}$ | $4.509244 \times 10^{-4}$ | True |
| thalamus_firstorder_Mean | 0.490995 | $5.082582 \times 10^{-5}$ | $6.774283 \times 10^{-4}$ | True |
| thalamus_firstorder_TotalEnergy | 0.488170 | $5.697995 \times 10^{-5}$ | $6.774283 \times 10^{-4}$ | True |
| thalamus_firstorder_Energy | 0.481359 | $7.474984 \times 10^{-5}$ | $7.998233 \times 10^{-4}$ | True |

##### 2.10 *Vermis*

TABLE X
SPEARMAN CORRELATION RESULTS FOR SIGNIFICANT VERMIS RADIOMIC FEATURES.

| Radiomic feature | Spearman $\rho$ | $p$-value | FDR-corrected $p$ | Significant |
|---|---|---|---|---|
| vermis_shape_Elongation | -0.475254 | 0.000095 | 0.007598 | True |
| vermis_glcm_Imc1 | -0.464659 | 0.000142 | 0.007598 | True |
| vermis_glcm_Imc2 | 0.441502 | 0.000328 | 0.007961 | True |
| vermis_glcm_MCC | 0.441477 | 0.000328 | 0.007961 | True |
| vermis_firstorder_Kurtosis | -0.435019 | 0.000410 | 0.007961 | True |
| vermis_glcm_Correlation | 0.432547 | 0.000446 | 0.007961 | True |
| vermis_shape_MinorAxisLength | -0.415772 | 0.000777 | 0.011884 | True |
| vermis_glrlm_LongRunLowGrayLevelEmphasis | 0.390521 | 0.001701 | 0.021028 | True |
| vermis_glrlm_RunEntropy | 0.389209 | 0.001769 | 0.021028 | True |
| vermis_gldm_DependenceVariance | 0.375310 | 0.002649 | 0.028346 | True |

### 3 Complete Welch t-test Results

Complete Welch t-test results for Alzheimer's disease comparisons are provided in this section. The $t$-value column reports the Welch test statistic. The $n$ normal column gives the number of normal-group samples within the corresponding age bin, and the $n$ AD column gives the number of Alzheimer's disease samples within the corresponding age bin. Reported $p$-values include the raw Welch test $p$-value and the FDR-adjusted $p$-value.

#### 3.1 *Thalamus Welch t-test Results*

TABLE XI: Complete Welch t-test results for thalamus radiomic features comparing normal and Alzheimer's disease age bins.

| Feature | Normal bin | AD bin | $t$-value | $p$-value | $n$ normal | $n$ AD | FDR $p$ | Significant FDR |
|---|---|---|---|---|---|---|---|---|
| **Thalamus** | | | | | | | | |
| glrlm_GrayLevelNonUniformityNormalized | 60–69 | 60–69 | 1.583472 | 0.129187 | 10 | 7 | 0.422794 | False |
| | 70–79 | 70–79 | 1.679039 | 0.104990 | 27 | 18 | 0.413239 | False |
| | 80–89 | 80–89 | 0.005600 | 0.995700 | 25 | 29 | 0.995700 | False |
| glrlm_GrayLevelVariance | 60–69 | 60–69 | -1.788166 | 0.087191 | 10 | 7 | 0.413239 | False |
| | 70–79 | 70–79 | -0.776850 | 0.422558 | 27 | 18 | 0.842316 | False |
| | 80–89 | 80–89 | 0.326078 | 0.749725 | 25 | 29 | 0.902746 | False |
| glcm_ClusterProminence | 60–69 | 60–69 | -1.792820 | 0.045395 | 10 | 7 | 0.413239 | False |
| | 70–79 | 70–79 | -1.148572 | 0.238776 | 27 | 18 | 0.538252 | False |
| | 80–89 | 80–89 | -0.411033 | 0.699730 | 25 | 29 | 0.902746 | False |
| firstorder_Variance | 60–69 | 60–69 | -1.271722 | 0.192581 | 10 | 7 | 0.533301 | False |
| | 70–79 | 70–79 | -1.130333 | 0.251975 | 27 | 18 | 0.538252 | False |
| | 80–89 | 80–89 | -0.653138 | 0.520748 | 25 | 29 | 0.842316 | False |
| firstorder_RootMeanSquared | 60–69 | 60–69 | -1.885506 | 0.054395 | 10 | 7 | 0.413239 | False |
| | 70–79 | 70–79 | 0.270711 | 0.792921 | 27 | 18 | 0.902746 | False |
| | 80–89 | 80–89 | -0.685016 | 0.502950 | 25 | 29 | 0.842316 | False |
| firstorder_90Percentile | 60–69 | 60–69 | -1.324692 | 0.189381 | 10 | 7 | 0.533301 | False |
| | 70–79 | 70–79 | 0.227143 | 0.827517 | 27 | 18 | 0.902746 | False |
| | 80–89 | 80–89 | -0.418489 | 0.691931 | 25 | 29 | 0.902746 | False |
| firstorder_Median | 60–69 | 60–69 | -1.681248 | 0.100590 | 10 | 7 | 0.413239 | False |
| | 70–79 | 70–79 | 0.288020 | 0.761924 | 27 | 18 | 0.902746 | False |
| | 80–89 | 80–89 | -0.637453 | 0.513149 | 25 | 29 | 0.842316 | False |
| firstorder_Mean | 60–69 | 60–69 | -1.541050 | 0.114789 | 10 | 7 | 0.413239 | False |
| | 70–79 | 70–79 | 0.372954 | 0.703130 | 27 | 18 | 0.902746 | False |
| | 80–89 | 80–89 | -0.458102 | 0.665533 | 25 | 29 | 0.902746 | False |
| firstorder_TotalEnergy | 60–69 | 60–69 | -1.603198 | 0.077992 | 10 | 7 | 0.413239 | False |
| | 70–79 | 70–79 | 0.252906 | 0.813719 | 27 | 18 | 0.902746 | False |
| | 80–89 | 80–89 | -0.003666 | 0.978102 | 25 | 29 | 0.995700 | False |
| firstorder_Energy | 60–69 | 60–69 | -1.603198 | 0.089391 | 10 | 7 | 0.413239 | False |
| | 70–79 | 70–79 | 0.263543 | 0.805919 | 27 | 18 | 0.902746 | False |
| | 80–89 | 80–89 | 0.028263 | 0.977302 | 25 | 29 | 0.995700 | False |

#### 3.2 *Cingulum Welch t-test Results*

TABLE XII: Complete Welch t-test results for cingulum radiomic features comparing normal and Alzheimer's disease age bins.

| Feature | Normal bin | AD bin | $t$-value | $p$-value | $n$ normal | $n$ AD | FDR $p$ | Significant FDR |
|---|---|---|---|---|---|---|---|---|
| firstorder_Median | 60–69 | 60–69 | -1.425608 | 0.156384 | 10 | 7 | 0.234577 | False |
| | 70–79 | 70–79 | -1.844651 | 0.069393 | 27 | 18 | 0.140786 | False |
| | 80–89 | 80–89 | -0.904011 | 0.374563 | 25 | 29 | 0.488560 | False |
| firstorder_Kurtosis | 60–69 | 60–69 | 3.086475 | 0.007599 | 10 | 7 | 0.025331 | True |
| | 70–79 | 70–79 | 3.108392 | 0.004000 | 27 | 18 | 0.014999 | True |
| | 80–89 | 80–89 | 5.750663 | 0.000200 | 25 | 29 | 0.005999 | True |
| firstorder_InterquartileRange | 60–69 | 60–69 | -1.914115 | 0.070393 | 10 | 7 | 0.140786 | False |
| | 70–79 | 70–79 | -2.986220 | 0.003200 | 27 | 18 | 0.013713 | True |
| | 80–89 | 80–89 | -3.915657 | 0.000600 | 25 | 29 | 0.008999 | True |
| firstorder_RobustMeanAbsoluteDeviation | 60–69 | 60–69 | -1.680061 | 0.113989 | 10 | 7 | 0.189139 | False |
| | 70–79 | 70–79 | -2.661045 | 0.009799 | 27 | 18 | 0.029397 | True |
| | 80–89 | 80–89 | -3.414790 | 0.001000 | 25 | 29 | 0.008999 | True |
| firstorder_Mean | 60–69 | 60–69 | -0.825177 | 0.414559 | 10 | 7 | 0.518198 | False |
| | 70–79 | 70–79 | -1.335462 | 0.181382 | 27 | 18 | 0.259117 | False |
| | 80–89 | 80–89 | -0.450874 | 0.663134 | 25 | 29 | 0.734712 | False |
| glszm_GrayLevelNonUniformityNormalized | 60–69 | 60–69 | -2.343420 | 0.013199 | 10 | 7 | 0.035996 | True |
| | 70–79 | 70–79 | -2.109358 | 0.019398 | 27 | 18 | 0.048495 | True |
| | 80–89 | 80–89 | -3.364285 | 0.002200 | 25 | 29 | 0.010999 | True |

Continued on next page

TABLE XII – continued from previous page

| Feature | Normal bin | AD bin | $t$-value | $p$-value | $n$ normal | $n$ AD | FDR $p$ | Significant FDR |
|---|---|---|---|---|---|---|---|---|
| firstorder_Minimum | 60–69 | 60–69 | -3.376632 | 0.001200 | 10 | 7 | 0.008999 | True |
| | 70–79 | 70–79 | -2.187042 | 0.032797 | 27 | 18 | 0.075685 | False |
| | 80–89 | 80–89 | -3.313819 | 0.002000 | 25 | 29 | 0.010999 | True |
| firstorder_90Percentile | 60–69 | 60–69 | -0.278109 | 0.780922 | 10 | 7 | 0.807850 | False |
| | 70–79 | 70–79 | -0.976957 | 0.322368 | 27 | 18 | 0.439592 | False |
| | 80–89 | 80–89 | -0.423012 | 0.685731 | 25 | 29 | 0.734712 | False |
| firstorder_MeanAbsoluteDeviation | 60–69 | 60–69 | -0.752869 | 0.450755 | 10 | 7 | 0.540906 | False |
| | 70–79 | 70–79 | -1.600568 | 0.119788 | 27 | 18 | 0.189139 | False |
| | 80–89 | 80–89 | -1.708793 | 0.095590 | 25 | 29 | 0.168689 | False |
| firstorder_Energy | 60–69 | 60–69 | 2.088036 | 0.080592 | 10 | 7 | 0.151110 | False |
| | 70–79 | 70–79 | 0.210697 | 0.828517 | 27 | 18 | 0.828517 | False |
| | 80–89 | 80–89 | 0.661141 | 0.521548 | 25 | 29 | 0.601786 | False |

#### *3.3 Heschl's Gyrus Welch t-test Results*

TABLE XIII: Complete Welch t-test results for Heschl's gyrus radiomic features comparing normal and Alzheimer's disease age bins.

| Feature | Normal bin | AD bin | $t$-value | $p$-value | $n$ normal | $n$ AD | FDR $p$ | Significant FDR |
|---|---|---|---|---|---|---|---|---|
| firstorder_Minimum | 60–69 | 60–69 | -3.597101 | 0.003000 | 10 | 7 | 0.022998 | True |
| | 70–79 | 70–79 | -2.729869 | 0.010399 | 27 | 18 | 0.034663 | True |
| | 80–89 | 80–89 | -3.275752 | 0.002600 | 25 | 29 | 0.022998 | True |
| firstorder_10Percentile | 60–69 | 60–69 | -1.310041 | 0.196780 | 10 | 7 | 0.295170 | False |
| | 70–79 | 70–79 | -2.044255 | 0.038796 | 27 | 18 | 0.089530 | False |
| | 80–89 | 80–89 | -1.047618 | 0.305369 | 25 | 29 | 0.381712 | False |
| glcm_Idmn | 60–69 | 60–69 | 2.401641 | 0.005599 | 10 | 7 | 0.023998 | True |
| | 70–79 | 70–79 | 1.145607 | 0.255374 | 27 | 18 | 0.348238 | False |
| | 80–89 | 80–89 | 0.377748 | 0.698130 | 25 | 29 | 0.747997 | False |
| firstorder_Kurtosis | 60–69 | 60–69 | 2.049401 | 0.066593 | 10 | 7 | 0.132362 | False |
| | 70–79 | 70–79 | 1.075156 | 0.279372 | 27 | 18 | 0.364398 | False |
| | 80–89 | 80–89 | 3.451963 | 0.000800 | 25 | 29 | 0.022998 | True |
| firstorder_Range | 60–69 | 60–69 | 3.549507 | 0.004600 | 10 | 7 | 0.022998 | True |
| | 70–79 | 70–79 | 2.851945 | 0.006999 | 27 | 18 | 0.026247 | True |
| | 80–89 | 80–89 | 2.545642 | 0.014399 | 25 | 29 | 0.039269 | True |
| ngtdm_Contrast | 60–69 | 60–69 | -2.252562 | 0.004000 | 10 | 7 | 0.022998 | True |
| | 70–79 | 70–79 | -0.451461 | 0.650535 | 27 | 18 | 0.722817 | False |
| | 80–89 | 80–89 | -0.328310 | 0.737526 | 25 | 29 | 0.762958 | False |
| glrlm_RunEntropy | 60–69 | 60–69 | 2.207065 | 0.047395 | 10 | 7 | 0.101561 | False |
| | 70–79 | 70–79 | 1.786474 | 0.070593 | 27 | 18 | 0.132362 | False |
| | 80–89 | 80–89 | 1.622222 | 0.117388 | 25 | 29 | 0.188823 | False |
| glcm_Idn | 60–69 | 60–69 | 2.239171 | 0.012799 | 10 | 7 | 0.038396 | True |
| | 70–79 | 70–79 | 0.954904 | 0.348565 | 27 | 18 | 0.411882 | False |
| | 80–89 | 80–89 | 0.204592 | 0.828117 | 25 | 29 | 0.828117 | False |
| gldm_DependenceEntropy | 60–69 | 60–69 | 1.207280 | 0.247975 | 10 | 7 | 0.348238 | False |
| | 70–79 | 70–79 | 1.588575 | 0.119588 | 27 | 18 | 0.188823 | False |
| | 80–89 | 80–89 | 2.093589 | 0.036596 | 25 | 29 | 0.089530 | False |
| glrlm_LongRunHighGrayLevelEmphasis | 60–69 | 60–69 | 3.553918 | 0.004600 | 10 | 7 | 0.022998 | True |
| | 70–79 | 70–79 | 0.964040 | 0.356964 | 27 | 18 | 0.411882 | False |
| | 80–89 | 80–89 | 1.703002 | 0.093991 | 25 | 29 | 0.165866 | False |

#### *3.4 Vermis Welch t-test Results*

TABLE XIV: Complete Welch t-test results for vermis radiomic features comparing normal and Alzheimer's disease age bins.

| Feature | Normal bin | AD bin | $t$-value | $p$-value | $n$ normal | $n$ AD | FDR $p$ | Significant FDR |
|---|---|---|---|---|---|---|---|---|
| shape_Elongation | 60–69 | 60–69 | 0.388666 | 0.702930 | 10 | 7 | 0.781033 | False |
| | 70–79 | 70–79 | 2.122347 | 0.047195 | 27 | 18 | 0.088866 | False |
| | 80–89 | 80–89 | -1.020315 | 0.310569 | 25 | 29 | 0.388211 | False |
| glcm_Imc1 | 60–69 | 60–69 | 2.746783 | 0.008999 | 10 | 7 | 0.064660 | False |
| | 70–79 | 70–79 | 2.691628 | 0.005599 | 27 | 18 | 0.064660 | False |
| | 80–89 | 80–89 | 2.347179 | 0.021598 | 25 | 29 | 0.064794 | False |
| glcm_Imc2 | 60–69 | 60–69 | -2.272881 | 0.033597 | 10 | 7 | 0.083069 | False |
| | 70–79 | 70–79 | -2.313923 | 0.017198 | 27 | 18 | 0.064660 | False |
| | 80–89 | 80–89 | -1.728058 | 0.093791 | 25 | 29 | 0.156318 | False |
| glcm_MCC | 60–69 | 60–69 | -2.078978 | 0.047395 | 10 | 7 | 0.088866 | False |
| | 70–79 | 70–79 | -2.399473 | 0.013999 | 27 | 18 | 0.064660 | False |

Continued on next page

TABLE XIV – continued from previous page

| Feature | Normal bin | AD bin | $t$-value | $p$-value | $n$ normal | $n$ AD | FDR $p$ | Significant FDR |
|---|---|---|---|---|---|---|---|---|
| | 80–89 | 80–89 | -1.469887 | 0.142186 | 25 | 29 | 0.203123 | False |
| firstorder_Kurtosis | 60–69 | 60–69 | 0.794825 | 0.441756 | 10 | 7 | 0.530107 | False |
| | 70–79 | 70–79 | -0.167880 | 0.877712 | 27 | 18 | 0.919108 | False |
| | 80–89 | 80–89 | 0.128377 | 0.897310 | 25 | 29 | 0.919108 | False |
| glcm_Correlation | 60–69 | 60–69 | -2.249399 | 0.035996 | 10 | 7 | 0.083069 | False |
| | 70–79 | 70–79 | -2.489235 | 0.012599 | 27 | 18 | 0.064660 | False |
| | 80–89 | 80–89 | -1.991234 | 0.050395 | 25 | 29 | 0.088932 | False |
| shape_MinorAxisLength | 60–69 | 60–69 | 0.134912 | 0.919108 | 10 | 7 | 0.919108 | False |
| | 70–79 | 70–79 | 1.064819 | 0.305969 | 27 | 18 | 0.388211 | False |
| | 80–89 | 80–89 | 0.479909 | 0.635136 | 25 | 29 | 0.732850 | False |
| glrlm_LongRunLowGrayLevelEmphasis | 60–69 | 60–69 | -2.400215 | 0.019398 | 10 | 7 | 0.064660 | False |
| | 70–79 | 70–79 | -3.347341 | 0.002400 | 27 | 18 | 0.064660 | False |
| | 80–89 | 80–89 | -2.333989 | 0.018398 | 25 | 29 | 0.064660 | False |
| glrlm_RunEntropy | 60–69 | 60–69 | -1.721288 | 0.105589 | 10 | 7 | 0.159884 | False |
| | 70–79 | 70–79 | -2.177749 | 0.028197 | 27 | 18 | 0.076901 | False |
| | 80–89 | 80–89 | -1.631359 | 0.106589 | 25 | 29 | 0.159884 | False |
| gldm_DependenceVariance | 60–69 | 60–69 | -3.078174 | 0.006799 | 10 | 7 | 0.064660 | False |
| | 70–79 | 70–79 | -2.007745 | 0.039796 | 27 | 18 | 0.085277 | False |
| | 80–89 | 80–89 | -1.173299 | 0.246575 | 25 | 29 | 0.336239 | False |

#### 3.5 *Putamen Welch t-test Results*

TABLE XV: Complete Welch t-test results for putamen radiomic features comparing normal and Alzheimer's disease age bins.

| Feature | Normal bin | AD bin | $t$-value | $p$-value | $n$ normal | $n$ AD | FDR $p$ | Significant FDR |
|---|---|---|---|---|---|---|---|---|
| firstorder_Median | 60–69 | 60–69 | -1.306587 | 0.199380 | 10 | 7 | 0.260061 | False |
| | 70–79 | 70–79 | -1.937186 | 0.066793 | 27 | 18 | 0.182164 | False |
| | 80–89 | 80–89 | -1.375933 | 0.177582 | 25 | 29 | 0.260061 | False |
| firstorder_Mean | 60–69 | 60–69 | -1.296500 | 0.195180 | 10 | 7 | 0.260061 | False |
| | 70–79 | 70–79 | -1.878445 | 0.076192 | 27 | 18 | 0.190481 | False |
| | 80–89 | 80–89 | -1.353985 | 0.190581 | 25 | 29 | 0.260061 | False |
| glcm_MaximumProbability | 60–69 | 60–69 | -2.656421 | 0.018398 | 10 | 7 | 0.127987 | False |
| | 70–79 | 70–79 | -1.387445 | 0.182782 | 27 | 18 | 0.260061 | False |
| | 80–89 | 80–89 | -0.905720 | 0.369963 | 25 | 29 | 0.426880 | False |
| gldm_DependenceEntropy | 60–69 | 60–69 | 2.504524 | 0.022998 | 10 | 7 | 0.127987 | False |
| | 70–79 | 70–79 | 1.634556 | 0.120988 | 27 | 18 | 0.226852 | False |
| | 80–89 | 80–89 | 0.744398 | 0.450155 | 25 | 29 | 0.482309 | False |
| firstorder_90Percentile | 60–69 | 60–69 | -0.200216 | 0.866713 | 10 | 7 | 0.866713 | False |
| | 70–79 | 70–79 | -2.038969 | 0.049595 | 27 | 18 | 0.161384 | False |
| | 80–89 | 80–89 | -2.005953 | 0.053795 | 25 | 29 | 0.161384 | False |
| firstorder_Uniformity | 60–69 | 60–69 | -2.454724 | 0.017598 | 10 | 7 | 0.127987 | False |
| | 70–79 | 70–79 | -1.597688 | 0.113789 | 27 | 18 | 0.226852 | False |
| | 80–89 | 80–89 | -0.883988 | 0.384362 | 25 | 29 | 0.427068 | False |
| glcm_JointEnergy | 60–69 | 60–69 | -2.500774 | 0.015198 | 10 | 7 | 0.127987 | False |
| | 70–79 | 70–79 | -1.635341 | 0.103790 | 27 | 18 | 0.222406 | False |
| | 80–89 | 80–89 | -1.024979 | 0.311369 | 25 | 29 | 0.373643 | False |
| ngtdm_Coarseness | 60–69 | 60–69 | -2.099913 | 0.025597 | 10 | 7 | 0.127987 | False |
| | 70–79 | 70–79 | -2.205414 | 0.014399 | 27 | 18 | 0.127987 | False |
| | 80–89 | 80–89 | -1.407542 | 0.177382 | 25 | 29 | 0.260061 | False |
| firstorder_10Percentile | 60–69 | 60–69 | -2.304544 | 0.036196 | 10 | 7 | 0.155127 | False |
| | 70–79 | 70–79 | -1.461897 | 0.165383 | 27 | 18 | 0.260061 | False |
| | 80–89 | 80–89 | -0.261347 | 0.799720 | 25 | 29 | 0.827297 | False |
| glrlm_GrayLevelNonUniformity | 60–69 | 60–69 | 1.870628 | 0.084192 | 10 | 7 | 0.194288 | False |
| | 70–79 | 70–79 | 2.043896 | 0.044996 | 27 | 18 | 0.161384 | False |
| | 80–89 | 80–89 | 1.062418 | 0.299570 | 25 | 29 | 0.373643 | False |

#### 3.6 *Insula Welch t-test Results*

TABLE XVI: Complete Welch t-test results for insula radiomic features comparing normal and Alzheimer's disease age bins.

| Feature | Normal bin | AD bin | $t$-value | $p$-value | $n$ normal | $n$ AD | FDR $p$ | Significant FDR |
|---|---|---|---|---|---|---|---|---|
| glrlm_RunEntropy | 60–69 | 60–69 | 1.890836 | 0.076992 | 10 | 7 | 0.164984 | False |
| | 70–79 | 70–79 | 3.139198 | 0.001800 | 27 | 18 | 0.017998 | True |
| | 80–89 | 80–89 | 2.176860 | 0.035996 | 25 | 29 | 0.098172 | False |
| firstorder_RootMeanSquared | 60–69 | 60–69 | 0.173356 | 0.875912 | 10 | 7 | 0.906116 | False |

Continued on next page

TABLE XVI – continued from previous page

| Feature | Normal bin | AD bin | $t$-value | $p$-value | $n$ normal | $n$ AD | FDR $p$ | Significant FDR |
|---|---|---|---|---|---|---|---|---|
| | 70–79 | 70–79 | 0.877272 | 0.383162 | 27 | 18 | 0.463634 | False |
| | 80–89 | 80–89 | -0.695272 | 0.491151 | 25 | 29 | 0.566713 | False |
| firstorder_Median | 60–69 | 60–69 | -0.272803 | 0.786321 | 10 | 7 | 0.873690 | False |
| | 70–79 | 70–79 | -1.969407 | 0.055994 | 27 | 18 | 0.139986 | False |
| | 80–89 | 80–89 | -2.277094 | 0.027197 | 25 | 29 | 0.081592 | False |
| glrlm_GrayLevelVariance | 60–69 | 60–69 | 2.841914 | 0.009199 | 10 | 7 | 0.034497 | True |
| | 70–79 | 70–79 | 1.380269 | 0.186581 | 27 | 18 | 0.373163 | False |
| | 80–89 | 80–89 | -0.101726 | 0.914909 | 25 | 29 | 0.914909 | False |
| firstorder_Energy | 60–69 | 60–69 | 0.973862 | 0.361564 | 10 | 7 | 0.463634 | False |
| | 70–79 | 70–79 | 0.971684 | 0.322368 | 27 | 18 | 0.463634 | False |
| | 80–89 | 80–89 | 1.042189 | 0.308769 | 25 | 29 | 0.463634 | False |
| firstorder_TotalEnergy | 60–69 | 60–69 | 0.973862 | 0.372363 | 10 | 7 | 0.463634 | False |
| | 70–79 | 70–79 | 0.875201 | 0.386361 | 27 | 18 | 0.463634 | False |
| | 80–89 | 80–89 | 1.012559 | 0.312769 | 25 | 29 | 0.463634 | False |
| firstorder_Mean | 60–69 | 60–69 | 0.191273 | 0.824718 | 10 | 7 | 0.883626 | False |
| | 70–79 | 70–79 | -1.252604 | 0.217178 | 27 | 18 | 0.407209 | False |
| | 80–89 | 80–89 | -0.992322 | 0.333567 | 25 | 29 | 0.463634 | False |
| firstorder_Minimum | 60–69 | 60–69 | -3.155577 | 0.007999 | 10 | 7 | 0.034282 | True |
| | 70–79 | 70–79 | -2.330420 | 0.022798 | 27 | 18 | 0.075992 | False |
| | 80–89 | 80–89 | -2.853254 | 0.005799 | 25 | 29 | 0.029997 | True |
| glcm_Idn | 60–69 | 60–69 | 0.964021 | 0.339766 | 10 | 7 | 0.463634 | False |
| | 70–79 | 70–79 | 4.003342 | 0.000600 | 27 | 18 | 0.008999 | True |
| | 80–89 | 80–89 | 3.225434 | 0.002400 | 25 | 29 | 0.017998 | True |
| glcm_Idmn | 60–69 | 60–69 | 1.959901 | 0.063194 | 10 | 7 | 0.145832 | False |
| | 70–79 | 70–79 | 3.695942 | 0.000200 | 27 | 18 | 0.005999 | True |
| | 80–89 | 80–89 | 2.902081 | 0.005999 | 25 | 29 | 0.029997 | True |

#### *3.7 Caudate Nucleus Welch t-test Results*

TABLE XVII: Complete Welch t-test results for caudate nucleus radiomic features comparing normal and Alzheimer's disease age bins.

| Feature | Normal bin | AD bin | $t$-value | $p$-value | $n$ normal | $n$ AD | FDR $p$ | Significant FDR |
|---|---|---|---|---|---|---|---|---|
| firstorder_Uniformity | 60–69 | 60–69 | 0.686713 | 0.523748 | 10 | 7 | 0.716656 | False |
| | 70–79 | 70–79 | 1.560411 | 0.140186 | 27 | 18 | 0.262849 | False |
| | 80–89 | 80–89 | -2.865272 | 0.006799 | 25 | 29 | 0.040796 | True |
| glcm_SumSquares | 60–69 | 60–69 | -0.475155 | 0.634737 | 10 | 7 | 0.761684 | False |
| | 70–79 | 70–79 | -1.379254 | 0.183582 | 27 | 18 | 0.323968 | False |
| | 80–89 | 80–89 | 2.604860 | 0.009599 | 25 | 29 | 0.047995 | True |
| glrlm_RunVariance | 60–69 | 60–69 | 0.285059 | 0.810719 | 10 | 7 | 0.900799 | False |
| | 70–79 | 70–79 | 2.012709 | 0.057394 | 27 | 18 | 0.132910 | False |
| | 80–89 | 80–89 | -3.277358 | 0.003200 | 25 | 29 | 0.040496 | True |
| glcm_JointEnergy | 60–69 | 60–69 | 0.706637 | 0.525547 | 10 | 7 | 0.716656 | False |
| | 70–79 | 70–79 | 1.632875 | 0.120988 | 27 | 18 | 0.257974 | False |
| | 80–89 | 80–89 | -3.316926 | 0.002400 | 25 | 29 | 0.040496 | True |
| gldm_LargeDependenceEmphasis | 60–69 | 60–69 | 0.518004 | 0.632937 | 10 | 7 | 0.761684 | False |
| | 70–79 | 70–79 | 2.019775 | 0.055994 | 27 | 18 | 0.132910 | False |
| | 80–89 | 80–89 | -2.774295 | 0.005399 | 25 | 29 | 0.040496 | True |
| glrlm_RunPercentage | 60–69 | 60–69 | -0.424800 | 0.687131 | 10 | 7 | 0.792844 | False |
| | 70–79 | 70–79 | -2.004135 | 0.057594 | 27 | 18 | 0.132910 | False |
| | 80–89 | 80–89 | 2.565092 | 0.014399 | 25 | 29 | 0.056661 | False |
| glcm_ClusterTendency | 60–69 | 60–69 | -0.606589 | 0.584942 | 10 | 7 | 0.761684 | False |
| | 70–79 | 70–79 | -1.209716 | 0.241176 | 27 | 18 | 0.401960 | False |
| | 80–89 | 80–89 | 2.407392 | 0.015198 | 25 | 29 | 0.056661 | False |
| firstorder_Median | 60–69 | 60–69 | -2.375851 | 0.016998 | 10 | 7 | 0.056661 | False |
| | 70–79 | 70–79 | -0.733515 | 0.447555 | 27 | 18 | 0.706666 | False |
| | 80–89 | 80–89 | -0.113435 | 0.898310 | 25 | 29 | 0.962475 | False |
| glrlm_LongRunEmphasis | 60–69 | 60–69 | 0.035673 | 0.995900 | 10 | 7 | 0.997900 | False |
| | 70–79 | 70–79 | 2.083223 | 0.054395 | 27 | 18 | 0.132910 | False |
| | 80–89 | 80–89 | -3.022020 | 0.004400 | 25 | 29 | 0.040496 | True |
| firstorder_90Percentile | 60–69 | 60–69 | -1.512745 | 0.128987 | 10 | 7 | 0.257974 | False |
| | 70–79 | 70–79 | -0.676380 | 0.490551 | 27 | 18 | 0.716656 | False |
| | 80–89 | 80–89 | -0.005259 | 0.997900 | 25 | 29 | 0.997900 | False |

#### *3.8 Rolandic Operculum Welch t-test Results*

TABLE XVIII: Complete Welch t-test results for rolandic operculum radiomic features comparing normal and Alzheimer's disease age bins.

| Feature | Normal bin | AD bin | $t$-value | $p$-value | $n$ normal | $n$ AD | FDR $p$ | Significant FDR |
|---|---|---|---|---|---|---|---|---|
| firstorder_Kurtosis | 60–69 | 60–69 | 1.763294 | 0.091591 | 10 | 7 | 0.183182 | False |
| | 70–79 | 70–79 | 1.814892 | 0.076392 | 27 | 18 | 0.176290 | False |
| | 80–89 | 80–89 | 1.899462 | 0.065793 | 25 | 29 | 0.164484 | False |
| ngtdm_Contrast | 60–69 | 60–69 | 0.101206 | 0.930707 | 10 | 7 | 0.944106 | False |
| | 70–79 | 70–79 | -0.367582 | 0.700530 | 27 | 18 | 0.808304 | False |
| | 80–89 | 80–89 | -1.691502 | 0.087991 | 25 | 29 | 0.183182 | False |
| firstorder_Minimum | 60–69 | 60–69 | -2.866857 | 0.015198 | 10 | 7 | 0.073707 | False |
| | 70–79 | 70–79 | -2.560211 | 0.014399 | 27 | 18 | 0.073707 | False |
| | 80–89 | 80–89 | -3.829649 | 0.000600 | 25 | 29 | 0.017998 | True |
| glcm_Idmn | 60–69 | 60–69 | 0.137182 | 0.856514 | 10 | 7 | 0.917694 | False |
| | 70–79 | 70–79 | 0.662250 | 0.517548 | 27 | 18 | 0.621058 | False |
| | 80–89 | 80–89 | 1.647061 | 0.104590 | 25 | 29 | 0.184570 | False |
| firstorder_Range | 60–69 | 60–69 | 2.530075 | 0.017198 | 10 | 7 | 0.073707 | False |
| | 70–79 | 70–79 | 1.397380 | 0.167983 | 27 | 18 | 0.265237 | False |
| | 80–89 | 80–89 | 2.668727 | 0.010199 | 25 | 29 | 0.073707 | False |
| glcm_MaximumProbability | 60–69 | 60–69 | -0.267834 | 0.790921 | 10 | 7 | 0.878801 | False |
| | 70–79 | 70–79 | 1.938857 | 0.062994 | 27 | 18 | 0.164484 | False |
| | 80–89 | 80–89 | 1.941812 | 0.054395 | 25 | 29 | 0.163184 | False |
| glcm_Idn | 60–69 | 60–69 | -0.789772 | 0.448555 | 10 | 7 | 0.568943 | False |
| | 70–79 | 70–79 | 0.831190 | 0.398560 | 27 | 18 | 0.543491 | False |
| | 80–89 | 80–89 | 1.237244 | 0.222178 | 25 | 29 | 0.333267 | False |
| glszm_GrayLevelNonUniformity | 60–69 | 60–69 | -2.540097 | 0.015198 | 10 | 7 | 0.073707 | False |
| | 70–79 | 70–79 | -1.026693 | 0.284772 | 27 | 18 | 0.406816 | False |
| | 80–89 | 80–89 | -0.759364 | 0.455154 | 25 | 29 | 0.568943 | False |
| glszm_ZoneEntropy | 60–69 | 60–69 | 1.584425 | 0.116188 | 10 | 7 | 0.193647 | False |
| | 70–79 | 70–79 | 2.906177 | 0.006999 | 27 | 18 | 0.073707 | False |
| | 80–89 | 80–89 | 1.689540 | 0.104590 | 25 | 29 | 0.184570 | False |
| glrlm_LongRunHighGrayLevelEmphasis | 60–69 | 60–69 | 0.088093 | 0.944106 | 10 | 7 | 0.944106 | False |
| | 70–79 | 70–79 | 2.322182 | 0.023798 | 27 | 18 | 0.089241 | False |
| | 80–89 | 80–89 | 2.004519 | 0.052995 | 25 | 29 | 0.163184 | False |

##### *3.9 Hippocampus Welch t-test Results*

TABLE XIX: Complete Welch t-test results for hippocampus radiomic features comparing normal and Alzheimer's disease age bins.

| Feature | Normal bin | AD bin | $t$-value | $p$-value | $n$ normal | $n$ AD | FDR $p$ | Significant FDR |
|---|---|---|---|---|---|---|---|---|
| glcm_ClusterShade | 60–69 | 60–69 | 2.122118 | 0.046995 | 10 | 7 | 0.061298 | False |
| | 70–79 | 70–79 | 4.045729 | 0.000200 | 27 | 18 | 0.000353 | True |
| | 80–89 | 80–89 | 4.538478 | 0.000200 | 25 | 29 | 0.000353 | True |
| gldm_GrayLevelNonUniformity | 60–69 | 60–69 | 2.633727 | 0.021598 | 10 | 7 | 0.029452 | True |
| | 70–79 | 70–79 | 6.284979 | 0.000200 | 27 | 18 | 0.000353 | True |
| | 80–89 | 80–89 | 6.722051 | 0.000200 | 25 | 29 | 0.000353 | True |
| firstorder_Uniformity | 60–69 | 60–69 | 0.316077 | 0.763524 | 10 | 7 | 0.763524 | False |
| | 70–79 | 70–79 | 4.497279 | 0.000600 | 27 | 18 | 0.000947 | True |
| | 80–89 | 80–89 | 6.267591 | 0.000200 | 25 | 29 | 0.000353 | True |
| glcm_ClusterTendency | 60–69 | 60–69 | -1.433810 | 0.166383 | 10 | 7 | 0.207979 | False |
| | 70–79 | 70–79 | -3.939627 | 0.000200 | 27 | 18 | 0.000353 | True |
| | 80–89 | 80–89 | -5.873615 | 0.000200 | 25 | 29 | 0.000353 | True |
| glcm_MCC | 60–69 | 60–69 | -3.760032 | 0.000200 | 10 | 7 | 0.000353 | True |
| | 70–79 | 70–79 | -2.694331 | 0.008599 | 27 | 18 | 0.012284 | True |
| | 80–89 | 80–89 | -4.780716 | 0.000200 | 25 | 29 | 0.000353 | True |
| glcm_SumSquares | 60–69 | 60–69 | -0.771664 | 0.433357 | 10 | 7 | 0.481507 | False |
| | 70–79 | 70–79 | -4.110778 | 0.000200 | 27 | 18 | 0.000353 | True |
| | 80–89 | 80–89 | -5.544352 | 0.000200 | 25 | 29 | 0.000353 | True |
| gldm_GrayLevelVariance | 60–69 | 60–69 | -0.867169 | 0.391561 | 10 | 7 | 0.451801 | False |
| | 70–79 | 70–79 | -3.992477 | 0.000200 | 27 | 18 | 0.000353 | True |
| | 80–89 | 80–89 | -5.386304 | 0.000200 | 25 | 29 | 0.000353 | True |
| firstorder_Entropy | 60–69 | 60–69 | -0.391257 | 0.703930 | 10 | 7 | 0.728203 | False |
| | 70–79 | 70–79 | -3.950009 | 0.000800 | 27 | 18 | 0.001200 | True |
| | 80–89 | 80–89 | -4.844296 | 0.000200 | 25 | 29 | 0.000353 | True |
| firstorder_Variance | 60–69 | 60–69 | -1.016047 | 0.302970 | 10 | 7 | 0.363564 | False |
| | 70–79 | 70–79 | -3.804961 | 0.000400 | 27 | 18 | 0.000667 | True |
| | 80–89 | 80–89 | -5.183429 | 0.000200 | 25 | 29 | 0.000353 | True |
| glcm_SumEntropy | 60–69 | 60–69 | -0.462720 | 0.643736 | 10 | 7 | 0.689717 | False |
| | 70–79 | 70–79 | -4.126107 | 0.000200 | 27 | 18 | 0.000353 | True |
| | 80–89 | 80–89 | -5.477922 | 0.000200 | 25 | 29 | 0.000353 | True |

#### 3.10 Entorhinal Cortex Welch t-test Results

TABLE XX: Complete Welch t-test results for entorhinal cortex radiomic features comparing normal and Alzheimer's disease age bins.

| Feature | Normal bin | AD bin | $t$-value | $p$-value | $n$ normal | $n$ AD | FDR $p$ | Significant FDR |
|---|---|---|---|---|---|---|---|---|
| glcm_ClusterShade | 60–69 | 60–69 | 2.850014 | 0.009199 | 10 | 7 | 0.013142 | True |
| | 70–79 | 70–79 | 4.641229 | 0.000200 | 27 | 18 | 0.000750 | True |
| | 80–89 | 80–89 | 3.877441 | 0.000400 | 25 | 29 | 0.001091 | True |
| firstorder_Skewness | 60–69 | 60–69 | 2.528000 | 0.028197 | 10 | 7 | 0.035246 | True |
| | 70–79 | 70–79 | 1.247736 | 0.226977 | 27 | 18 | 0.234804 | False |
| | 80–89 | 80–89 | -1.469178 | 0.151985 | 25 | 29 | 0.168872 | False |
| shape_SurfaceVolumeRatio | 60–69 | 60–69 | -2.208078 | 0.034797 | 10 | 7 | 0.041756 | True |
| | 70–79 | 70–79 | -3.726391 | 0.000200 | 27 | 18 | 0.000750 | True |
| | 80–89 | 80–89 | -3.689211 | 0.000400 | 25 | 29 | 0.001091 | True |
| firstorder_Median | 60–69 | 60–69 | -1.544755 | 0.136986 | 10 | 7 | 0.158061 | False |
| | 70–79 | 70–79 | -2.801645 | 0.008399 | 27 | 18 | 0.012599 | True |
| | 80–89 | 80–89 | -2.382114 | 0.019398 | 25 | 29 | 0.025563 | True |
| glrlm_ShortRunLowGrayLevelEmphasis | 60–69 | 60–69 | -2.585210 | 0.019598 | 10 | 7 | 0.025563 | True |
| | 70–79 | 70–79 | -3.310305 | 0.002400 | 27 | 18 | 0.004800 | True |
| | 80–89 | 80–89 | -3.139659 | 0.003000 | 25 | 29 | 0.005294 | True |
| gldm_DependenceNonUniformity | 60–69 | 60–69 | 3.304681 | 0.002800 | 10 | 7 | 0.005249 | True |
| | 70–79 | 70–79 | 4.641737 | 0.000200 | 27 | 18 | 0.000750 | True |
| | 80–89 | 80–89 | 3.890121 | 0.000400 | 25 | 29 | 0.001091 | True |
| gldm_DependenceEntropy | 60–69 | 60–69 | -0.417177 | 0.694331 | 10 | 7 | 0.694331 | False |
| | 70–79 | 70–79 | -1.295002 | 0.210179 | 27 | 18 | 0.225192 | False |
| | 80–89 | 80–89 | -4.297483 | 0.000200 | 25 | 29 | 0.000750 | True |
| shape_VoxelVolume | 60–69 | 60–69 | 3.391793 | 0.008199 | 10 | 7 | 0.012599 | True |
| | 70–79 | 70–79 | 3.493108 | 0.000800 | 27 | 18 | 0.001846 | True |
| | 80–89 | 80–89 | 4.519004 | 0.000200 | 25 | 29 | 0.000750 | True |
| shape_MeshVolume | 60–69 | 60–69 | 3.347486 | 0.005999 | 10 | 7 | 0.009999 | True |
| | 70–79 | 70–79 | 3.518587 | 0.000600 | 27 | 18 | 0.001500 | True |
| | 80–89 | 80–89 | 4.518875 | 0.000200 | 25 | 29 | 0.000750 | True |
| glrlm_GrayLevelNonUniformity | 60–69 | 60–69 | 3.573880 | 0.002000 | 10 | 7 | 0.004285 | True |
| | 70–79 | 70–79 | 5.058072 | 0.000200 | 27 | 18 | 0.000750 | True |
| | 80–89 | 80–89 | 5.961795 | 0.000200 | 25 | 29 | 0.000750 | True |

### 4 Complete AUC and Effect Size Results

#### 4.1 Hippocampus

TABLE XXI: Complete AUC, Cohen's $d$, and Hedges' $g$ results for hippocampus radiomic features.

| Feature | Age bin | AUC [95% CI] | Direction | Cohen's $d$ [95% CI] | Hedges' $g$ [95% CI] |
|---|---|---|---|---|---|
| | 80–89 | 0.815 [0.690, 0.927] | AD < CN | -1.237 [-1.951, -0.691] | -1.220 [-1.923, -0.681] |
| hippocampus_glcm_ClusterShade | 70–79 | 0.837 [0.703, 0.947] | AD < CN | -1.357 [-2.190, -0.763] | -1.333 [-2.152, -0.750] |
| | 60–69 | 0.786 [0.533, 1.000] | AD < CN | -1.146 [-3.299, -0.154] | -1.087 [-3.131, -0.147] |
| | 80–89 | 0.902 [0.812, 0.971] | AD < CN | -1.846 [-2.623, -1.284] | -1.819 [-2.585, -1.265] |
| hippocampus_gldm_GrayLevelNonUniformity | 70–79 | 0.912 [0.816, 0.982] | AD < CN | -1.756 [-2.531, -1.220] | -1.725 [-2.487, -1.198] |
| | 60–69 | 0.814 [0.569, 0.986] | AD < CN | -1.314 [-2.511, -0.527] | -1.247 [-2.383, -0.500] |
| | 80–89 | 0.898 [0.791, 0.979] | AD < CN | -1.728 [-2.643, -1.106] | -1.703 [-2.605, -1.090] |
| hippocampus_firstorder_Uniformity | 70–79 | 0.825 [0.692, 0.934] | AD < CN | -1.282 [-1.947, -0.797] | -1.259 [-1.913, -0.783] |
| | 60–69 | 0.571 [0.500, 0.957] | AD < CN | -0.175 [-1.721, 0.934] | -0.166 [-1.634, 0.886] |
| | 80–89 | 0.880 [0.781, 0.957] | AD > CN | 1.572 [1.067, 2.253] | 1.549 [1.051, 2.220] |
| hippocampus_glcm_ClusterTendency | 70–79 | 0.823 [0.686, 0.938] | AD > CN | 1.274 [0.733, 2.009] | 1.252 [0.721, 1.974] |
| | 60–69 | 0.700 [0.514, 0.981] | AD > CN | 0.754 [-0.215, 2.541] | 0.716 [-0.204, 2.411] |
| | 80–89 | 0.829 [0.707, 0.930] | AD > CN | 1.308 [0.794, 1.977] | 1.290 [0.782, 1.948] |
| hippocampus_glcm_MCC | 70–79 | 0.728 [0.563, 0.870] | AD > CN | 0.832 [0.260, 1.509] | 0.818 [0.255, 1.483] |
| | 60–69 | 0.971 [0.867, 1.000] | AD > CN | 2.131 [1.466, 3.816] | 2.023 [1.392, 3.622] |
| | 80–89 | 0.855 [0.746, 0.944] | AD > CN | 1.482 [0.985, 2.139] | 1.460 [0.970, 2.108] |
| hippocampus_glcm_SumSquares | 70–79 | 0.819 [0.682, 0.933] | AD > CN | 1.305 [0.742, 2.077] | 1.282 [0.729, 2.041] |
| | 60–69 | 0.600 [0.500, 0.923] | AD > CN | 0.406 [-0.577, 1.946] | 0.385 [-0.547, 1.847] |
| | 80–89 | 0.843 [0.730, 0.935] | AD > CN | 1.446 [0.941, 2.085] | 1.426 [0.928, 2.054] |
| hippocampus_gldm_GrayLevelVariance | 70–79 | 0.813 [0.675, 0.927] | AD > CN | 1.253 [0.695, 2.016] | 1.231 [0.683, 1.980] |
| | 60–69 | 0.600 [0.500, 0.923] | AD > CN | 0.449 [-0.510, 1.973] | 0.427 [-0.484, 1.873] |
| | 80–89 | 0.822 [0.703, 0.924] | AD > CN | 1.309 [0.741, 2.041] | 1.290 [0.731, 2.011] |
| hippocampus_firstorder_Entropy | 70–79 | 0.800 [0.660, 0.921] | AD > CN | 1.173 [0.616, 1.936] | 1.152 [0.606, 1.902] |
| | 60–69 | 0.571 [0.500, 0.917] | AD > CN | 0.204 [-0.774, 1.647] | 0.193 [-0.735, 1.563] |
| | 80–89 | 0.840 [0.728, 0.933] | AD > CN | 1.380 [0.894, 2.004] | 1.360 [0.881, 1.975] |
| hippocampus_firstorder_Variance | 70–79 | 0.815 [0.675, 0.929] | AD > CN | 1.213 [0.671, 1.898] | 1.191 [0.659, 1.865] |

Continued on next page

TABLE XXI – continued from previous page

| Feature | Age bin | AUC [95% CI] | Direction | Cohen's $d$ [95% CI] | Hedges' $g$ [95% CI] |
|---|---|---|---|---|---|
| | 60–69 | 0.643 [0.500, 0.929] | AD > CN | 0.506 [-0.401, 2.104] | 0.481 [-0.380, 1.997] |
| | 80–89 | 0.862 [0.750, 0.951] | AD > CN | 1.484 [0.899, 2.270] | 1.463 [0.886, 2.237] |
| hippocampus_glcm_SumEntropy | 70–79 | 0.805 [0.665, 0.924] | AD > CN | 1.242 [0.692, 2.002] | 1.221 [0.679, 1.966] |
| | 60–69 | 0.543 [0.500, 0.923] | AD > CN | 0.248 [-0.796, 1.822] | 0.235 [-0.755, 1.729] |

#### *4.2 Entorhinal Cortex*

TABLE XXII: Complete AUC, Cohen's $d$, and Hedges' $g$ results for entorhinal cortex radiomic features.

| Feature | Age bin | AUC [95% CI] | Direction | Cohen's $d$ [95% CI] | Hedges' $g$ [95% CI] |
|---|---|---|---|---|---|
| | 80–89 | 0.766 [0.632, 0.883] | AD < CN | -1.048 [-1.650, -0.559] | -1.033 [-1.626, -0.551] |
| entorhinal_cortex_glcm_ClusterShade | 70–79 | 0.835 [0.692, 0.953] | AD < CN | -1.421 [-2.177, -0.858] | -1.396 [-2.139, -0.843] |
| | 60–69 | 0.886 [0.681, 1.000] | AD < CN | -1.419 [-2.711, -0.634] | -1.346 [-2.573, -0.602] |
| | 80–89 | 0.623 [0.509, 0.774] | AD > CN | 0.404 [-0.136, 0.995] | 0.398 [-0.134, 0.981] |
| entorhinal_cortex_firstorder_Skewness | 70–79 | 0.591 [0.505, 0.752] | AD < CN | -0.369 [-0.959, 0.196] | -0.362 [-0.942, 0.192] |
| | 60–69 | 0.800 [0.556, 1.000] | AD < CN | -1.127 [-2.280, -0.400] | -1.069 [-2.164, -0.380] |
| | 80–89 | 0.801 [0.669, 0.922] | AD > CN | 0.984 [0.491, 1.664] | 0.969 [0.484, 1.640] |
| entorhinal_cortex_shape_SurfaceVolumeRatio | 70–79 | 0.854 [0.702, 0.973] | AD > CN | 1.322 [0.780, 2.025] | 1.299 [0.766, 1.989] |
| | 60–69 | 0.800 [0.550, 0.986] | AD > CN | 1.106 [0.257, 2.275] | 1.050 [0.244, 2.160] |
| | 80–89 | 0.670 [0.532, 0.809] | AD > CN | 0.642 [0.160, 1.144] | 0.632 [0.158, 1.128] |
| entorhinal_cortex_firstorder_Median | 70–79 | 0.737 [0.560, 0.887] | AD > CN | 0.855 [0.274, 1.516] | 0.840 [0.269, 1.490] |
| | 60–69 | 0.729 [0.517, 0.944] | AD > CN | 0.806 [-0.106, 2.154] | 0.765 [-0.100, 2.045] |
| | 80–89 | 0.752 [0.609, 0.872] | AD > CN | 0.858 [0.347, 1.485] | 0.846 [0.342, 1.463] |
| entorhinal_cortex_glrlm_ShortRunLowGrayLevelEmphasis | 70–79 | 0.765 [0.611, 0.904] | AD > CN | 1.007 [0.449, 1.640] | 0.990 [0.441, 1.612] |
| | 60–69 | 0.857 [0.625, 1.000] | AD > CN | 1.272 [0.303, 3.053] | 1.208 [0.288, 2.898] |
| | 80–89 | 0.770 [0.635, 0.889] | AD < CN | -1.064 [-1.772, -0.540] | -1.049 [-1.746, -0.532] |
| entorhinal_cortex_gldm_DependenceNonUniformity | 70–79 | 0.819 [0.684, 0.926] | AD < CN | -1.340 [-1.995, -0.838] | -1.317 [-1.960, -0.824] |
| | 60–69 | 0.914 [0.722, 1.000] | AD < CN | -1.718 [-2.878, -1.036] | -1.631 [-2.732, -0.984] |
| | 80–89 | 0.800 [0.666, 0.918] | AD > CN | 1.151 [0.588, 1.935] | 1.135 [0.580, 1.907] |
| entorhinal_cortex_gldm_DependenceEntropy | 70–79 | 0.609 [0.506, 0.784] | AD > CN | 0.419 [-0.203, 1.119] | 0.411 [-0.200, 1.099] |
| | 60–69 | 0.557 [0.500, 0.848] | AD > CN | 0.199 [-0.774, 1.243] | 0.189 [-0.734, 1.179] |
| | 80–89 | 0.818 [0.697, 0.923] | AD < CN | -1.241 [-1.911, -0.764] | -1.223 [-1.883, -0.753] |
| entorhinal_cortex_shape_VoxelVolume | 70–79 | 0.767 [0.606, 0.904] | AD < CN | -1.140 [-1.795, -0.597] | -1.120 [-1.764, -0.586] |
| | 60–69 | 0.871 [0.636, 1.000] | AD < CN | -1.588 [-2.824, -0.826] | -1.507 [-2.681, -0.784] |
| | 80–89 | 0.817 [0.694, 0.923] | AD < CN | -1.241 [-1.910, -0.766] | -1.223 [-1.882, -0.755] |
| entorhinal_cortex_shape_MeshVolume | 70–79 | 0.770 [0.609, 0.905] | AD < CN | -1.148 [-1.800, -0.607] | -1.128 [-1.769, -0.596] |
| | 60–69 | 0.871 [0.636, 1.000] | AD < CN | -1.569 [-2.812, -0.804] | -1.489 [-2.669, -0.763] |
| | 80–89 | 0.876 [0.777, 0.960] | AD < CN | -1.629 [-2.421, -1.085] | -1.605 [-2.386, -1.069] |
| entorhinal_cortex_glrlm_GrayLevelNonUniformity | 70–79 | 0.868 [0.750, 0.957] | AD < CN | -1.678 [-2.364, -1.178] | -1.649 [-2.322, -1.157] |
| | 60–69 | 0.914 [0.700, 1.000] | AD < CN | -1.872 [-3.532, -0.972] | -1.777 [-3.352, -0.923] |

#### *4.3 Cingulum*

TABLE XXIII: Complete AUC, Cohen's $d$, and Hedges' $g$ results for cingulum radiomic features.

| Feature | Age bin | AUC [95% CI] | Direction | Cohen's $d$ [95% CI] | Hedges' $g$ [95% CI] |
|---|---|---|---|---|---|
| | 80–89 | 0.590 [0.505, 0.743] | AD > CN | 0.247 [-0.267, 0.879] | 0.243 [-0.263, 0.867] |
| cingulum_firstorder_Median | 70–79 | 0.636 [0.510, 0.796] | AD > CN | 0.573 [-0.017, 1.256] | 0.563 [-0.017, 1.234] |
| | 60–69 | 0.671 [0.514, 0.939] | AD > CN | 0.772 [-0.270, 1.989] | 0.733 [-0.256, 1.888] |
| | 80–89 | 0.868 [0.756, 0.958] | AD < CN | -1.589 [-2.505, -0.967] | -1.566 [-2.468, -0.953] |
| cingulum_firstorder_Kurtosis | 70–79 | 0.741 [0.561, 0.889] | AD < CN | -1.018 [-1.827, -0.380] | -1.000 [-1.795, -0.373] |
| | 60–69 | 0.871 [0.628, 1.000] | AD < CN | -1.573 [-3.425, -0.576] | -1.493 [-3.251, -0.547] |
| | 80–89 | 0.783 [0.645, 0.901] | AD > CN | 1.073 [0.541, 1.757] | 1.058 [0.533, 1.732] |
| cingulum_firstorder_InterquartileRange | 70–79 | 0.737 [0.567, 0.881] | AD > CN | 0.990 [0.373, 1.721] | 0.973 [0.367, 1.691] |
| | 60–69 | 0.771 [0.530, 1.000] | AD > CN | 1.038 [0.066, 2.629] | 0.985 [0.063, 2.495] |
| | 80–89 | 0.753 [0.612, 0.878] | AD > CN | 0.933 [0.418, 1.569] | 0.920 [0.412, 1.547] |
| cingulum_firstorder_RobustMeanAbsoluteDeviation | 70–79 | 0.685 [0.524, 0.836] | AD > CN | 0.879 [0.251, 1.604] | 0.863 [0.246, 1.575] |
| | 60–69 | 0.700 [0.514, 0.957] | AD > CN | 0.910 [-0.099, 2.429] | 0.864 [-0.094, 2.306] |
| | 80–89 | 0.534 [0.502, 0.700] | AD > CN | 0.123 [-0.393, 0.738] | 0.121 [-0.387, 0.727] |
| cingulum_firstorder_Mean | 70–79 | 0.607 [0.506, 0.770] | AD > CN | 0.414 [-0.188, 1.069] | 0.407 [-0.185, 1.050] |
| | 60–69 | 0.600 [0.500, 0.955] | AD > CN | 0.458 [-0.650, 1.923] | 0.435 [-0.617, 1.825] |
| | 80–89 | 0.778 [0.646, 0.896] | AD > CN | 0.892 [0.434, 1.440] | 0.879 [0.428, 1.419] |
| cingulum_glszm_GrayLevelNonUniformityNormalized | 70–79 | 0.698 [0.532, 0.860] | AD > CN | 0.751 [0.168, 1.351] | 0.738 [0.165, 1.328] |
| | 60–69 | 0.829 [0.571, 1.000] | AD > CN | 1.340 [0.610, 2.514] | 1.272 [0.579, 2.387] |
| | 80–89 | 0.732 [0.585, 0.863] | AD > CN | 0.895 [0.372, 1.582] | 0.882 [0.367, 1.559] |
| cingulum_firstorder_Minimum | 70–79 | 0.718 [0.553, 0.878] | AD > CN | 0.697 [0.082, 1.529] | 0.684 [0.080, 1.503] |

Continued on next page

TABLE XXIII – continued from previous page

| Feature | Age bin | AUC [95% CI] | Direction | Cohen's $d$ [95% CI] | Hedges' $g$ [95% CI] |
|---|---|---|---|---|---|
| | 60–69 | 0.929 [0.773, 1.000] | AD > CN | 1.534 [1.027, 2.594] | 1.456 [0.974, 2.462] |
| | 80–89 | 0.539 [0.503, 0.703] | AD > CN | 0.116 [-0.401, 0.745] | 0.114 [-0.396, 0.734] |
| cingulum_firstorder_90Percentile | 70–79 | 0.572 [0.504, 0.738] | AD > CN | 0.303 [-0.305, 0.950] | 0.297 [-0.299, 0.933] |
| | 60–69 | 0.614 [0.500, 0.944] | AD > CN | 0.159 [-1.040, 1.863] | 0.151 [-0.987, 1.768] |
| | 80–89 | 0.659 [0.521, 0.801] | AD > CN | 0.463 [-0.046, 1.063] | 0.457 [-0.046, 1.048] |
| cingulum_firstorder_MeanAbsoluteDeviation | 70–79 | 0.611 [0.506, 0.782] | AD > CN | 0.526 [-0.115, 1.223] | 0.516 [-0.113, 1.202] |
| | 60–69 | 0.586 [0.500, 0.885] | AD > CN | 0.408 [-0.695, 1.718] | 0.387 [-0.659, 1.630] |
| | 80–89 | 0.541 [0.503, 0.699] | AD < CN | -0.177 [-0.721, 0.356] | -0.175 [-0.711, 0.351] |
| cingulum_firstorder_Energy | 70–79 | 0.574 [0.504, 0.752] | AD < CN | -0.068 [-0.829, 0.533] | -0.067 [-0.815, 0.524] |
| | 60–69 | 0.743 [0.517, 0.971] | AD < CN | -0.931 [-2.066, -0.151] | -0.883 [-1.961, -0.143] |

#### *4.4 Caudate Nucleus*

TABLE XXIV: Complete AUC, Cohen's $d$, and Hedges' $g$ results for caudate nucleus radiomic features.

| Feature | Age bin | AUC [95% CI] | Direction | Cohen's $d$ [95% CI] | Hedges' $g$ [95% CI] |
|---|---|---|---|---|---|
| | 80–89 | 0.720 [0.571, 0.844] | AD > CN | 0.775 [0.271, 1.328] | 0.764 [0.267, 1.309] |
| caudate_nucleus_firstorder_Uniformity | 70–79 | 0.619 [0.508, 0.778] | AD < CN | -0.448 [-1.023, 0.094] | -0.440 [-1.005, 0.092] |
| | 60–69 | 0.686 [0.514, 0.962] | AD < CN | -0.359 [-2.308, 0.562] | -0.341 [-2.191, 0.534] |
| | 80–89 | 0.680 [0.533, 0.818] | AD < CN | -0.729 [-1.270, -0.233] | -0.719 [-1.251, -0.230] |
| caudate_nucleus_glcm_SumSquares | 70–79 | 0.599 [0.504, 0.762] | AD > CN | 0.397 [-0.164, 0.999] | 0.390 [-0.161, 0.981] |
| | 60–69 | 0.600 [0.500, 0.909] | AD > CN | 0.237 [-0.664, 1.804] | 0.225 [-0.630, 1.712] |
| | 80–89 | 0.724 [0.576, 0.851] | AD > CN | 0.888 [0.388, 1.479] | 0.875 [0.383, 1.457] |
| caudate_nucleus_glrlm_RunVariance | 70–79 | 0.640 [0.510, 0.798] | AD < CN | -0.557 [-1.083, -0.052] | -0.547 [-1.064, -0.051] |
| | 60–69 | 0.671 [0.514, 0.944] | AD < CN | -0.153 [-1.873, 0.746] | -0.145 [-1.778, 0.708] |
| | 80–89 | 0.746 [0.602, 0.865] | AD > CN | 0.890 [0.396, 1.463] | 0.877 [0.390, 1.442] |
| caudate_nucleus_glcm_JointEnergy | 70–79 | 0.613 [0.506, 0.774] | AD < CN | -0.463 [-1.027, 0.079] | -0.455 [-1.009, 0.077] |
| | 60–69 | 0.671 [0.514, 0.950] | AD < CN | -0.369 [-2.236, 0.543] | -0.350 [-2.122, 0.516] |
| | 80–89 | 0.698 [0.549, 0.831] | AD > CN | 0.765 [0.261, 1.341] | 0.753 [0.257, 1.321] |
| caudate_nucleus_gldm_LargeDependenceEmphasis | 70–79 | 0.660 [0.516, 0.815] | AD < CN | -0.575 [-1.185, -0.039] | -0.565 [-1.165, -0.039] |
| | 60–69 | 0.671 [0.514, 0.944] | AD < CN | -0.264 [-2.059, 0.617] | -0.251 [-1.954, 0.586] |
| | 80–89 | 0.677 [0.530, 0.817] | AD < CN | -0.710 [-1.288, -0.191] | -0.700 [-1.269, -0.188] |
| caudate_nucleus_glrlm_RunPercentage | 70–79 | 0.663 [0.517, 0.819] | AD > CN | 0.571 [0.036, 1.177] | 0.561 [0.035, 1.156] |
| | 60–69 | 0.686 [0.514, 0.967] | AD > CN | 0.217 [-0.655, 2.014] | 0.206 [-0.621, 1.912] |
| | 80–89 | 0.673 [0.530, 0.811] | AD < CN | -0.676 [-1.200, -0.176] | -0.666 [-1.183, -0.174] |
| caudate_nucleus_glcm_ClusterTendency | 70–79 | 0.599 [0.506, 0.761] | AD > CN | 0.353 [-0.217, 0.964] | 0.347 [-0.213, 0.948] |
| | 60–69 | 0.614 [0.500, 0.917] | AD > CN | 0.305 [-0.612, 1.896] | 0.289 [-0.580, 1.800] |
| | 80–89 | 0.568 [0.504, 0.731] | AD > CN | 0.031 [-0.489, 0.627] | 0.031 [-0.482, 0.618] |
| caudate_nucleus_firstorder_Median | 70–79 | 0.576 [0.504, 0.742] | AD > CN | 0.232 [-0.394, 0.880] | 0.227 [-0.387, 0.864] |
| | 60–69 | 0.829 [0.550, 1.000] | AD > CN | 1.335 [0.433, 2.882] | 1.267 [0.411, 2.735] |
| | 80–89 | 0.712 [0.564, 0.840] | AD > CN | 0.821 [0.328, 1.409] | 0.810 [0.324, 1.388] |
| caudate_nucleus_glrlm_LongRunEmphasis | 70–79 | 0.644 [0.509, 0.802] | AD < CN | -0.569 [-1.066, -0.073] | -0.559 [-1.047, -0.072] |
| | 60–69 | 0.671 [0.514, 0.957] | AD < CN | -0.020 [-1.798, 0.858] | -0.019 [-1.707, 0.814] |
| | 80–89 | 0.553 [0.503, 0.714] | AD > CN | 0.001 [-0.517, 0.596] | 0.001 [-0.509, 0.587] |
| caudate_nucleus_firstorder_90Percentile | 70–79 | 0.564 [0.504, 0.735] | AD > CN | 0.211 [-0.405, 0.862] | 0.208 [-0.397, 0.847] |
| | 60–69 | 0.757 [0.514, 1.000] | AD > CN | 0.860 [-0.189, 2.398] | 0.816 [-0.179, 2.276] |

#### *4.5 Thalamus*

TABLE XXV: Complete AUC, Cohen's $d$, and Hedges' $g$ results for thalamus radiomic features.

| Feature | Age bin | AUC [95% CI] | Direction | Cohen's $d$ [95% CI] | Hedges' $g$ [95% CI] |
|---|---|---|---|---|---|
| | 80–89 | 0.508 [0.503, 0.682] | AD > CN | -0.002 [-0.546, 0.553] | -0.002 [-0.538, 0.545] |
| thalamus_glrlm_GrayLevelNonUniformityNormalized | 70–79 | 0.665 [0.514, 0.826] | AD < CN | -0.538 [-1.311, 0.095] | -0.529 [-1.288, 0.093] |
| | 60–69 | 0.729 [0.515, 0.967] | AD < CN | -0.803 [-2.210, 0.123] | -0.763 [-2.098, 0.117] |
| | 80–89 | 0.523 [0.503, 0.687] | AD < CN | -0.089 [-0.622, 0.475] | -0.088 [-0.613, 0.468] |
| thalamus_glrlm_GrayLevelVariance | 70–79 | 0.572 [0.504, 0.737] | AD > CN | 0.245 [-0.367, 0.941] | 0.240 [-0.361, 0.924] |
| | 60–69 | 0.743 [0.519, 0.981] | AD > CN | 0.976 [-0.009, 2.715] | 0.927 [-0.009, 2.577] |
| | 80–89 | 0.543 [0.503, 0.705] | AD > CN | 0.113 [-0.408, 0.721] | 0.111 [-0.402, 0.711] |
| thalamus_glcm_ClusterProminence | 70–79 | 0.586 [0.504, 0.761] | AD > CN | 0.380 [-0.288, 0.980] | 0.373 [-0.283, 0.963] |
| | 60–69 | 0.743 [0.515, 1.000] | AD > CN | 1.039 [0.238, 2.194] | 0.987 [0.226, 2.082] |
| | 80–89 | 0.559 [0.504, 0.717] | AD > CN | 0.179 [-0.357, 0.787] | 0.176 [-0.352, 0.776] |
| thalamus_firstorder_Variance | 70–79 | 0.586 [0.505, 0.758] | AD > CN | 0.360 [-0.271, 1.015] | 0.354 [-0.266, 0.997] |
| | 60–69 | 0.671 [0.500, 0.971] | AD > CN | 0.724 [-0.339, 1.956] | 0.687 [-0.322, 1.857] |
| | 80–89 | 0.557 [0.503, 0.718] | AD > CN | 0.188 [-0.343, 0.805] | 0.185 [-0.338, 0.793] |
| thalamus_firstorder_RootMeanSquared | 70–79 | 0.506 [0.502, 0.698] | AD < CN | -0.082 [-0.698, 0.543] | -0.080 [-0.685, 0.534] |
| | 60–69 | 0.714 [0.514, 1.000] | AD > CN | 1.115 [0.213, 2.498] | 1.058 [0.202, 2.371] |

Continued on next page

TABLE XXV – continued from previous page

| Feature | Age bin | AUC [95% CI] | Direction | Cohen's $d$ [95% CI] | Hedges' $g$ [95% CI] |
|---|---|---|---|---|---|
| | 80–89 | 0.545 [0.503, 0.710] | AD > CN | 0.115 [-0.413, 0.717] | 0.113 [-0.407, 0.707] |
| thalamus_firstorder_90Percentile | 70–79 | 0.516 [0.504, 0.704] | AD < CN | -0.069 [-0.697, 0.562] | -0.068 [-0.685, 0.553] |
| | 60–69 | 0.671 [0.500, 1.000] | AD > CN | 0.779 [-0.413, 2.437] | 0.739 [-0.392, 2.313] |
| | 80–89 | 0.572 [0.504, 0.733] | AD > CN | 0.172 [-0.356, 0.784] | 0.170 [-0.351, 0.773] |
| thalamus_firstorder_Median | 70–79 | 0.525 [0.504, 0.702] | AD < CN | -0.087 [-0.701, 0.530] | -0.085 [-0.689, 0.521] |
| | 60–69 | 0.657 [0.500, 1.000] | AD > CN | 0.970 [-0.121, 2.734] | 0.921 [-0.115, 2.595] |
| | 80–89 | 0.560 [0.504, 0.723] | AD > CN | 0.125 [-0.396, 0.725] | 0.123 [-0.391, 0.714] |
| thalamus_firstorder_Mean | 70–79 | 0.531 [0.503, 0.707] | AD < CN | -0.112 [-0.727, 0.501] | -0.110 [-0.715, 0.492] |
| | 60–69 | 0.671 [0.500, 1.000] | AD > CN | 0.896 [-0.231, 2.619] | 0.851 [-0.219, 2.486] |
| | 80–89 | 0.545 [0.503, 0.709] | AD > CN | 0.001 [-0.515, 0.650] | 0.001 [-0.508, 0.641] |
| thalamus_firstorder_TotalEnergy | 70–79 | 0.500 [0.502, 0.699] | AD > CN | -0.076 [-0.686, 0.544] | -0.075 [-0.674, 0.535] |
| | 60–69 | 0.657 [0.514, 0.967] | AD > CN | 0.954 [0.046, 2.118] | 0.905 [0.044, 2.011] |
| | 80–89 | 0.542 [0.503, 0.707] | AD > CN | -0.008 [-0.520, 0.640] | -0.008 [-0.513, 0.631] |
| thalamus_firstorder_Energy | 70–79 | 0.500 [0.502, 0.699] | AD > CN | -0.080 [-0.694, 0.543] | -0.078 [-0.681, 0.533] |
| | 60–69 | 0.657 [0.514, 0.967] | AD > CN | 0.954 [0.046, 2.118] | 0.905 [0.044, 2.011] |

#### *4.6 Rolandic Operculum*

TABLE XXVI: Complete AUC, Cohen's $d$, and Hedges' $g$ results for rolandic operculum radiomic features.

| Feature | Age bin | AUC [95% CI] | Direction | Cohen's $d$ [95% CI] | Hedges' $g$ [95% CI] |
|---|---|---|---|---|---|
| | 80–89 | 0.661 [0.520, 0.801] | AD < CN | -0.515 [-1.164, 0.006] | -0.507 [-1.147, 0.006] |
| rolandic_operculum_firstorder_Kurtosis | 70–79 | 0.681 [0.519, 0.841] | AD < CN | -0.592 [-1.403, 0.051] | -0.581 [-1.378, 0.050] |
| | 60–69 | 0.686 [0.514, 0.943] | AD < CN | -0.928 [-2.249, 0.052] | -0.881 [-2.135, 0.050] |
| | 80–89 | 0.636 [0.510, 0.784] | AD > CN | 0.462 [-0.065, 1.099] | 0.456 [-0.064, 1.083] |
| rolandic_operculum_ngtdm_Contrast | 70–79 | 0.525 [0.502, 0.710] | AD > CN | 0.115 [-0.503, 0.745] | 0.113 [-0.494, 0.732] |
| | 60–69 | 0.543 [0.500, 0.900] | AD < CN | -0.053 [-1.390, 0.960] | -0.051 [-1.319, 0.912] |
| | 80–89 | 0.768 [0.626, 0.884] | AD > CN | 1.041 [0.538, 1.672] | 1.026 [0.530, 1.648] |
| rolandic_operculum_firstorder_Minimum | 70–79 | 0.722 [0.556, 0.875] | AD > CN | 0.808 [0.215, 1.614] | 0.794 [0.211, 1.586] |
| | 60–69 | 0.843 [0.621, 1.000] | AD > CN | 1.305 [0.625, 2.322] | 1.239 [0.594, 2.204] |
| | 80–89 | 0.636 [0.512, 0.789] | AD < CN | -0.446 [-1.090, 0.066] | -0.440 [-1.074, 0.065] |
| rolandic_operculum_glcm_Idmn | 70–79 | 0.553 [0.504, 0.728] | AD < CN | -0.206 [-0.839, 0.404] | -0.202 [-0.825, 0.397] |
| | 60–69 | 0.514 [0.500, 0.867] | AD > CN | -0.075 [-1.074, 1.460] | -0.071 [-1.020, 1.385] |
| | 80–89 | 0.714 [0.558, 0.854] | AD < CN | -0.737 [-1.472, -0.187] | -0.726 [-1.451, -0.184] |
| rolandic_operculum_firstorder_Range | 70–79 | 0.607 [0.506, 0.788] | AD < CN | -0.460 [-1.164, 0.167] | -0.452 [-1.144, 0.164] |
| | 60–69 | 0.829 [0.596, 1.000] | AD < CN | -1.380 [-2.581, -0.624] | -1.310 [-2.449, -0.593] |
| | 80–89 | 0.641 [0.512, 0.784] | AD < CN | -0.535 [-1.132, -0.007] | -0.527 [-1.116, -0.007] |
| rolandic_operculum_glcm_MaximumProbability | 70–79 | 0.656 [0.516, 0.817] | AD < CN | -0.591 [-1.256, -0.003] | -0.580 [-1.234, -0.003] |
| | 60–69 | 0.571 [0.500, 0.914] | AD < CN | 0.154 [-1.693, 1.306] | 0.146 [-1.606, 1.240] |
| | 80–89 | 0.626 [0.510, 0.779] | AD < CN | -0.334 [-0.955, 0.171] | -0.329 [-0.942, 0.168] |
| rolandic_operculum_glcm_Idn | 70–79 | 0.580 [0.504, 0.748] | AD < CN | -0.255 [-0.885, 0.347] | -0.251 [-0.870, 0.341] |
| | 60–69 | 0.557 [0.500, 0.857] | AD > CN | 0.396 [-0.576, 1.563] | 0.376 [-0.546, 1.483] |
| | 80–89 | 0.581 [0.504, 0.746] | AD > CN | 0.210 [-0.323, 0.820] | 0.207 [-0.319, 0.808] |
| rolandic_operculum_glszm_GrayLevelNonUniformityNormalized | 70–79 | 0.617 [0.506, 0.793] | AD > CN | 0.315 [-0.278, 1.044] | 0.309 [-0.273, 1.026] |
| | 60–69 | 0.800 [0.543, 1.000] | AD > CN | 1.485 [0.637, 3.195] | 1.409 [0.605, 3.032] |
| | 80–89 | 0.629 [0.510, 0.781] | AD < CN | -0.445 [-0.947, 0.051] | -0.439 [-0.934, 0.051] |
| rolandic_operculum_glszm_ZoneEntropy | 70–79 | 0.718 [0.552, 0.868] | AD < CN | -0.905 [-1.657, -0.302] | -0.889 [-1.628, -0.297] |
| | 60–69 | 0.729 [0.500, 1.000] | AD < CN | -0.926 [-2.680, 0.169] | -0.879 [-2.543, 0.160] |
| | 80–89 | 0.641 [0.511, 0.786] | AD < CN | -0.549 [-1.197, -0.020] | -0.541 [-1.180, -0.020] |
| rolandic_operculum_glrlm_LongRunHighGrayLevelEmphasis | 70–79 | 0.695 [0.537, 0.850] | AD < CN | -0.729 [-1.508, -0.116] | -0.716 [-1.481, -0.114] |
| | 60–69 | 0.543 [0.500, 0.885] | AD > CN | -0.042 [-0.948, 1.136] | -0.040 [-0.900, 1.079] |

#### *4.7 Vermis*

TABLE XXVII: Complete AUC, Cohen's $d$, and Hedges' $g$ results for vermis radiomic features.

| Feature | Age bin | AUC [95% CI] | Direction | Cohen's $d$ [95% CI] | Hedges' $g$ [95% CI] |
|---|---|---|---|---|---|
| | 80–89 | 0.561 [0.503, 0.714] | AD > CN | 0.277 [-0.261, 0.852] | 0.273 [-0.257, 0.840] |
| vermis_shape_Elongation | 70–79 | 0.673 [0.525, 0.826] | AD < CN | -0.588 [-1.160, -0.078] | -0.578 [-1.139, -0.077] |
| | 60–69 | 0.586 [0.500, 0.867] | AD < CN | -0.196 [-1.325, 0.830] | -0.186 [-1.257, 0.788] |
| | 80–89 | 0.668 [0.525, 0.807] | AD < CN | -0.638 [-1.256, -0.105] | -0.628 [-1.238, -0.103] |
| vermis_glcm_Imc1 | 70–79 | 0.722 [0.560, 0.871] | AD < CN | -0.926 [-1.574, -0.326] | -0.909 [-1.547, -0.320] |
| | 60–69 | 0.886 [0.686, 1.000] | AD < CN | -1.464 [-2.924, -0.671] | -1.389 [-2.775, -0.637] |
| | 80–89 | 0.632 [0.509, 0.778] | AD > CN | 0.472 [-0.058, 1.068] | 0.465 [-0.057, 1.053] |
| vermis_glcm_Imc2 | 70–79 | 0.702 [0.541, 0.858] | AD > CN | 0.763 [0.136, 1.432] | 0.750 [0.133, 1.407] |
| | 60–69 | 0.786 [0.543, 0.981] | AD > CN | 1.117 [0.262, 2.542] | 1.060 [0.248, 2.413] |

Continued on next page

TABLE XXVII – continued from previous page

| Feature | Age bin | AUC [95% CI] | Direction | Cohen's $d$ [95% CI] | Hedges' $g$ [95% CI] |
|---|---|---|---|---|---|
| | 80–89 | 0.614 [0.506, 0.764] | AD > CN | 0.401 [-0.136, 0.985] | 0.395 [-0.134, 0.970] |
| vermis_glcm_MCC | 70–79 | 0.702 [0.536, 0.859] | AD > CN | 0.805 [0.164, 1.501] | 0.791 [0.161, 1.474] |
| | 60–69 | 0.743 [0.528, 0.950] | AD > CN | 1.032 [0.171, 2.280] | 0.979 [0.162, 2.164] |
| | 80–89 | 0.514 [0.503, 0.677] | AD < CN | -0.035 [-0.587, 0.486] | -0.034 [-0.578, 0.479] |
| vermis_firstorder_Kurtosis | 70–79 | 0.506 [0.502, 0.720] | AD > CN | 0.054 [-0.606, 0.700] | 0.053 [-0.595, 0.688] |
| | 60–69 | 0.586 [0.500, 0.871] | AD < CN | -0.381 [-1.561, 0.581] | -0.362 [-1.482, 0.552] |
| | 80–89 | 0.636 [0.511, 0.787] | AD > CN | 0.543 [0.017, 1.152] | 0.535 [0.017, 1.135] |
| vermis_glcm_Correlation | 70–79 | 0.712 [0.546, 0.867] | AD > CN | 0.827 [0.204, 1.530] | 0.812 [0.200, 1.503] |
| | 60–69 | 0.786 [0.545, 0.972] | AD > CN | 1.121 [0.275, 2.472] | 1.064 [0.261, 2.346] |
| | 80–89 | 0.521 [0.503, 0.686] | AD < CN | -0.132 [-0.683, 0.413] | -0.130 [-0.673, 0.407] |
| vermis_shape_MinorAxisLength | 70–79 | 0.586 [0.505, 0.753] | AD < CN | -0.314 [-0.912, 0.255] | -0.309 [-0.896, 0.250] |
| | 60–69 | 0.514 [0.500, 0.865] | AD < CN | -0.070 [-1.170, 0.999] | -0.066 [-1.110, 0.948] |
| | 80–89 | 0.659 [0.519, 0.801] | AD > CN | 0.654 [0.105, 1.323] | 0.645 [0.104, 1.304] |
| vermis_glrlm_LongRunLowGrayLevelEmphasis | 70–79 | 0.755 [0.601, 0.887] | AD > CN | 0.976 [0.391, 1.703] | 0.959 [0.384, 1.673] |
| | 60–69 | 0.800 [0.543, 1.000] | AD > CN | 1.308 [0.355, 3.027] | 1.241 [0.337, 2.873] |
| | 80–89 | 0.615 [0.507, 0.757] | AD > CN | 0.440 [-0.105, 0.995] | 0.433 [-0.103, 0.981] |
| vermis_glrlm_RunEntropy | 70–79 | 0.660 [0.514, 0.830] | AD > CN | 0.738 [0.115, 1.446] | 0.725 [0.113, 1.420] |
| | 60–69 | 0.743 [0.528, 0.955] | AD > CN | 0.820 [-0.043, 1.985] | 0.778 [-0.041, 1.884] |
| | 80–89 | 0.597 [0.505, 0.747] | AD > CN | 0.318 [-0.225, 0.893] | 0.314 [-0.222, 0.880] |
| vermis_gldm_DependenceVariance | 70–79 | 0.640 [0.510, 0.806] | AD > CN | 0.670 [0.103, 1.222] | 0.658 [0.101, 1.201] |
| | 60–69 | 0.871 [0.654, 1.000] | AD > CN | 1.643 [0.820, 3.091] | 1.559 [0.779, 2.933] |

#### *4.8 Putamen*

TABLE XXVIII: Complete AUC, Cohen's $d$, and Hedges' $g$ results for putamen radiomic features.

| Feature | Age bin | AUC [95% CI] | Direction | Cohen's $d$ [95% CI] | Hedges' $g$ [95% CI] |
|---|---|---|---|---|---|
| | 80–89 | 0.599 [0.505, 0.752] | AD > CN | 0.371 [-0.139, 0.946] | 0.366 [-0.137, 0.933] |
| putamen_firstorder_Median | 70–79 | 0.693 [0.525, 0.858] | AD > CN | 0.616 [0.019, 1.343] | 0.605 [0.019, 1.319] |
| | 60–69 | 0.714 [0.514, 0.981] | AD > CN | 0.735 [-0.314, 2.473] | 0.698 [-0.298, 2.348] |
| | 80–89 | 0.590 [0.505, 0.745] | AD > CN | 0.363 [-0.149, 0.938] | 0.357 [-0.147, 0.925] |
| putamen_firstorder_Mean | 70–79 | 0.691 [0.524, 0.858] | AD > CN | 0.599 [0.001, 1.336] | 0.588 [0.001, 1.312] |
| | 60–69 | 0.700 [0.514, 0.981] | AD > CN | 0.721 [-0.320, 2.417] | 0.684 [-0.303, 2.294] |
| | 80–89 | 0.586 [0.505, 0.739] | AD > CN | 0.244 [-0.271, 0.821] | 0.240 [-0.267, 0.809] |
| putamen_glcm_MaximumProbability | 70–79 | 0.634 [0.508, 0.806] | AD > CN | 0.437 [-0.164, 1.133] | 0.429 [-0.161, 1.114] |
| | 60–69 | 0.814 [0.556, 1.000] | AD > CN | 1.447 [0.455, 3.428] | 1.374 [0.431, 3.253] |
| | 80–89 | 0.554 [0.503, 0.710] | AD < CN | -0.197 [-0.766, 0.319] | -0.194 [-0.755, 0.314] |
| putamen_gldm_DependenceEntropy | 70–79 | 0.638 [0.508, 0.810] | AD < CN | -0.512 [-1.229, 0.077] | -0.503 [-1.207, 0.076] |
| | 60–69 | 0.786 [0.543, 0.972] | AD < CN | -1.362 [-2.875, -0.456] | -1.292 [-2.729, -0.433] |
| | 80–89 | 0.648 [0.518, 0.797] | AD > CN | 0.544 [0.030, 1.132] | 0.536 [0.030, 1.116] |
| putamen_firstorder_90Percentile | 70–79 | 0.691 [0.524, 0.856] | AD > CN | 0.650 [0.057, 1.375] | 0.638 [0.056, 1.351] |
| | 60–69 | 0.571 [0.500, 1.000] | AD > CN | 0.117 [-1.112, 1.758] | 0.111 [-1.055, 1.669] |
| | 80–89 | 0.578 [0.504, 0.730] | AD > CN | 0.236 [-0.284, 0.811] | 0.232 [-0.279, 0.799] |
| putamen_firstorder_Uniformity | 70–79 | 0.619 [0.507, 0.792] | AD > CN | 0.506 [-0.083, 1.214] | 0.497 [-0.082, 1.193] |
| | 60–69 | 0.814 [0.567, 1.000] | AD > CN | 1.373 [0.507, 2.940] | 1.304 [0.481, 2.791] |
| | 80–89 | 0.581 [0.504, 0.735] | AD > CN | 0.275 [-0.234, 0.842] | 0.271 [-0.231, 0.830] |
| putamen_glcm_JointEnergy | 70–79 | 0.630 [0.508, 0.800] | AD > CN | 0.528 [-0.066, 1.217] | 0.518 [-0.064, 1.195] |
| | 60–69 | 0.814 [0.556, 1.000] | AD > CN | 1.431 [0.591, 3.040] | 1.358 [0.561, 2.885] |
| | 80–89 | 0.607 [0.507, 0.758] | AD > CN | 0.376 [-0.138, 0.891] | 0.371 [-0.136, 0.878] |
| putamen_ngtdm_Coarseness | 70–79 | 0.648 [0.510, 0.810] | AD > CN | 0.774 [0.223, 1.374] | 0.760 [0.219, 1.350] |
| | 60–69 | 0.771 [0.528, 1.000] | AD > CN | 1.230 [0.402, 2.650] | 1.167 [0.382, 2.515] |
| | 80–89 | 0.590 [0.505, 0.747] | AD > CN | 0.068 [-0.407, 0.673] | 0.067 [-0.401, 0.663] |
| putamen_firstorder_10Percentile | 70–79 | 0.628 [0.508, 0.794] | AD > CN | 0.461 [-0.129, 1.185] | 0.453 [-0.127, 1.164] |
| | 60–69 | 0.786 [0.542, 0.985] | AD > CN | 1.145 [0.325, 2.376] | 1.087 [0.309, 2.255] |
| | 80–89 | 0.581 [0.504, 0.734] | AD < CN | -0.293 [-0.865, 0.244] | -0.288 [-0.852, 0.240] |
| putamen_glrlm_GrayLevelNonUniformity | 70–79 | 0.638 [0.512, 0.796] | AD < CN | -0.650 [-1.273, -0.095] | -0.639 [-1.250, -0.093] |
| | 60–69 | 0.771 [0.528, 1.000] | AD < CN | -0.988 [-2.651, 0.009] | -0.938 [-2.516, 0.009] |

#### *4.9 Insula*

TABLE XXIX: Complete AUC, Cohen's $d$, and Hedges' $g$ results for insula radiomic features.

| Feature | Age bin | AUC [95% CI] | Direction | Cohen's $d$ [95% CI] | Hedges' $g$ [95% CI] |
|---|---|---|---|---|---|
| | 80–89 | 0.662 [0.526, 0.803] | AD < CN | -0.578 [-1.137, -0.066] | -0.570 [-1.120, -0.065] |
| insula_glrlm_RunEntropy | 70–79 | 0.755 [0.591, 0.898] | AD < CN | -1.023 [-1.834, -0.411] | -1.005 [-1.802, -0.404] |
| | 60–69 | 0.743 [0.517, 1.000] | AD < CN | -0.996 [-2.470, 0.030] | -0.945 [-2.345, 0.028] |
| | 80–89 | 0.542 [0.503, 0.702] | AD > CN | 0.189 [-0.356, 0.714] | 0.186 [-0.350, 0.703] |
| insula_firstorder_RootMeanSquared | | | | | |

Continued on next page

TABLE XXIX – continued from previous page

| Feature | Age bin | AUC [95% CI] | Direction | Cohen's $d$ [95% CI] | Hedges' $g$ [95% CI] |
|---|---|---|---|---|---|
| | 70–79 | 0.603 [0.506, 0.774] | AD $<$ CN | -0.277 [-0.958, 0.332] | -0.272 [-0.941, 0.326] |
| | 60–69 | 0.586 [0.500, 0.904] | AD $<$ CN | -0.092 [-1.378, 0.973] | -0.087 [-1.308, 0.923] |
| | 80–89 | 0.701 [0.553, 0.839] | AD $>$ CN | 0.624 [0.104, 1.278] | 0.615 [0.102, 1.260] |
| insula_firstorder_Median | 70–79 | 0.667 [0.516, 0.828] | AD $>$ CN | 0.630 [0.018, 1.318] | 0.619 [0.018, 1.295] |
| | 60–69 | 0.600 [0.500, 0.962] | AD $>$ CN | 0.147 [-0.941, 1.567] | 0.140 [-0.893, 1.487] |
| | 80–89 | 0.502 [0.502, 0.679] | AD $>$ CN | 0.027 [-0.527, 0.563] | 0.027 [-0.519, 0.555] |
| insula_glrlm_GrayLevelVariance | 70–79 | 0.599 [0.505, 0.774] | AD $<$ CN | -0.417 [-1.110, 0.158] | -0.410 [-1.090, 0.155] |
| | 60–69 | 0.829 [0.586, 1.000] | AD $<$ CN | -1.568 [-3.375, -0.641] | -1.488 [-3.203, -0.609] |
| | 80–89 | 0.577 [0.505, 0.731] | AD $<$ CN | -0.282 [-0.867, 0.240] | -0.278 [-0.855, 0.236] |
| insula_firstorder_Energy | 70–79 | 0.595 [0.506, 0.765] | AD $<$ CN | -0.301 [-1.018, 0.297] | -0.296 [-1.000, 0.291] |
| | 60–69 | 0.686 [0.514, 0.950] | AD $<$ CN | -0.490 [-1.800, 0.531] | -0.465 [-1.709, 0.504] |
| | 80–89 | 0.579 [0.505, 0.733] | AD $<$ CN | -0.274 [-0.863, 0.248] | -0.270 [-0.850, 0.245] |
| insula_firstorder_TotalEnergy | 70–79 | 0.591 [0.504, 0.762] | AD $<$ CN | -0.273 [-0.988, 0.325] | -0.268 [-0.970, 0.319] |
| | 60–69 | 0.686 [0.514, 0.950] | AD $<$ CN | -0.490 [-1.800, 0.531] | -0.465 [-1.709, 0.504] |
| | 80–89 | 0.582 [0.504, 0.740] | AD $>$ CN | 0.269 [-0.246, 0.828] | 0.265 [-0.242, 0.816] |
| insula_firstorder_Mean | 70–79 | 0.613 [0.506, 0.789] | AD $>$ CN | 0.411 [-0.229, 1.117] | 0.404 [-0.225, 1.097] |
| | 60–69 | 0.543 [0.500, 0.894] | AD $>$ CN | -0.105 [-1.183, 1.364] | -0.100 [-1.123, 1.295] |
| | 80–89 | 0.705 [0.558, 0.836] | AD $>$ CN | 0.775 [0.263, 1.388] | 0.764 [0.259, 1.368] |
| insula_firstorder_Minimum | 70–79 | 0.687 [0.524, 0.849] | AD $>$ CN | 0.737 [0.142, 1.519] | 0.724 [0.139, 1.492] |
| | 60–69 | 0.886 [0.686, 1.000] | AD $>$ CN | 1.445 [0.786, 2.572] | 1.372 [0.746, 2.441] |
| | 80–89 | 0.734 [0.591, 0.863] | AD $<$ CN | -0.879 [-1.529, -0.342] | -0.866 [-1.506, -0.337] |
| insula_glcm_Idn | 70–79 | 0.802 [0.657, 0.919] | AD $<$ CN | -1.328 [-2.202, -0.693] | -1.305 [-2.163, -0.680] |
| | 60–69 | 0.600 [0.500, 0.886] | AD $<$ CN | -0.470 [-1.645, 0.483] | -0.446 [-1.561, 0.459] |
| | 80–89 | 0.745 [0.603, 0.874] | AD $<$ CN | -0.792 [-1.462, -0.242] | -0.781 [-1.441, -0.238] |
| insula_glcm_Idmn | 70–79 | 0.792 [0.645, 0.915] | AD $<$ CN | -1.267 [-2.127, -0.653] | -1.245 [-2.090, -0.641] |
| | 60–69 | 0.743 [0.519, 0.958] | AD $<$ CN | -0.989 [-2.214, -0.085] | -0.938 [-2.102, -0.081] |

#### *4.10 Heschl's Gyrus*

TABLE XXX: Complete AUC, Cohen's $d$, and Hedges' $g$ results for Heschl's gyrus radiomic features.

| Feature | Age bin | AUC [95% CI] | Direction | Cohen's $d$ [95% CI] | Hedges' $g$ [95% CI] |
|---|---|---|---|---|---|
| | 80–89 | 0.748 [0.607, 0.866] | AD $>$ CN | 0.894 [0.379, 1.527] | 0.881 [0.373, 1.505] |
| heschls_gyrus_firstorder_Minimum | 70–79 | 0.726 [0.557, 0.876] | AD $>$ CN | 0.848 [0.273, 1.567] | 0.833 [0.268, 1.540] |
| | 60–69 | 0.900 [0.700, 1.000] | AD $>$ CN | 1.566 [0.854, 3.010] | 1.487 [0.810, 2.857] |
| | 80–89 | 0.571 [0.504, 0.722] | AD $>$ CN | 0.286 [-0.227, 0.848] | 0.282 [-0.224, 0.836] |
| heschls_gyrus_firstorder_10Percentile | 70–79 | 0.660 [0.516, 0.819] | AD $>$ CN | 0.666 [0.074, 1.300] | 0.655 [0.072, 1.277] |
| | 60–69 | 0.686 [0.514, 0.970] | AD $>$ CN | 0.682 [-0.322, 2.259] | 0.647 [-0.306, 2.144] |
| | 80–89 | 0.534 [0.503, 0.694] | AD $<$ CN | -0.102 [-0.679, 0.419] | -0.101 [-0.669, 0.413] |
| heschls_gyrus_glcm_Idmn | 70–79 | 0.591 [0.504, 0.765] | AD $<$ CN | -0.358 [-1.023, 0.237] | -0.352 [-1.005, 0.233] |
| | 60–69 | 0.886 [0.667, 1.000] | AD $<$ CN | -1.355 [-2.384, -0.836] | -1.286 [-2.263, -0.793] |
| | 80–89 | 0.754 [0.607, 0.879] | AD $<$ CN | -0.981 [-1.567, -0.475] | -0.967 [-1.545, -0.469] |
| heschls_gyrus_firstorder_Kurtosis | 70–79 | 0.591 [0.504, 0.775] | AD $<$ CN | -0.357 [-1.069, 0.288] | -0.351 [-1.050, 0.283] |
| | 60–69 | 0.786 [0.538, 1.000] | AD $<$ CN | -1.042 [-2.410, -0.092] | -0.989 [-2.287, -0.087] |
| | 80–89 | 0.695 [0.543, 0.832] | AD $<$ CN | -0.700 [-1.380, -0.176] | -0.689 [-1.360, -0.174] |
| heschls_gyrus_firstorder_Range | 70–79 | 0.737 [0.571, 0.879] | AD $<$ CN | -0.903 [-1.618, -0.306] | -0.887 [-1.589, -0.300] |
| | 60–69 | 0.871 [0.667, 1.000] | AD $<$ CN | -1.803 [-3.525, -0.905] | -1.711 [-3.346, -0.859] |
| | 80–89 | 0.526 [0.503, 0.688] | AD $>$ CN | 0.088 [-0.434, 0.643] | 0.087 [-0.427, 0.634] |
| heschls_gyrus_ngtdm_Contrast | 70–79 | 0.523 [0.502, 0.716] | AD $>$ CN | 0.147 [-0.491, 0.759] | 0.144 [-0.483, 0.746] |
| | 60–69 | 0.900 [0.694, 1.000] | AD $>$ CN | 1.284 [0.894, 2.367] | 1.218 [0.848, 2.246] |
| | 80–89 | 0.662 [0.517, 0.812] | AD $<$ CN | -0.451 [-1.151, 0.090] | -0.445 [-1.135, 0.088] |
| heschls_gyrus_glrlm_RunEntropy | 70–79 | 0.609 [0.506, 0.776] | AD $<$ CN | -0.563 [-1.165, 0.026] | -0.553 [-1.144, 0.026] |
| | 60–69 | 0.743 [0.528, 0.958] | AD $<$ CN | -1.086 [-2.159, -0.266] | -1.031 [-2.049, -0.252] |
| | 80–89 | 0.512 [0.502, 0.680] | AD $<$ CN | -0.056 [-0.608, 0.483] | -0.055 [-0.599, 0.476] |
| heschls_gyrus_glcm_Idn | 70–79 | 0.588 [0.505, 0.763] | AD $<$ CN | -0.303 [-1.000, 0.301] | -0.298 [-0.983, 0.296] |
| | 60–69 | 0.829 [0.591, 1.000] | AD $<$ CN | -1.221 [-2.174, -0.605] | -1.159 [-2.063, -0.574] |
| | 80–89 | 0.690 [0.540, 0.828] | AD $<$ CN | -0.565 [-1.294, -0.036] | -0.557 [-1.275, -0.035] |
| heschls_gyrus_gldm_DependenceEntropy | 70–79 | 0.628 [0.509, 0.793] | AD $<$ CN | -0.467 [-1.075, 0.109] | -0.459 [-1.057, 0.107] |
| | 60–69 | 0.643 [0.500, 0.914] | AD $<$ CN | -0.628 [-1.795, 0.399] | -0.596 [-1.704, 0.378] |
| | 80–89 | 0.601 [0.506, 0.761] | AD $<$ CN | -0.480 [-1.096, 0.055] | -0.473 [-1.081, 0.054] |
| heschls_gyrus_glrlm_LongRunHighGrayLevelEmphasis | 70–79 | 0.578 [0.504, 0.757] | AD $<$ CN | -0.299 [-0.986, 0.291] | -0.293 [-0.968, 0.286] |
| | 60–69 | 0.900 [0.671, 1.000] | AD $<$ CN | -1.709 [-3.910, -0.754] | -1.622 [-3.711, -0.716] |

### 5 LMCI Supplementary Statistics

#### *5.1 Hippocampus*

##### *5.1.1 Normal–LMCI t-test Results:*

TABLE XXXI: Normal–LMCI t-test results for hippocampus radiomic features.

| Feature | Normal bin | LMCI bin | $t$-value | $p$-value | $n$ normal | $n$ LMCI | FDR $p$ | Significant FDR |
|---|---|---|---|---|---|---|---|---|
| | 60–69 | 60–69 | 0.818332 | 0.436956 | 10 | 10 | 0.436956 | False |
| hippocampus_glcm_ClusterShade | 70–79 | 70–79 | 6.335718 | 0.000200 | 27 | 20 | 0.000667 | True |
| | 80–89 | 80–89 | 2.917828 | 0.007599 | 25 | 14 | 0.012665 | True |
| | 60–69 | 60–69 | 1.624304 | 0.122988 | 10 | 10 | 0.131773 | False |
| hippocampus_gldm_GrayLevelNonUniformity | 70–79 | 70–79 | 5.458502 | 0.000200 | 27 | 20 | 0.000667 | True |
| | 80–89 | 80–89 | 3.728086 | 0.001400 | 25 | 14 | 0.004200 | True |
| | 60–69 | 60–69 | 2.301007 | 0.032597 | 10 | 10 | 0.043039 | True |
| hippocampus_firstorder_Uniformity | 70–79 | 70–79 | 5.902989 | 0.000200 | 27 | 20 | 0.000667 | True |
| | 80–89 | 80–89 | 3.188944 | 0.003600 | 25 | 14 | 0.009230 | True |
| | 60–69 | 60–69 | -2.312297 | 0.032997 | 10 | 10 | 0.043039 | True |
| hippocampus_glcm_ClusterTendency | 70–79 | 70–79 | -5.256798 | 0.000200 | 27 | 20 | 0.000667 | True |
| | 80–89 | 80–89 | -2.887913 | 0.004000 | 25 | 14 | 0.009230 | True |
| | 60–69 | 60–69 | -3.217627 | 0.006999 | 10 | 10 | 0.012352 | True |
| hippocampus_glcm_MCC | 70–79 | 70–79 | -3.070118 | 0.003800 | 27 | 20 | 0.009230 | True |
| | 80–89 | 80–89 | -1.389055 | 0.177982 | 25 | 14 | 0.184120 | False |
| | 60–69 | 60–69 | -1.741501 | 0.096990 | 10 | 10 | 0.107767 | False |
| hippocampus_glcm_SumSquares | 70–79 | 70–79 | -5.578908 | 0.000200 | 27 | 20 | 0.000667 | True |
| | 80–89 | 80–89 | -2.889642 | 0.005000 | 25 | 14 | 0.010713 | True |
| | 60–69 | 60–69 | -1.861218 | 0.078792 | 10 | 10 | 0.094551 | False |
| hippocampus_gldm_GrayLevelVariance | 70–79 | 70–79 | -5.540538 | 0.000200 | 27 | 20 | 0.000667 | True |
| | 80–89 | 80–89 | -2.792253 | 0.006599 | 25 | 14 | 0.012352 | True |
| | 60–69 | 60–69 | -2.187074 | 0.044796 | 10 | 10 | 0.055994 | False |
| hippocampus_firstorder_Entropy | 70–79 | 70–79 | -5.260762 | 0.000200 | 27 | 20 | 0.000667 | True |
| | 80–89 | 80–89 | -2.544060 | 0.010999 | 25 | 14 | 0.017367 | True |
| | 60–69 | 60–69 | -1.816285 | 0.094591 | 10 | 10 | 0.107767 | False |
| hippocampus_firstorder_Variance | 70–79 | 70–79 | -4.829862 | 0.000200 | 27 | 20 | 0.000667 | True |
| | 80–89 | 80–89 | -2.508927 | 0.014199 | 25 | 14 | 0.021298 | True |
| | 60–69 | 60–69 | -2.460212 | 0.020798 | 10 | 10 | 0.029711 | True |
| hippocampus_glcm_SumEntropy | 70–79 | 70–79 | -5.335098 | 0.000200 | 27 | 20 | 0.000667 | True |
| | 80–89 | 80–89 | -2.912552 | 0.005799 | 25 | 14 | 0.011599 | True |

###### *5.1.2 LMCI–Alzheimer t-test Results:*

TABLE XXXII: LMCI–Alzheimer t-test results for hippocampus radiomic features.

| Feature | LMCI bin | Alzheimer bin | $t$-value | $p$-value | $n$ LMCI | $n$ Alzheimer | FDR $p$ | Significant FDR |
|---|---|---|---|---|---|---|---|---|
| | 60–69 | 60–69 | 1.752986 | 0.086391 | 10 | 7 | 0.492551 | False |
| hippocampus_glcm_ClusterShade | 70–79 | 70–79 | -0.532279 | 0.621738 | 20 | 18 | 0.703858 | False |
| | 80–89 | 80–89 | 1.008384 | 0.328367 | 14 | 29 | 0.492551 | False |
| | 60–69 | 60–69 | 0.394492 | 0.694531 | 10 | 7 | 0.718480 | False |
| hippocampus_gldm_GrayLevelNonUniformity | 70–79 | 70–79 | 0.724647 | 0.472353 | 20 | 18 | 0.616112 | False |
| | 80–89 | 80–89 | 1.480713 | 0.132387 | 14 | 29 | 0.492551 | False |
| | 60–69 | 60–69 | -1.174663 | 0.263174 | 10 | 7 | 0.492551 | False |
| hippocampus_firstorder_Uniformity | 70–79 | 70–79 | -1.432092 | 0.171983 | 20 | 18 | 0.492551 | False |
| | 80–89 | 80–89 | 1.430541 | 0.150385 | 14 | 29 | 0.492551 | False |
| | 60–69 | 60–69 | 0.482389 | 0.655734 | 10 | 7 | 0.703858 | False |
| hippocampus_glcm_ClusterTendency | 70–79 | 70–79 | 1.012870 | 0.323768 | 20 | 18 | 0.492551 | False |
| | 80–89 | 80–89 | -1.244383 | 0.241176 | 14 | 29 | 0.492551 | False |
| | 60–69 | 60–69 | -0.937938 | 0.345165 | 10 | 7 | 0.493094 | False |
| hippocampus_glcm_MCC | 70–79 | 70–79 | 0.148112 | 0.896710 | 20 | 18 | 0.896710 | False |
| | 80–89 | 80–89 | -2.403627 | 0.021398 | 14 | 29 | 0.492551 | False |
| | 60–69 | 60–69 | 0.504210 | 0.629537 | 10 | 7 | 0.703858 | False |
| hippocampus_glcm_SumSquares | 70–79 | 70–79 | 1.185911 | 0.234777 | 20 | 18 | 0.492551 | False |
| | 80–89 | 80–89 | -1.023864 | 0.324568 | 14 | 29 | 0.492551 | False |
| | 60–69 | 60–69 | 0.528241 | 0.608539 | 10 | 7 | 0.703858 | False |
| hippocampus_gldm_GrayLevelVariance | 70–79 | 70–79 | 1.301176 | 0.213979 | 20 | 18 | 0.492551 | False |
| | 80–89 | 80–89 | -1.119309 | 0.282572 | 14 | 29 | 0.492551 | False |
| | 60–69 | 60–69 | 1.243033 | 0.227177 | 10 | 7 | 0.492551 | False |
| hippocampus_firstorder_Entropy | 70–79 | 70–79 | 1.413446 | 0.157184 | 20 | 18 | 0.492551 | False |
| | 80–89 | 80–89 | -1.142413 | 0.272573 | 14 | 29 | 0.492551 | False |
| | 60–69 | 60–69 | 0.470613 | 0.656934 | 10 | 7 | 0.703858 | False |
| hippocampus_firstorder_Variance | 70–79 | 70–79 | 0.916975 | 0.372563 | 20 | 18 | 0.508040 | False |
| | 80–89 | 80–89 | -1.266610 | 0.227377 | 14 | 29 | 0.492551 | False |
| | 60–69 | 60–69 | 1.314231 | 0.198980 | 10 | 7 | 0.492551 | False |
| hippocampus_glcm_SumEntropy | 70–79 | 70–79 | 1.249848 | 0.223378 | 20 | 18 | 0.492551 | False |
| | 80–89 | 80–89 | -1.124486 | 0.259574 | 14 | 29 | 0.492551 | False |

##### *5.2 Entorhinal Cortex*

###### *5.2.1 Normal–LMCI t-test Results:*

TABLE XXXIII: Normal–LMCI t-test results for entorhinal cortex radiomic features.

| Feature | Normal bin | LMCI bin | $t$-value | $p$-value | $n$ normal | $n$ LMCI | FDR $p$ | Significant FDR |
|---|---|---|---|---|---|---|---|---|
| | 60–69 | 60–69 | 2.075665 | 0.047995 | 10 | 10 | 0.084697 | False |
| entorhinal_cortex_glcm_ClusterShade | 70–79 | 70–79 | 6.022986 | 0.000200 | 27 | 20 | 0.001200 | True |
| | 80–89 | 80–89 | 2.345168 | 0.012799 | 25 | 14 | 0.027854 | True |
| | 60–69 | 60–69 | 0.438153 | 0.676332 | 10 | 10 | 0.724642 | False |
| entorhinal_cortex_firstorder_Skewness | 70–79 | 70–79 | 3.010770 | 0.006399 | 27 | 20 | 0.021331 | True |
| | 80–89 | 80–89 | 1.193735 | 0.270773 | 25 | 14 | 0.324928 | False |
| | 60–69 | 60–69 | -1.924615 | 0.067993 | 10 | 10 | 0.113322 | False |
| entorhinal_cortex_shape_SurfaceVolumeRatio | 70–79 | 70–79 | -3.525245 | 0.000400 | 27 | 20 | 0.001714 | True |
| | 80–89 | 80–89 | -2.710713 | 0.008399 | 25 | 14 | 0.025197 | True |
| | 60–69 | 60–69 | -1.823114 | 0.078192 | 10 | 10 | 0.123461 | False |
| entorhinal_cortex_firstorder_Median | 70–79 | 70–79 | -4.400698 | 0.000200 | 27 | 20 | 0.001200 | True |
| | 80–89 | 80–89 | -4.005116 | 0.000200 | 25 | 14 | 0.001200 | True |
| | 60–69 | 60–69 | -1.655889 | 0.116788 | 10 | 10 | 0.166840 | False |
| entorhinal_cortex_glrlm_ShortRunLowGrayLevelEmphasis | 70–79 | 70–79 | -2.518751 | 0.016398 | 27 | 20 | 0.030747 | True |
| | 80–89 | 80–89 | -0.296647 | 0.787921 | 25 | 14 | 0.815091 | False |
| | 60–69 | 60–69 | 1.508776 | 0.139786 | 10 | 10 | 0.183982 | False |
| entorhinal_cortex_gldm_DependenceNonUniformity | 70–79 | 70–79 | 4.849041 | 0.000400 | 27 | 20 | 0.001714 | True |
| | 80–89 | 80–89 | 4.719966 | 0.000200 | 25 | 14 | 0.001200 | True |
| | 60–69 | 60–69 | 0.602458 | 0.553745 | 10 | 10 | 0.615272 | False |
| entorhinal_cortex_gldm_DependenceEntropy | 70–79 | 70–79 | 0.584255 | 0.548145 | 27 | 20 | 0.615272 | False |
| | 80–89 | 80–89 | 0.199338 | 0.851715 | 25 | 14 | 0.851715 | False |
| | 60–69 | 60–69 | 1.529487 | 0.143786 | 10 | 10 | 0.183982 | False |
| entorhinal_cortex_shape_VoxelVolume | 70–79 | 70–79 | 2.544767 | 0.012599 | 27 | 20 | 0.027854 | True |
| | 80–89 | 80–89 | 2.547334 | 0.015198 | 25 | 14 | 0.030397 | True |
| | 60–69 | 60–69 | 1.526999 | 0.147185 | 10 | 10 | 0.183982 | False |
| entorhinal_cortex_shape_MeshVolume | 70–79 | 70–79 | 2.571174 | 0.011599 | 27 | 20 | 0.027854 | True |
| | 80–89 | 80–89 | 2.564627 | 0.012999 | 25 | 14 | 0.027854 | True |
| | 60–69 | 60–69 | 1.717034 | 0.099990 | 10 | 10 | 0.149985 | False |
| entorhinal_cortex_glrlm_GrayLevelNonUniformity | 70–79 | 70–79 | 4.614355 | 0.000200 | 27 | 20 | 0.001200 | True |
| | 80–89 | 80–89 | 3.687063 | 0.000800 | 25 | 14 | 0.003000 | True |

###### *5.2.2 LMCI–Alzheimer t-test Results:*

TABLE XXXIV: LMCI–Alzheimer t-test results for entorhinal cortex radiomic features.

| Feature | LMCI bin | Alzheimer bin | $t$-value | $p$-value | $n$ LMCI | $n$ Alzheimer | FDR $p$ | Significant FDR |
|---|---|---|---|---|---|---|---|---|
| | 60–69 | 60–69 | 1.493005 | 0.137986 | 10 | 7 | 0.579692 | False |
| entorhinal_cortex_glcm_ClusterShade | 70–79 | 70–79 | -0.512370 | 0.602740 | 20 | 18 | 0.709544 | False |
| | 80–89 | 80–89 | 0.169003 | 0.891711 | 14 | 29 | 0.955404 | False |
| | 60–69 | 60–69 | 3.212405 | 0.003000 | 10 | 7 | 0.089991 | False |
| entorhinal_cortex_firstorder_Skewness | 70–79 | 70–79 | -1.417388 | 0.154585 | 20 | 18 | 0.579692 | False |
| | 80–89 | 80–89 | -2.628277 | 0.017398 | 14 | 29 | 0.130487 | False |
| | 60–69 | 60–69 | 0.053584 | 0.967103 | 10 | 7 | 0.967103 | False |
| entorhinal_cortex_shape_SurfaceVolumeRatio | 70–79 | 70–79 | -0.994920 | 0.316968 | 20 | 18 | 0.593607 | False |
| | 80–89 | 80–89 | -0.486647 | 0.652735 | 14 | 29 | 0.725261 | False |
| | 60–69 | 60–69 | -0.087096 | 0.945105 | 10 | 7 | 0.967103 | False |
| entorhinal_cortex_firstorder_Median | 70–79 | 70–79 | 1.070944 | 0.273173 | 20 | 18 | 0.593607 | False |
| | 80–89 | 80–89 | 1.713123 | 0.082592 | 14 | 29 | 0.495550 | False |
| | 60–69 | 60–69 | -0.876984 | 0.419558 | 10 | 7 | 0.624795 | False |
| entorhinal_cortex_glrlm_ShortRunLowGrayLevelEmphasis | 70–79 | 70–79 | -1.002474 | 0.324768 | 20 | 18 | 0.593607 | False |
| | 80–89 | 80–89 | -2.472997 | 0.015398 | 14 | 29 | 0.130487 | False |
| | 60–69 | 60–69 | 1.151916 | 0.294571 | 10 | 7 | 0.593607 | False |
| entorhinal_cortex_gldm_DependenceNonUniformity | 70–79 | 70–79 | -0.498185 | 0.614939 | 20 | 18 | 0.709544 | False |
| | 80–89 | 80–89 | -0.977948 | 0.356164 | 14 | 29 | 0.593607 | False |
| | 60–69 | 60–69 | -1.101896 | 0.278772 | 10 | 7 | 0.593607 | False |
| entorhinal_cortex_gldm_DependenceEntropy | 70–79 | 70–79 | -1.676589 | 0.108989 | 20 | 18 | 0.544946 | False |
| | 80–89 | 80–89 | -2.556969 | 0.013199 | 14 | 29 | 0.130487 | False |
| | 60–69 | 60–69 | 0.717991 | 0.507949 | 10 | 7 | 0.639936 | False |
| entorhinal_cortex_shape_VoxelVolume | 70–79 | 70–79 | 1.254240 | 0.222178 | 20 | 18 | 0.593607 | False |
| | 80–89 | 80–89 | 0.762222 | 0.436356 | 14 | 29 | 0.624795 | False |
| | 60–69 | 60–69 | 0.699928 | 0.511949 | 10 | 7 | 0.639936 | False |
| entorhinal_cortex_shape_MeshVolume | 70–79 | 70–79 | 1.252110 | 0.220378 | 20 | 18 | 0.593607 | False |
| | 80–89 | 80–89 | 0.755101 | 0.465753 | 14 | 29 | 0.635118 | False |
| | 60–69 | 60–69 | 0.994063 | 0.343566 | 10 | 7 | 0.593607 | False |
| entorhinal_cortex_glrlm_GrayLevelNonUniformity | 70–79 | 70–79 | 0.796308 | 0.437356 | 20 | 18 | 0.624795 | False |
| | 80–89 | 80–89 | 0.954756 | 0.338166 | 14 | 29 | 0.593607 | False |

##### *5.3 Cingulum*

###### *5.3.1 Normal–LMCI t-test Results:*

TABLE XXXV: Normal–LMCI t-test results for cingulum radiomic features.

| Feature | Normal bin | LMCI bin | $t$-value | $p$-value | $n$ normal | $n$ LMCI | FDR $p$ | Significant FDR |
|---|---|---|---|---|---|---|---|---|
| | 60–69 | 60–69 | -2.915121 | 0.012999 | 10 | 10 | 0.035451 | True |
| cingulum_firstorder_Median | 70–79 | 70–79 | -3.387616 | 0.001200 | 27 | 20 | 0.007199 | True |
| | 80–89 | 80–89 | -0.824666 | 0.389161 | 25 | 14 | 0.466993 | False |
| | 60–69 | 60–69 | 2.626779 | 0.014999 | 10 | 10 | 0.037496 | True |
| cingulum_firstorder_Kurtosis | 70–79 | 70–79 | 4.672280 | 0.000200 | 27 | 20 | 0.003000 | True |
| | 80–89 | 80–89 | 1.582868 | 0.123388 | 25 | 14 | 0.185081 | False |
| | 60–69 | 60–69 | -2.550643 | 0.024598 | 10 | 10 | 0.052709 | False |
| cingulum_firstorder_InterquartileRange | 70–79 | 70–79 | -4.372087 | 0.000200 | 27 | 20 | 0.003000 | True |
| | 80–89 | 80–89 | -1.137351 | 0.264374 | 25 | 14 | 0.344835 | False |
| | 60–69 | 60–69 | -2.222377 | 0.037796 | 10 | 10 | 0.075592 | False |
| cingulum_firstorder_RobustMeanAbsoluteDeviation | 70–79 | 70–79 | -4.190221 | 0.000400 | 27 | 20 | 0.004000 | True |
| | 80–89 | 80–89 | -0.919926 | 0.350965 | 25 | 14 | 0.438706 | False |
| | 60–69 | 60–69 | -3.259768 | 0.005399 | 10 | 10 | 0.020248 | True |
| cingulum_firstorder_Mean | 70–79 | 70–79 | -2.777055 | 0.007199 | 27 | 20 | 0.023998 | True |
| | 80–89 | 80–89 | -0.756844 | 0.417158 | 25 | 14 | 0.481336 | False |
| | 60–69 | 60–69 | -2.664057 | 0.020798 | 10 | 10 | 0.047995 | True |
| cingulum_glszm_GrayLevelNonUniformityNormalized | 70–79 | 70–79 | -1.423921 | 0.155184 | 27 | 20 | 0.221692 | False |
| | 80–89 | 80–89 | -0.670915 | 0.515348 | 25 | 14 | 0.572609 | False |
| | 60–69 | 60–69 | -1.740973 | 0.093791 | 10 | 10 | 0.156318 | False |
| cingulum_firstorder_Minimum | 70–79 | 70–79 | -3.239555 | 0.002600 | 27 | 20 | 0.012999 | True |
| | 80–89 | 80–89 | -1.644041 | 0.099790 | 25 | 14 | 0.157563 | False |
| | 60–69 | 60–69 | -3.278674 | 0.003800 | 10 | 10 | 0.016284 | True |
| cingulum_firstorder_90Percentile | 70–79 | 70–79 | -2.680114 | 0.010599 | 27 | 20 | 0.031797 | True |
| | 80–89 | 80–89 | -0.531389 | 0.575142 | 25 | 14 | 0.594975 | False |
| | 60–69 | 60–69 | -1.128004 | 0.262574 | 10 | 10 | 0.344835 | False |
| cingulum_firstorder_MeanAbsoluteDeviation | 70–79 | 70–79 | -3.337387 | 0.001200 | 27 | 20 | 0.007199 | True |
| | 80–89 | 80–89 | -0.398028 | 0.683132 | 25 | 14 | 0.683132 | False |
| | 60–69 | 60–69 | 1.824599 | 0.086791 | 10 | 10 | 0.153161 | False |
| cingulum_firstorder_Energy | 70–79 | 70–79 | 0.637878 | 0.547345 | 27 | 20 | 0.586441 | False |
| | 80–89 | 80–89 | 2.013310 | 0.061394 | 25 | 14 | 0.115113 | False |

##### *5.3.2 LMCI–Alzheimer t-test Results:*

TABLE XXXVI: LMCI–Alzheimer t-test results for cingulum radiomic features.

| Feature | LMCI bin | Alzheimer bin | $t$-value | $p$-value | $n$ LMCI | $n$ Alzheimer | FDR $p$ | Significant FDR |
|---|---|---|---|---|---|---|---|---|
| | 60–69 | 60–69 | 0.227629 | 0.848315 | 10 | 7 | 0.962126 | False |
| cingulum_firstorder_Median | 70–79 | 70–79 | 0.841282 | 0.394561 | 20 | 18 | 0.657601 | False |
| | 80–89 | 80–89 | 0.041688 | 0.955704 | 14 | 29 | 0.980302 | False |
| | 60–69 | 60–69 | 0.652942 | 0.527747 | 10 | 7 | 0.833285 | False |
| cingulum_firstorder_Kurtosis | 70–79 | 70–79 | -0.288224 | 0.788721 | 20 | 18 | 0.962126 | False |
| | 80–89 | 80–89 | 2.525832 | 0.013599 | 14 | 29 | 0.407959 | False |
| | 60–69 | 60–69 | -0.337928 | 0.738126 | 10 | 7 | 0.962126 | False |
| cingulum_firstorder_InterquartileRange | 70–79 | 70–79 | 0.917681 | 0.364564 | 20 | 18 | 0.657601 | False |
| | 80–89 | 80–89 | -1.809754 | 0.079592 | 14 | 29 | 0.657601 | False |
| | 60–69 | 60–69 | -0.331340 | 0.739326 | 10 | 7 | 0.962126 | False |
| cingulum_firstorder_RobustMeanAbsoluteDeviation | 70–79 | 70–79 | 0.989586 | 0.335166 | 20 | 18 | 0.657601 | False |
| | 80–89 | 80–89 | -1.675974 | 0.109589 | 14 | 29 | 0.657601 | False |
| | 60–69 | 60–69 | 0.871131 | 0.392761 | 10 | 7 | 0.657601 | False |
| cingulum_firstorder_Mean | 70–79 | 70–79 | 0.985324 | 0.344566 | 20 | 18 | 0.657601 | False |
| | 80–89 | 80–89 | 0.373814 | 0.715328 | 14 | 29 | 0.962126 | False |
| | 60–69 | 60–69 | -0.998294 | 0.315168 | 10 | 7 | 0.657601 | False |
| cingulum_glszm_GrayLevelNonUniformityNormalized | 70–79 | 70–79 | -1.353243 | 0.177982 | 20 | 18 | 0.657601 | False |
| | 80–89 | 80–89 | -2.330803 | 0.038796 | 14 | 29 | 0.581942 | False |
| | 60–69 | 60–69 | -1.434068 | 0.196980 | 10 | 7 | 0.657601 | False |
| cingulum_firstorder_Minimum | 70–79 | 70–79 | 0.488265 | 0.627937 | 20 | 18 | 0.941906 | False |
| | 80–89 | 80–89 | -1.219165 | 0.238976 | 14 | 29 | 0.657601 | False |
| | 60–69 | 60–69 | 1.110251 | 0.264174 | 10 | 7 | 0.657601 | False |
| cingulum_firstorder_90Percentile | 70–79 | 70–79 | 1.393625 | 0.170583 | 20 | 18 | 0.657601 | False |
| | 80–89 | 80–89 | 0.173367 | 0.865913 | 14 | 29 | 0.962126 | False |
| | 60–69 | 60–69 | -0.020579 | 0.980302 | 10 | 7 | 0.980302 | False |
| cingulum_firstorder_MeanAbsoluteDeviation | 70–79 | 70–79 | 1.149574 | 0.257974 | 20 | 18 | 0.657601 | False |
| | 80–89 | 80–89 | -0.999869 | 0.333767 | 14 | 29 | 0.657601 | False |
| | 60–69 | 60–69 | -0.027147 | 0.978502 | 10 | 7 | 0.980302 | False |
| cingulum_firstorder_Energy | 70–79 | 70–79 | -0.249075 | 0.810519 | 20 | 18 | 0.962126 | False |
| | 80–89 | 80–89 | -1.316571 | 0.206379 | 14 | 29 | 0.657601 | False |

#### 6 Female Results

##### 6.1 *Spearman Correlation Results*

###### 6.1.1 *Caudate Nucleus:*

Table XXXVII. Female subgroup Spearman correlation results for caudate nucleus radiomic features.

| Radiomic feature | Spearman $\rho$ | $p$-value | FDR $p$ | Significant |
|---|---|---|---|---|
| caudate_nucleus_glrlm_RunPercentage | 0.520362 | 0.000047 | 0.000747 | True |
| caudate_nucleus_gldm_LargeDependenceEmphasis | -0.519855 | 0.000048 | 0.000747 | True |
| caudate_nucleus_glcm_InverseVariance | 0.511827 | 0.000065 | 0.000747 | True |
| caudate_nucleus_glcm_DifferenceEntropy | 0.510453 | 0.000068 | 0.000747 | True |
| caudate_nucleus_glrlm_RunLengthNonUniformityNormalized | 0.509115 | 0.000072 | 0.000747 | True |
| caudate_nucleus_glcm_DifferenceVariance | 0.508572 | 0.000074 | 0.000747 | True |
| caudate_nucleus_glcm_Id | -0.507777 | 0.000076 | 0.000747 | True |
| caudate_nucleus_glrlm_LongRunEmphasis | -0.507704 | 0.000076 | 0.000747 | True |
| caudate_nucleus_glrlm_ShortRunEmphasis | 0.507596 | 0.000076 | 0.000747 | True |
| caudate_nucleus_glcm_Contrast | 0.505715 | 0.000082 | 0.000747 | True |

###### 6.1.2 *Entorhinal Cortex:*

Table XXXVIII. Female subgroup Spearman correlation results for entorhinal cortex radiomic features.

| Radiomic feature | Spearman $\rho$ | $p$-value | FDR $p$ | Significant |
|---|---|---|---|---|
| entorhinal_cortex_glcm_Idmn | -0.549691 | 0.000014 | 0.001125 | True |
| entorhinal_cortex_firstorder_Skewness | -0.535117 | 0.000026 | 0.001125 | True |
| entorhinal_cortex_glrlm_ShortRunLowGrayLevelEmphasis | 0.527341 | 0.000035 | 0.001125 | True |
| entorhinal_cortex_glcm_Idn | -0.522929 | 0.000042 | 0.001125 | True |
| entorhinal_cortex_ngtdm_Contrast | 0.516167 | 0.000055 | 0.001175 | True |
| entorhinal_cortex_glszm_HighGrayLevelZoneEmphasis | -0.494939 | 0.000122 | 0.001691 | True |
| entorhinal_cortex_firstorder_Range | -0.494577 | 0.000124 | 0.001691 | True |
| entorhinal_cortex_glrlm_HighGrayLevelRunEmphasis | -0.494034 | 0.000126 | 0.001691 | True |
| entorhinal_cortex_glszm_LowGrayLevelZoneEmphasis | 0.475446 | 0.000244 | 0.002672 | True |
| entorhinal_cortex_ngtdm_Strength | -0.474795 | 0.000250 | 0.002672 | True |

###### 6.1.3 *Hippocampus:*

Table XXXIX. Female subgroup Spearman correlation results for hippocampus radiomic features.

| Radiomic feature | Spearman $\rho$ | $p$-value | FDR $p$ | Significant |
|---|---|---|---|---|
| hippocampus_gldm_GrayLevelNonUniformity | -0.572185 | 0.000005 | 0.000236 | True |
| hippocampus_firstorder_Uniformity | -0.559346 | 0.000009 | 0.000236 | True |
| hippocampus_glcm_JointEnergy | -0.556309 | 0.000010 | 0.000236 | True |
| hippocampus_glcm_ClusterShade | -0.553379 | 0.000012 | 0.000236 | True |
| hippocampus_glrlm_RunVariance | -0.550450 | 0.000013 | 0.000236 | True |
| hippocampus_gldm_DependenceVariance | -0.547593 | 0.000015 | 0.000236 | True |
| hippocampus_glcm_MaximumProbability | -0.546798 | 0.000016 | 0.000236 | True |
| hippocampus_firstorder_Median | 0.543977 | 0.000018 | 0.000236 | True |
| hippocampus_firstorder_Skewness | -0.532260 | 0.000029 | 0.000343 | True |
| hippocampus_glrlm_LongRunEmphasis | -0.525967 | 0.000037 | 0.000398 | True |

###### 6.1.4 *Heschl's Gyrus:*

Table XL. Female subgroup Spearman correlation results for Heschl's gyrus radiomic features.

| Radiomic feature | Spearman $\rho$ | $p$-value | FDR $p$ | Significant |
|---|---|---|---|---|
| heschls_gyrus_firstorder_Median | 0.509332 | 0.000071 | 0.007648 | True |
| heschls_gyrus_firstorder_Mean | 0.468792 | 0.000306 | 0.011792 | True |
| heschls_gyrus_firstorder_Minimum | 0.466514 | 0.000331 | 0.011792 | True |
| heschls_gyrus_gldm_DependenceEntropy | -0.426444 | 0.001168 | 0.028772 | True |
| heschls_gyrus_firstorder_90Percentile | 0.421671 | 0.001344 | 0.028772 | True |
| heschls_gyrus_glrlm_RunEntropy | -0.405795 | 0.002114 | 0.037697 | True |
| heschls_gyrus_firstorder_Kurtosis | -0.389015 | 0.003332 | 0.050837 | False |
| heschls_gyrus_glszm_GrayLevelNonUniformityNormalized | 0.378599 | 0.004369 | 0.050837 | False |
| heschls_gyrus_firstorder_TotalEnergy | -0.375309 | 0.004751 | 0.050837 | False |
| heschls_gyrus_firstorder_Energy | -0.375309 | 0.004751 | 0.050837 | False |

###### 6.1.5 *Insula:*

Table XLI. Female subgroup Spearman correlation results for insula radiomic features.

| Radiomic feature | Spearman $\rho$ | $p$-value | FDR $p$ | Significant |
|---|---|---|---|---|
| insula_firstorder_Median | 0.640100 | $1.424444 \times 10^{-7}$ | 0.000015 | True |
| insula_firstorder_Mean | 0.574535 | $4.498389 \times 10^{-6}$ | 0.000241 | True |
| insula_firstorder_RootMeanSquared | -0.547376 | $1.527327 \times 10^{-5}$ | 0.000545 | True |
| insula_firstorder_Minimum | 0.486549 | $1.655825 \times 10^{-4}$ | 0.004429 | True |
| insula_firstorder_Energy | -0.466550 | $3.302288 \times 10^{-4}$ | 0.005889 | True |
| insula_firstorder_TotalEnergy | -0.466550 | $3.302288 \times 10^{-4}$ | 0.005889 | True |
| insula_firstorder_90Percentile | 0.424130 | $1.250983 \times 10^{-3}$ | 0.018366 | True |
| insula_firstorder_Skewness | -0.420947 | $1.373125 \times 10^{-3}$ | 0.018366 | True |
| insula_firstorder_10Percentile | 0.385760 | $3.629816 \times 10^{-3}$ | 0.043154 | True |
| insula_ngtdm_Strength | -0.381601 | $4.044394 \times 10^{-3}$ | 0.043275 | True |

###### 6.1.6 *Putamen:*

Table XLII. Female subgroup Spearman correlation results for putamen radiomic features.

| Radiomic feature | Spearman $\rho$ | $p$-value | FDR $p$ | Significant |
|---|---|---|---|---|
| putamen_firstorder_90Percentile | 0.518879 | 0.000049 | 0.002200 | True |
| putamen_firstorder_Median | 0.516348 | 0.000055 | 0.002200 | True |
| putamen_glcm_ClusterShade | -0.511936 | 0.000065 | 0.002200 | True |
| putamen_firstorder_Mean | 0.505643 | 0.000082 | 0.002200 | True |
| putamen_shape_Flatness | -0.438053 | 0.000823 | 0.017619 | True |
| putamen_firstorder_10Percentile | 0.419682 | 0.001425 | 0.025405 | True |
| putamen_shape_MinorAxisLength | 0.381782 | 0.004026 | 0.061533 | False |
| putamen_glrlm_ShortRunEmphasis | -0.373356 | 0.004991 | 0.066760 | False |
| putamen_shape_Maximum2DDiameterSlice | 0.356649 | 0.007523 | 0.089444 | False |
| putamen_glrlm_RunLengthNonUniformityNormalized | -0.347462 | 0.009344 | 0.099983 | False |

###### 6.1.7 *Rolandic Operculum:*

Table XLIII. Female subgroup Spearman correlation results for rolandic operculum radiomic features.

| Radiomic feature | Spearman $\rho$ | $p$-value | FDR $p$ | Significant |
|---|---|---|---|---|
| rolandic_operculum_firstorder_Median | 0.558840 | 0.000009 | 0.000622 | True |
| rolandic_operculum_firstorder_Minimum | 0.539926 | 0.000021 | 0.000622 | True |
| rolandic_operculum_firstorder_Energy | -0.537467 | 0.000023 | 0.000622 | True |
| rolandic_operculum_firstorder_TotalEnergy | -0.537467 | 0.000023 | 0.000622 | True |
| rolandic_operculum_firstorder_Mean | 0.513671 | 0.000060 | 0.001198 | True |
| rolandic_operculum_glcm_MaximumProbability | -0.510959 | 0.000067 | 0.001198 | True |
| rolandic_operculum_firstorder_RootMeanSquared | -0.469407 | 0.000300 | 0.004586 | True |
| rolandic_operculum_firstorder_Range | -0.462680 | 0.000376 | 0.005024 | True |
| rolandic_operculum_glszm_GrayLevelVariance | -0.400985 | 0.002414 | 0.028701 | True |
| rolandic_operculum_firstorder_90Percentile | 0.389738 | 0.003269 | 0.034977 | True |

###### 6.1.8 *Thalamus:*

Table XLIV. Female subgroup Spearman correlation results for thalamus radiomic features.

| Radiomic feature | Spearman $\rho$ | $p$-value | FDR $p$ | Significant |
|---|---|---|---|---|
| thalamus_shape_LeastAxisLength | -0.557068 | 0.000010 | 0.001070 | True |
| thalamus_shape_Sphericity | -0.452627 | 0.000521 | 0.022988 | True |
| thalamus_shape_Flatness | -0.445937 | 0.000645 | 0.022988 | True |
| thalamus_firstorder_Minimum | 0.412087 | 0.001771 | 0.047384 | True |
| thalamus_shape_SurfaceVolumeRatio | 0.402793 | 0.002297 | 0.049157 | True |
| thalamus_firstorder_RootMeanSquared | 0.366123 | 0.005977 | 0.106592 | False |
| thalamus_firstorder_TotalEnergy | 0.335275 | 0.012341 | 0.165060 | False |
| thalamus_firstorder_Energy | 0.335275 | 0.012341 | 0.165060 | False |
| thalamus_firstorder_Variance | 0.325619 | 0.015270 | 0.171380 | False |
| thalamus_glcm_ClusterProminence | 0.322690 | 0.016269 | 0.171380 | False |

###### 6.1.9 *Vermis:*

Table XLV. Female subgroup Spearman correlation results for vermis radiomic features.

| Radiomic feature | Spearman $\rho$ | $p$-value | FDR $p$ | Significant |
|---|---|---|---|---|
| vermis_firstorder_Kurtosis | -0.345329 | 0.009818 | 0.332802 | False |
| vermis_glcm_Imc2 | 0.343412 | 0.010261 | 0.332802 | False |
| vermis_glcm_MCC | 0.336975 | 0.011879 | 0.332802 | False |
| vermis_glcm_Correlation | 0.334913 | 0.012441 | 0.332802 | False |
| vermis_gldm_DependenceEntropy | 0.306525 | 0.022838 | 0.465439 | False |
| vermis_glrlm_RunEntropy | 0.296110 | 0.028160 | 0.465439 | False |
| vermis_glcm_Imc1 | -0.292132 | 0.030449 | 0.465439 | False |
| vermis_shape_MeshVolume | 0.218394 | 0.109189 | 0.926170 | False |
| vermis_shape_VoxelVolume | 0.215392 | 0.114267 | 0.926170 | False |
| vermis_firstorder_Minimum | 0.213041 | 0.118369 | 0.926170 | False |

###### 6.1.10 *Cingulum:*

Table XLVI. Female subgroup Spearman correlation results for cingulum radiomic features.

| Radiomic feature | Spearman $\rho$ | $p$-value | FDR $p$ | Significant |
|---|---|---|---|---|
| cingulum_firstorder_Kurtosis | -0.597463 | 0.000001 | 0.000126 | True |
| cingulum_firstorder_Median | 0.587952 | 0.000002 | 0.000126 | True |
| cingulum_firstorder_Mean | 0.509802 | 0.000070 | 0.002504 | True |
| cingulum_firstorder_Minimum | 0.456894 | 0.000454 | 0.010208 | True |
| cingulum_firstorder_RobustMeanAbsoluteDeviation | 0.455375 | 0.000477 | 0.010208 | True |
| cingulum_firstorder_RootMeanSquared | -0.438993 | 0.000800 | 0.013117 | True |
| cingulum_firstorder_TotalEnergy | -0.427855 | 0.001120 | 0.013117 | True |
| cingulum_firstorder_Energy | -0.427855 | 0.001120 | 0.013117 | True |
| cingulum_firstorder_InterquartileRange | 0.426372 | 0.001171 | 0.013117 | True |
| cingulum_glszm_GrayLevelNonUniformityNormalized | 0.424817 | 0.001226 | 0.013117 | True |

##### 6.2 *T-test Results*

###### 6.2.1 *Cingulum:*

Table XLVII. Female subgroup t-test results for cingulum radiomic features by age bin.

| Feature | Normal bin | AD bin | $t$-stat | $p$-value | $n$ normal | $n$ AD | FDR $p$ | Significant FDR |
|---|---|---|---|---|---|---|---|---|
| | 60–69 | 60–69 | 4.598171 | 0.000400 | 14 | 7 | 0.004000 | True |
| cingulum_firstorder_Kurtosis | 70–79 | 70–79 | 4.842073 | 0.000200 | 29 | 25 | 0.004000 | True |
| | 80–89 | 80–89 | 2.709591 | 0.011999 | 11 | 7 | 0.044996 | True |
| | 60–69 | 60–69 | -0.228681 | 0.842516 | 14 | 7 | 0.907409 | False |
| cingulum_firstorder_Median | 70–79 | 70–79 | -2.228879 | 0.030997 | 29 | 25 | 0.092991 | False |
| | 80–89 | 80–89 | 0.595827 | 0.569143 | 11 | 7 | 0.761693 | False |
| | 60–69 | 60–69 | 0.023216 | 0.991501 | 14 | 7 | 0.991501 | False |
| cingulum_firstorder_Mean | 70–79 | 70–79 | -1.779543 | 0.081392 | 29 | 25 | 0.187827 | False |
| | 80–89 | 80–89 | 0.573210 | 0.582342 | 11 | 7 | 0.761693 | False |
| | 60–69 | 60–69 | -2.334179 | 0.040396 | 14 | 7 | 0.110171 | False |
| cingulum_firstorder_Minimum | 70–79 | 70–79 | -3.573064 | 0.001000 | 29 | 25 | 0.005999 | True |
| | 80–89 | 80–89 | -1.718162 | 0.104390 | 11 | 7 | 0.223692 | False |
| | 60–69 | 60–69 | -1.564189 | 0.116788 | 14 | 7 | 0.233577 | False |
| cingulum_firstorder_RobustMeanAbsoluteDeviation | 70–79 | 70–79 | -3.975616 | 0.000600 | 29 | 25 | 0.004500 | True |
| | 80–89 | 80–89 | -0.438683 | 0.656334 | 11 | 7 | 0.761693 | False |
| | 60–69 | 60–69 | -0.227061 | 0.846915 | 14 | 7 | 0.907409 | False |
| cingulum_firstorder_RootMeanSquared | 70–79 | 70–79 | 0.701796 | 0.488351 | 29 | 25 | 0.761693 | False |
| | 80–89 | 80–89 | 0.006326 | 0.990901 | 11 | 7 | 0.991501 | False |
| | 60–69 | 60–69 | -0.510479 | 0.617138 | 14 | 7 | 0.761693 | False |
| cingulum_firstorder_TotalEnergy | 70–79 | 70–79 | 0.425198 | 0.660134 | 29 | 25 | 0.761693 | False |
| | 80–89 | 80–89 | 0.566561 | 0.576342 | 11 | 7 | 0.761693 | False |
| | 60–69 | 60–69 | -0.510478 | 0.616338 | 14 | 7 | 0.761693 | False |
| cingulum_firstorder_Energy | 70–79 | 70–79 | 0.425198 | 0.651735 | 29 | 25 | 0.761693 | False |
| | 80–89 | 80–89 | 0.566561 | 0.574943 | 11 | 7 | 0.761693 | False |
| | 60–69 | 60–69 | -1.864359 | 0.061994 | 14 | 7 | 0.154985 | False |
| cingulum_firstorder_InterquartileRange | 70–79 | 70–79 | -3.755177 | 0.000400 | 29 | 25 | 0.004000 | True |
| | 80–89 | 80–89 | -0.922258 | 0.356364 | 11 | 7 | 0.668183 | False |
| | 60–69 | 60–69 | -2.910605 | 0.002200 | 14 | 7 | 0.010999 | True |
| cingulum_glszm_GrayLevelNonUniformityNormalized | 70–79 | 70–79 | -2.301246 | 0.017998 | 29 | 25 | 0.059994 | False |
| | 80–89 | 80–89 | -2.891860 | 0.011399 | 11 | 7 | 0.044996 | True |

##### *6.2.2 Caudate Nucleus:*

Table XLVIII. Female subgroup t-test results for caudate nucleus radiomic features by age bin.

| Feature | Normal bin | AD bin | $t$-stat | $p$-value | $n$ normal | $n$ AD | FDR $p$ | Significant FDR |
|---|---|---|---|---|---|---|---|---|
| | 60–69 | 60–69 | 1.017863 | 0.318968 | 14 | 7 | 0.478452 | False |
| caudate_nucleus_glrlm_RunPercentage | 70–79 | 70–79 | 0.095055 | 0.921708 | 29 | 25 | 0.953491 | False |
| | 80–89 | 80–89 | 1.341294 | 0.208779 | 11 | 7 | 0.424899 | False |
| | 60–69 | 60–69 | -1.042654 | 0.313969 | 14 | 7 | 0.478452 | False |
| caudate_nucleus_gldm_LargeDependenceEmphasis | 70–79 | 70–79 | -0.262649 | 0.804920 | 29 | 25 | 0.928574 | False |
| | 80–89 | 80–89 | -1.293312 | 0.223978 | 11 | 7 | 0.424899 | False |
| | 60–69 | 60–69 | 1.482005 | 0.158984 | 14 | 7 | 0.424899 | False |
| caudate_nucleus_glcm_InverseVariance | 70–79 | 70–79 | 0.428659 | 0.658534 | 29 | 25 | 0.866913 | False |
| | 80–89 | 80–89 | 1.441570 | 0.161984 | 11 | 7 | 0.424899 | False |
| | 60–69 | 60–69 | 1.304683 | 0.202980 | 14 | 7 | 0.424899 | False |
| caudate_nucleus_glcm_DifferenceEntropy | 70–79 | 70–79 | 0.175269 | 0.835716 | 29 | 25 | 0.928574 | False |
| | 80–89 | 80–89 | 1.502159 | 0.155584 | 11 | 7 | 0.424899 | False |
| | 60–69 | 60–69 | 1.151776 | 0.281772 | 14 | 7 | 0.469620 | False |
| caudate_nucleus_glrlm_ RunLengthNonUniformityNormalized | 70–79 | 70–79 | -0.418606 | 0.679932 | 29 | 25 | 0.866913 | False |
| | 80–89 | 80–89 | 1.535419 | 0.141786 | 11 | 7 | 0.424899 | False |
| | 60–69 | 60–69 | 1.252948 | 0.232977 | 14 | 7 | 0.424899 | False |
| caudate_nucleus_glcm_DifferenceVariance | 70–79 | 70–79 | 0.064661 | 0.956304 | 29 | 25 | 0.956304 | False |
| | 80–89 | 80–89 | 1.444764 | 0.174383 | 11 | 7 | 0.424899 | False |
| | 60–69 | 60–69 | -1.424109 | 0.181382 | 14 | 7 | 0.424899 | False |
| caudate_nucleus_glcm_Id | 70–79 | 70–79 | -0.342929 | 0.727727 | 29 | 25 | 0.873273 | False |
| | 80–89 | 80–89 | -1.393209 | 0.197180 | 11 | 7 | 0.424899 | False |
| | 60–69 | 60–69 | -1.238377 | 0.193381 | 14 | 7 | 0.424899 | False |
| caudate_nucleus_glrlm_LongRunEmphasis | 70–79 | 70–79 | -0.544779 | 0.576742 | 29 | 25 | 0.823918 | False |
| | 80–89 | 80–89 | -1.566216 | 0.111589 | 11 | 7 | 0.424899 | False |
| | 60–69 | 60–69 | 1.266161 | 0.210979 | 14 | 7 | 0.424899 | False |
| caudate_nucleus_glrlm_ShortRunEmphasis | 70–79 | 70–79 | -0.400033 | 0.693531 | 29 | 25 | 0.866913 | False |
| | 80–89 | 80–89 | 1.669993 | 0.116988 | 11 | 7 | 0.424899 | False |
| | 60–69 | 60–69 | 1.300034 | 0.240776 | 14 | 7 | 0.424899 | False |
| caudate_nucleus_glcm_Contrast | 70–79 | 70–79 | 0.142085 | 0.871113 | 29 | 25 | 0.933335 | False |
| | 80–89 | 80–89 | 1.277778 | 0.217778 | 11 | 7 | 0.424899 | False |

##### *6.2.3 Thalamus:*

Table XLIX. Female subgroup t-test results for thalamus radiomic features by age bin.

| Feature | Normal bin | AD bin | $t$-stat | $p$-value | $n$ normal | $n$ AD | FDR $p$ | Significant FDR |
|---|---|---|---|---|---|---|---|---|
| | 60–69 | 60–69 | 1.125657 | 0.267973 | 14 | 7 | 0.472894 | False |
| thalamus_shape_LeastAxisLength | 70–79 | 70–79 | 1.741626 | 0.087791 | 29 | 25 | 0.191552 | False |
| | 80–89 | 80–89 | -2.017226 | 0.059994 | 11 | 7 | 0.163620 | False |
| | 60–69 | 60–69 | 2.142030 | 0.059794 | 14 | 7 | 0.163620 | False |
| thalamus_shape_Sphericity | 70–79 | 70–79 | 4.048918 | 0.000600 | 29 | 25 | 0.014999 | True |
| | 80–89 | 80–89 | -0.208673 | 0.838316 | 11 | 7 | 0.838316 | False |
| | 60–69 | 60–69 | 1.850609 | 0.113589 | 14 | 7 | 0.227177 | False |
| thalamus_shape_Flatness | 70–79 | 70–79 | 3.655644 | 0.001000 | 29 | 25 | 0.014999 | True |
| | 80–89 | 80–89 | -0.829575 | 0.426157 | 11 | 7 | 0.639236 | False |
| | 60–69 | 60–69 | -1.918248 | 0.089391 | 14 | 7 | 0.191552 | False |
| thalamus_firstorder_Minimum | 70–79 | 70–79 | -2.677458 | 0.008999 | 29 | 25 | 0.033747 | True |
| | 80–89 | 80–89 | -0.761321 | 0.453955 | 11 | 7 | 0.648507 | False |
| | 60–69 | 60–69 | -1.989232 | 0.067193 | 14 | 7 | 0.167983 | False |
| thalamus_shape_SurfaceVolumeRatio | 70–79 | 70–79 | -2.556622 | 0.014599 | 29 | 25 | 0.048662 | True |
| | 80–89 | 80–89 | -1.597857 | 0.143786 | 11 | 7 | 0.269598 | False |
| | 60–69 | 60–69 | -0.368808 | 0.684932 | 14 | 7 | 0.811633 | False |
| thalamus_firstorder_RootMeanSquared | 70–79 | 70–79 | -2.820937 | 0.005399 | 29 | 25 | 0.023998 | True |
| | 80–89 | 80–89 | 0.613478 | 0.571343 | 11 | 7 | 0.779104 | False |
| | 60–69 | 60–69 | -0.305030 | 0.739926 | 14 | 7 | 0.811633 | False |
| thalamus_firstorder_TotalEnergy | 70–79 | 70–79 | -2.814031 | 0.005399 | 29 | 25 | 0.023998 | True |
| | 80–89 | 80–89 | 0.924250 | 0.397160 | 11 | 7 | 0.638147 | False |
| | 60–69 | 60–69 | -0.305030 | 0.736726 | 14 | 7 | 0.811633 | False |
| thalamus_firstorder_Energy | 70–79 | 70–79 | -2.814031 | 0.005599 | 29 | 25 | 0.023998 | True |
| | 80–89 | 80–89 | 0.924250 | 0.404160 | 11 | 7 | 0.638147 | False |
| | 60–69 | 60–69 | -0.280249 | 0.756724 | 14 | 7 | 0.811633 | False |
| thalamus_firstorder_Variance | 70–79 | 70–79 | -3.386985 | 0.001800 | 29 | 25 | 0.016498 | True |
| | 80–89 | 80–89 | -0.207199 | 0.815318 | 11 | 7 | 0.838316 | False |
| | 60–69 | 60–69 | -0.485516 | 0.603140 | 14 | 7 | 0.786704 | False |
| thalamus_glcm_ClusterProminence | 70–79 | 70–79 | -3.236553 | 0.002200 | 29 | 25 | 0.016498 | True |
| | 80–89 | 80–89 | 0.312449 | 0.757524 | 11 | 7 | 0.811633 | False |

##### *6.2.4 Rolandic Operculum:*

Table L. Female subgroup t-test results for rolandic operculum radiomic features by age bin.

| Feature | Normal bin | AD bin | $t$-stat | $p$-value | $n$ normal | $n$ AD | FDR $p$ | Significant FDR |
|---|---|---|---|---|---|---|---|---|
| | 60–69 | 60–69 | -2.062818 | 0.041996 | 14 | 7 | 0.126844 | False |
| rolandic_operculum_firstorder_Median | 70–79 | 70–79 | -1.762097 | 0.080392 | 29 | 25 | 0.160784 | False |
| | 80–89 | 80–89 | 0.833178 | 0.430157 | 11 | 7 | 0.679195 | False |
| | 60–69 | 60–69 | -2.461982 | 0.030997 | 14 | 7 | 0.126844 | False |
| rolandic_operculum_firstorder_Minimum | 70–79 | 70–79 | -3.671598 | 0.001400 | 29 | 25 | 0.020998 | True |
| | 80–89 | 80–89 | -0.073422 | 0.944506 | 11 | 7 | 0.977075 | False |
| | 60–69 | 60–69 | 1.949608 | 0.058794 | 14 | 7 | 0.126844 | False |
| rolandic_operculum_firstorder_Energy | 70–79 | 70–79 | 2.068891 | 0.048595 | 29 | 25 | 0.126844 | False |
| | 80–89 | 80–89 | 0.292216 | 0.773523 | 11 | 7 | 0.915980 | False |
| | 60–69 | 60–69 | 1.949608 | 0.050995 | 14 | 7 | 0.126844 | False |
| rolandic_operculum_firstorder_TotalEnergy | 70–79 | 70–79 | 2.068891 | 0.045995 | 29 | 25 | 0.126844 | False |
| | 80–89 | 80–89 | 0.292216 | 0.757924 | 11 | 7 | 0.915980 | False |
| | 60–69 | 60–69 | -2.148109 | 0.040596 | 14 | 7 | 0.126844 | False |
| rolandic_operculum_firstorder_Mean | 70–79 | 70–79 | -1.513108 | 0.128187 | 29 | 25 | 0.240351 | False |
| | 80–89 | 80–89 | 0.310773 | 0.805719 | 11 | 7 | 0.915980 | False |
| | 60–69 | 60–69 | 1.290652 | 0.206179 | 14 | 7 | 0.363846 | False |
| rolandic_operculum_glcm_MaximumProbability | 70–79 | 70–79 | -0.040816 | 0.980702 | 29 | 25 | 0.980702 | False |
| | 80–89 | 80–89 | -0.496084 | 0.629137 | 11 | 7 | 0.857914 | False |
| | 60–69 | 60–69 | 2.013551 | 0.045995 | 14 | 7 | 0.126844 | False |
| rolandic_operculum_firstorder_RootMeanSquared | 70–79 | 70–79 | 2.226225 | 0.031197 | 29 | 25 | 0.126844 | False |
| | 80–89 | 80–89 | 0.181312 | 0.847115 | 11 | 7 | 0.915980 | False |
| | 60–69 | 60–69 | 2.828292 | 0.017398 | 14 | 7 | 0.126844 | False |
| rolandic_operculum_firstorder_Range | 70–79 | 70–79 | 3.112421 | 0.002400 | 29 | 25 | 0.023998 | True |
| | 80–89 | 80–89 | 0.174971 | 0.854915 | 11 | 7 | 0.915980 | False |
| | 60–69 | 60–69 | 1.065001 | 0.297770 | 14 | 7 | 0.496284 | False |
| rolandic_operculum_glszm_GrayLevelVariance | 70–79 | 70–79 | 4.280063 | 0.000200 | 29 | 25 | 0.005999 | True |
| | 80–89 | 80–89 | -0.658597 | 0.512549 | 11 | 7 | 0.768823 | False |
| | 60–69 | 60–69 | -1.899952 | 0.059194 | 14 | 7 | 0.126844 | False |
| rolandic_operculum_firstorder_90Percentile | 70–79 | 70–79 | -0.488899 | 0.628137 | 29 | 25 | 0.857914 | False |
| | 80–89 | 80–89 | -0.295704 | 0.767323 | 11 | 7 | 0.915980 | False |

##### *6.2.5 Vermis:*

Table LI. Female subgroup t-test results for vermis radiomic features by age bin.

| Feature | Normal bin | AD bin | $t$-stat | $p$-value | $n$ normal | $n$ AD | FDR $p$ | Significant FDR |
|---|---|---|---|---|---|---|---|---|
| | 60–69 | 60–69 | -0.288302 | 0.764524 | 14 | 7 | 0.917216 | False |
| vermis_firstorder_Kurtosis | 70–79 | 70–79 | 1.354781 | 0.180782 | 29 | 25 | 0.640877 | False |
| | 80–89 | 80–89 | -1.230887 | 0.202780 | 11 | 7 | 0.640877 | False |
| | 60–69 | 60–69 | -0.423150 | 0.660134 | 14 | 7 | 0.883999 | False |
| vermis_glcm_Imc2 | 70–79 | 70–79 | -0.911591 | 0.350765 | 29 | 25 | 0.640877 | False |
| | 80–89 | 80–89 | 0.282433 | 0.794921 | 11 | 7 | 0.917216 | False |
| | 60–69 | 60–69 | -0.482120 | 0.622938 | 14 | 7 | 0.883999 | False |
| vermis_glcm_MCC | 70–79 | 70–79 | -0.931966 | 0.353365 | 29 | 25 | 0.640877 | False |
| | 80–89 | 80–89 | -0.134956 | 0.890311 | 11 | 7 | 0.953905 | False |
| | 60–69 | 60–69 | -0.408769 | 0.677732 | 14 | 7 | 0.883999 | False |
| vermis_glcm_Correlation | 70–79 | 70–79 | -0.930211 | 0.363164 | 29 | 25 | 0.640877 | False |
| | 80–89 | 80–89 | 0.195296 | 0.831517 | 11 | 7 | 0.923908 | False |
| | 60–69 | 60–69 | 1.734278 | 0.113589 | 14 | 7 | 0.640877 | False |
| vermis_gldm_DependenceEntropy | 70–79 | 70–79 | 1.009446 | 0.328567 | 29 | 25 | 0.640877 | False |
| | 80–89 | 80–89 | 1.426131 | 0.159984 | 11 | 7 | 0.640877 | False |
| | 60–69 | 60–69 | 0.936066 | 0.430157 | 14 | 7 | 0.716928 | False |
| vermis_glrlm_RunEntropy | 70–79 | 70–79 | 0.274530 | 0.774523 | 29 | 25 | 0.917216 | False |
| | 80–89 | 80–89 | 1.826131 | 0.099590 | 11 | 7 | 0.640877 | False |
| | 60–69 | 60–69 | 0.994215 | 0.289571 | 14 | 7 | 0.640877 | False |
| vermis_glcm_Imc1 | 70–79 | 70–79 | 1.206400 | 0.219778 | 29 | 25 | 0.640877 | False |
| | 80–89 | 80–89 | 1.004815 | 0.311169 | 11 | 7 | 0.640877 | False |
| | 60–69 | 60–69 | -0.440764 | 0.662734 | 14 | 7 | 0.883999 | False |
| vermis_shape_MeshVolume | 70–79 | 70–79 | -0.023027 | 0.974903 | 29 | 25 | 0.988901 | False |
| | 80–89 | 80–89 | 1.196626 | 0.228777 | 11 | 7 | 0.640877 | False |
| | 60–69 | 60–69 | -0.422789 | 0.659934 | 14 | 7 | 0.883999 | False |
| vermis_shape_VoxelVolume | 70–79 | 70–79 | -0.012305 | 0.988901 | 29 | 25 | 0.988901 | False |
| | 80–89 | 80–89 | 1.200132 | 0.232177 | 11 | 7 | 0.640877 | False |
| | 60–69 | 60–69 | -2.742208 | 0.037396 | 14 | 7 | 0.560944 | False |
| vermis_firstorder_Minimum | 70–79 | 70–79 | -2.659003 | 0.010999 | 29 | 25 | 0.329967 | False |
| | 80–89 | 80–89 | -1.030074 | 0.310169 | 11 | 7 | 0.640877 | False |

##### *6.2.6 Putamen:*

Table LII. Female subgroup t-test results for putamen radiomic features by age bin.

| Feature | Normal bin | AD bin | $t$-stat | $p$-value | $n$ normal | $n$ AD | FDR $p$ | Significant FDR |
|---|---|---|---|---|---|---|---|---|
| | 60–69 | 60–69 | -0.678797 | 0.465353 | 14 | 7 | 0.634573 | False |
| putamen_firstorder_90Percentile | 70–79 | 70–79 | -1.102160 | 0.266973 | 29 | 25 | 0.446955 | False |
| | 80–89 | 80–89 | 0.012878 | 0.989901 | 11 | 7 | 0.989901 | False |
| | 60–69 | 60–69 | -1.003905 | 0.311769 | 14 | 7 | 0.491379 | False |
| putamen_firstorder_Median | 70–79 | 70–79 | -1.127090 | 0.260374 | 29 | 25 | 0.446955 | False |
| | 80–89 | 80–89 | 0.119502 | 0.915708 | 11 | 7 | 0.981116 | False |
| | 60–69 | 60–69 | 0.012675 | 0.911909 | 14 | 7 | 0.981116 | False |
| putamen_glcm_ClusterShade | 70–79 | 70–79 | 0.398624 | 0.690531 | 29 | 25 | 0.900693 | False |
| | 80–89 | 80–89 | -2.223057 | 0.026397 | 11 | 7 | 0.298770 | False |
| | 60–69 | 60–69 | -1.243173 | 0.192381 | 14 | 7 | 0.446955 | False |
| putamen_firstorder_Mean | 70–79 | 70–79 | -1.156840 | 0.259974 | 29 | 25 | 0.446955 | False |
| | 80–89 | 80–89 | 0.059718 | 0.959904 | 11 | 7 | 0.989901 | False |
| | 60–69 | 60–69 | 1.876201 | 0.095990 | 14 | 7 | 0.435290 | False |
| putamen_shape_Flatness | 70–79 | 70–79 | 1.205556 | 0.246775 | 29 | 25 | 0.446955 | False |
| | 80–89 | 80–89 | -1.213462 | 0.239976 | 11 | 7 | 0.446955 | False |
| | 60–69 | 60–69 | -1.689461 | 0.118188 | 14 | 7 | 0.435290 | False |
| putamen_firstorder_10Percentile | 70–79 | 70–79 | -1.107841 | 0.268173 | 29 | 25 | 0.446955 | False |
| | 80–89 | 80–89 | 0.170454 | 0.894511 | 11 | 7 | 0.981116 | False |
| | 60–69 | 60–69 | -0.918702 | 0.343966 | 14 | 7 | 0.491379 | False |
| putamen_shape_MinorAxisLength | 70–79 | 70–79 | -1.983475 | 0.049795 | 29 | 25 | 0.298770 | False |
| | 80–89 | 80–89 | 0.242024 | 0.794121 | 11 | 7 | 0.981116 | False |
| | 60–69 | 60–69 | 1.410472 | 0.125987 | 14 | 7 | 0.435290 | False |
| putamen_glrlm_ShortRunEmphasis | 70–79 | 70–79 | 2.081175 | 0.041196 | 29 | 25 | 0.298770 | False |
| | 80–89 | 80–89 | 1.141835 | 0.259174 | 11 | 7 | 0.446955 | False |
| | 60–69 | 60–69 | -0.892649 | 0.340766 | 14 | 7 | 0.491379 | False |
| putamen_shape_Maximum2DDiameterSlice | 70–79 | 70–79 | -2.516050 | 0.015798 | 29 | 25 | 0.298770 | False |
| | 80–89 | 80–89 | 0.167244 | 0.869313 | 11 | 7 | 0.981116 | False |
| | 60–69 | 60–69 | 1.424635 | 0.130587 | 14 | 7 | 0.435290 | False |
| putamen_glrlm_RunLengthNonUniformityNormalized | 70–79 | 70–79 | 2.066245 | 0.043196 | 29 | 25 | 0.298770 | False |
| | 80–89 | 80–89 | 1.120007 | 0.261574 | 11 | 7 | 0.446955 | False |

##### *6.2.7 Insula:*

Table LIII. Female subgroup t-test results for insula radiomic features by age bin.

| Feature | Normal bin | AD bin | $t$-stat | $p$-value | $n$ normal | $n$ AD | FDR $p$ | Significant FDR |
|---|---|---|---|---|---|---|---|---|
| | 60–69 | 60–69 | -1.395617 | 0.145985 | 14 | 7 | 0.312826 | False |
| insula_firstorder_Median | 70–79 | 70–79 | -2.144070 | 0.040796 | 29 | 25 | 0.293221 | False |
| | 80–89 | 80–89 | 0.731633 | 0.506349 | 11 | 7 | 0.660456 | False |
| | 60–69 | 60–69 | -1.319290 | 0.178782 | 14 | 7 | 0.357564 | False |
| insula_firstorder_Mean | 70–79 | 70–79 | -1.559950 | 0.124388 | 29 | 25 | 0.312826 | False |
| | 80–89 | 80–89 | 1.009547 | 0.354765 | 11 | 7 | 0.537089 | False |
| | 60–69 | 60–69 | 1.618719 | 0.105189 | 14 | 7 | 0.312826 | False |
| insula_firstorder_RootMeanSquared | 70–79 | 70–79 | 1.535801 | 0.124988 | 29 | 25 | 0.312826 | False |
| | 80–89 | 80–89 | -1.372289 | 0.207179 | 11 | 7 | 0.365610 | False |
| | 60–69 | 60–69 | -2.030574 | 0.063994 | 14 | 7 | 0.293221 | False |
| insula_firstorder_Minimum | 70–79 | 70–79 | -3.321989 | 0.002600 | 29 | 25 | 0.038996 | True |
| | 80–89 | 80–89 | -0.770027 | 0.459354 | 11 | 7 | 0.626392 | False |
| | 60–69 | 60–69 | 1.928604 | 0.073793 | 14 | 7 | 0.293221 | False |
| insula_firstorder_Energy | 70–79 | 70–79 | 1.489925 | 0.145585 | 29 | 25 | 0.312826 | False |
| | 80–89 | 80–89 | -0.534394 | 0.582142 | 11 | 7 | 0.727677 | False |
| | 60–69 | 60–69 | 1.928604 | 0.071193 | 14 | 7 | 0.293221 | False |
| insula_firstorder_TotalEnergy | 70–79 | 70–79 | 1.489924 | 0.138186 | 29 | 25 | 0.312826 | False |
| | 80–89 | 80–89 | -0.534394 | 0.611139 | 11 | 7 | 0.733367 | False |
| | 60–69 | 60–69 | -0.934750 | 0.375962 | 14 | 7 | 0.537089 | False |
| insula_firstorder_90Percentile | 70–79 | 70–79 | -0.923540 | 0.371563 | 29 | 25 | 0.537089 | False |
| | 80–89 | 80–89 | 0.479232 | 0.655334 | 11 | 7 | 0.756155 | False |
| | 60–69 | 60–69 | -0.166286 | 0.855114 | 14 | 7 | 0.950127 | False |
| insula_firstorder_Skewness | 70–79 | 70–79 | -0.098209 | 0.926707 | 29 | 25 | 0.958663 | False |
| | 80–89 | 80–89 | -2.093050 | 0.049995 | 11 | 7 | 0.293221 | False |
| | 60–69 | 60–69 | -0.943901 | 0.349165 | 14 | 7 | 0.537089 | False |
| insula_firstorder_10Percentile | 70–79 | 70–79 | 0.022056 | 0.996700 | 29 | 25 | 0.996700 | False |
| | 80–89 | 80–89 | 2.018792 | 0.078192 | 11 | 7 | 0.293221 | False |
| | 60–69 | 60–69 | 1.498387 | 0.199980 | 14 | 7 | 0.365610 | False |
| insula_ngtdm_Strength | 70–79 | 70–79 | 3.227415 | 0.002000 | 29 | 25 | 0.038996 | True |
| | 80–89 | 80–89 | 0.112218 | 0.909709 | 11 | 7 | 0.958663 | False |

##### *6.2.8 Heschl's Gyrus:*

Table LIV. Female subgroup t-test results for Heschl's gyrus radiomic features by age bin.

| Feature | Normal bin | AD bin | $t$-stat | $p$-value | $n$ normal | $n$ AD | FDR $p$ | Significant FDR |
|---|---|---|---|---|---|---|---|---|
| | 60–69 | 60–69 | -0.735865 | 0.432157 | 14 | 7 | 0.499796 | False |
| heschls_gyrus_firstorder_Median | 70–79 | 70–79 | -2.031568 | 0.051595 | 29 | 25 | 0.169783 | False |
| | 80–89 | 80–89 | 1.196655 | 0.244776 | 11 | 7 | 0.386488 | False |
| | 60–69 | 60–69 | -0.988371 | 0.298370 | 14 | 7 | 0.424458 | False |
| heschls_gyrus_firstorder_Mean | 70–79 | 70–79 | -2.366509 | 0.024998 | 29 | 25 | 0.158384 | False |
| | 80–89 | 80–89 | 1.011062 | 0.336366 | 11 | 7 | 0.424458 | False |
| | 60–69 | 60–69 | -2.086714 | 0.056594 | 14 | 7 | 0.169783 | False |
| heschls_gyrus_firstorder_Minimum | 70–79 | 70–79 | -4.600507 | 0.000200 | 29 | 25 | 0.005999 | True |
| | 80–89 | 80–89 | -2.062136 | 0.048595 | 11 | 7 | 0.169783 | False |
| | 60–69 | 60–69 | 2.029731 | 0.037396 | 14 | 7 | 0.169783 | False |
| heschls_gyrus_gldm_DependenceEntropy | 70–79 | 70–79 | 1.749085 | 0.079192 | 29 | 25 | 0.210440 | False |
| | 80–89 | 80–89 | -0.406446 | 0.700530 | 11 | 7 | 0.719128 | False |
| | 60–69 | 60–69 | -0.461907 | 0.631137 | 14 | 7 | 0.676218 | False |
| heschls_gyrus_firstorder_90Percentile | 70–79 | 70–79 | -1.494388 | 0.139586 | 29 | 25 | 0.261724 | False |
| | 80–89 | 80–89 | 0.559588 | 0.581942 | 11 | 7 | 0.646602 | False |
| | 60–69 | 60–69 | 0.750795 | 0.433157 | 14 | 7 | 0.499796 | False |
| heschls_gyrus_glrlm_RunEntropy | 70–79 | 70–79 | 1.741647 | 0.088991 | 29 | 25 | 0.210440 | False |
| | 80–89 | 80–89 | 0.345698 | 0.719128 | 11 | 7 | 0.719128 | False |
| | 60–69 | 60–69 | 1.064699 | 0.329967 | 14 | 7 | 0.424458 | False |
| heschls_gyrus_firstorder_Kurtosis | 70–79 | 70–79 | 2.010785 | 0.050195 | 29 | 25 | 0.169783 | False |
| | 80–89 | 80–89 | 2.554484 | 0.026397 | 11 | 7 | 0.158384 | False |
| | 60–69 | 60–69 | -2.427442 | 0.021198 | 14 | 7 | 0.158384 | False |
| heschls_gyrus_glszm_GrayLevelNonUniformityNormalized | 70–79 | 70–79 | -3.655605 | 0.000400 | 29 | 25 | 0.005999 | True |
| | 80–89 | 80–89 | -1.657831 | 0.111789 | 11 | 7 | 0.223578 | False |
| | 60–69 | 60–69 | 1.837192 | 0.091191 | 14 | 7 | 0.210440 | False |
| heschls_gyrus_firstorder_TotalEnergy | 70–79 | 70–79 | 0.979452 | 0.339566 | 29 | 25 | 0.424458 | False |
| | 80–89 | 80–89 | -1.249800 | 0.216978 | 11 | 7 | 0.361631 | False |
| | 60–69 | 60–69 | 1.837192 | 0.101390 | 14 | 7 | 0.217264 | False |
| heschls_gyrus_firstorder_Energy | 70–79 | 70–79 | 0.979452 | 0.332367 | 29 | 25 | 0.424458 | False |
| | 80–89 | 80–89 | -1.249799 | 0.212779 | 11 | 7 | 0.361631 | False |

##### *6.2.9 Hippocampus:*

Table LV. Female subgroup t-test results for hippocampus radiomic features by age bin.

| Feature | Normal bin | AD bin | $t$-stat | $p$-value | $n$ normal | $n$ AD | FDR $p$ | Significant FDR |
|---|---|---|---|---|---|---|---|---|
| | 60–69 | 60–69 | 2.226257 | 0.036796 | 14 | 7 | 0.078392 | False |
| hippocampus_gldm_GrayLevelNonUniformity | 70–79 | 70–79 | 7.238190 | 0.000200 | 29 | 25 | 0.001200 | True |
| | 80–89 | 80–89 | 4.852834 | 0.000600 | 11 | 7 | 0.002571 | True |
| | 60–69 | 60–69 | 1.342458 | 0.167783 | 14 | 7 | 0.218848 | False |
| hippocampus_firstorder_Uniformity | 70–79 | 70–79 | 5.402764 | 0.000200 | 29 | 25 | 0.001200 | True |
| | 80–89 | 80–89 | 2.768713 | 0.020198 | 11 | 7 | 0.052995 | False |
| | 60–69 | 60–69 | 1.283467 | 0.188181 | 14 | 7 | 0.225817 | False |
| hippocampus_glcm_JointEnergy | 70–79 | 70–79 | 4.200538 | 0.000200 | 29 | 25 | 0.001200 | True |
| | 80–89 | 80–89 | 2.134957 | 0.061994 | 11 | 7 | 0.116238 | False |
| | 60–69 | 60–69 | 2.226507 | 0.033597 | 14 | 7 | 0.077531 | False |
| hippocampus_glcm_ClusterShade | 70–79 | 70–79 | 6.705670 | 0.000200 | 29 | 25 | 0.001200 | True |
| | 80–89 | 80–89 | 1.096566 | 0.256974 | 11 | 7 | 0.275330 | False |
| | 60–69 | 60–69 | 1.227874 | 0.255174 | 14 | 7 | 0.275330 | False |
| hippocampus_glrlm_RunVariance | 70–79 | 70–79 | 3.833977 | 0.000200 | 29 | 25 | 0.001200 | True |
| | 80–89 | 80–89 | 1.631169 | 0.139186 | 11 | 7 | 0.203980 | False |
| | 60–69 | 60–69 | 1.697687 | 0.066393 | 14 | 7 | 0.117165 | False |
| hippocampus_gldm_DependenceVariance | 70–79 | 70–79 | 2.124730 | 0.039196 | 29 | 25 | 0.078392 | False |
| | 80–89 | 80–89 | 1.394302 | 0.182982 | 11 | 7 | 0.225817 | False |
| | 60–69 | 60–69 | 1.085732 | 0.253975 | 14 | 7 | 0.275330 | False |
| hippocampus_glcm_MaximumProbability | 70–79 | 70–79 | 3.652549 | 0.001200 | 29 | 25 | 0.004500 | True |
| | 80–89 | 80–89 | 1.532739 | 0.157384 | 11 | 7 | 0.214615 | False |
| | 60–69 | 60–69 | -0.844389 | 0.393361 | 14 | 7 | 0.393361 | False |
| hippocampus_firstorder_Median | 70–79 | 70–79 | -1.712255 | 0.092791 | 29 | 25 | 0.154651 | False |
| | 80–89 | 80–89 | 1.624574 | 0.142786 | 11 | 7 | 0.203980 | False |
| | 60–69 | 60–69 | 2.518892 | 0.021198 | 14 | 7 | 0.052995 | False |
| hippocampus_firstorder_Skewness | 70–79 | 70–79 | 2.535742 | 0.015598 | 29 | 25 | 0.046795 | True |
| | 80–89 | 80–89 | -2.865663 | 0.008799 | 11 | 7 | 0.029330 | True |
| | 60–69 | 60–69 | 1.155494 | 0.275172 | 14 | 7 | 0.284661 | False |
| hippocampus_glrlm_LongRunEmphasis | 70–79 | 70–79 | 3.802897 | 0.000400 | 29 | 25 | 0.002000 | True |
| | 80–89 | 80–89 | 1.717198 | 0.115988 | 11 | 7 | 0.183140 | False |

##### 6.2.10 *Entorhinal Cortex:*

Table LVI. Female subgroup t-test results for entorhinal cortex radiomic features by age bin.

| Feature | Normal bin | AD bin | $t$-stat | $p$-value | $n$ normal | $n$ AD | FDR $p$ | Significant FDR |
|---|---|---|---|---|---|---|---|---|
| | 60–69 | 60–69 | 3.840283 | 0.003400 | 14 | 7 | 0.010799 | True |
| entorhinal_cortex_glcm_Idmn | 70–79 | 70–79 | 5.145588 | 0.000200 | 29 | 25 | 0.002000 | True |
| | 80–89 | 80–89 | 2.060799 | 0.062194 | 11 | 7 | 0.094491 | False |
| | 60–69 | 60–69 | 2.031496 | 0.067193 | 14 | 7 | 0.095990 | False |
| entorhinal_cortex_firstorder_Skewness | 70–79 | 70–79 | 1.534886 | 0.145585 | 29 | 25 | 0.181982 | False |
| | 80–89 | 80–89 | 1.119202 | 0.278372 | 11 | 7 | 0.321199 | False |
| | 60–69 | 60–69 | -3.713444 | 0.003600 | 14 | 7 | 0.010799 | True |
| entorhinal_cortex_glrlm_ShortRunLowGrayLevelEmphasis | 70–79 | 70–79 | -5.364743 | 0.000200 | 29 | 25 | 0.002000 | True |
| | 80–89 | 80–89 | -2.728065 | 0.011599 | 11 | 7 | 0.023198 | True |
| | 60–69 | 60–69 | 3.313012 | 0.005799 | 14 | 7 | 0.014499 | True |
| entorhinal_cortex_glcm_Idn | 70–79 | 70–79 | 4.356190 | 0.000600 | 29 | 25 | 0.003000 | True |
| | 80–89 | 80–89 | 0.841043 | 0.404560 | 11 | 7 | 0.433457 | False |
| | 60–69 | 60–69 | -4.334108 | 0.000600 | 14 | 7 | 0.003000 | True |
| entorhinal_cortex_ngtdm_Contrast | 70–79 | 70–79 | -5.906370 | 0.000200 | 29 | 25 | 0.002000 | True |
| | 80–89 | 80–89 | -2.919143 | 0.006999 | 11 | 7 | 0.016152 | True |
| | 60–69 | 60–69 | 3.416679 | 0.008999 | 14 | 7 | 0.019284 | True |
| entorhinal_cortex_glszm_HighGrayLevelZoneEmphasis | 70–79 | 70–79 | 3.307912 | 0.002400 | 29 | 25 | 0.008999 | True |
| | 80–89 | 80–89 | -0.605438 | 0.545145 | 11 | 7 | 0.563944 | False |
| | 60–69 | 60–69 | 2.693499 | 0.025597 | 14 | 7 | 0.047995 | True |
| entorhinal_cortex_firstorder_Range | 70–79 | 70–79 | 1.836573 | 0.078992 | 29 | 25 | 0.107717 | False |
| | 80–89 | 80–89 | 1.826584 | 0.114989 | 11 | 7 | 0.149985 | False |
| | 60–69 | 60–69 | 2.688936 | 0.033397 | 14 | 7 | 0.055661 | False |
| entorhinal_cortex_glrlm_HighGrayLevelRunEmphasis | 70–79 | 70–79 | 3.443484 | 0.002200 | 29 | 25 | 0.008999 | True |
| | 80–89 | 80–89 | -0.436346 | 0.669733 | 11 | 7 | 0.669733 | False |
| | 60–69 | 60–69 | -3.476219 | 0.005000 | 14 | 7 | 0.013635 | True |
| entorhinal_cortex_glszm_LowGrayLevelZoneEmphasis | 70–79 | 70–79 | -4.358130 | 0.000400 | 29 | 25 | 0.003000 | True |
| | 80–89 | 80–89 | -0.911488 | 0.361364 | 11 | 7 | 0.401515 | False |
| | 60–69 | 60–69 | 2.168098 | 0.062994 | 14 | 7 | 0.094491 | False |
| entorhinal_cortex_ngtdm_Strength | 70–79 | 70–79 | 1.415607 | 0.168383 | 29 | 25 | 0.202060 | False |
| | 80–89 | 80–89 | -2.508918 | 0.028797 | 11 | 7 | 0.050818 | False |
